# When can whole-genome SNP heritability be reliably estimated from summary statistics?

**DOI:** 10.64898/2026.05.13.724972

**Authors:** Benjamin Pham, Samuel Davenport, David Azriel, Armin Schwartzman

## Abstract

LD Score Regression (LDSC) is a prominent method, which estimates whole-genome SNP heritability from summary statistics via the slope of a linear regression of GWAS test statistics corresponding to a trait of interest against LD scores. It was claimed by the LDSC authors that the free intercept in the regression accounts for confounding bias such as population stratification. In this study, we argue that the intercept in LDSC must be fixed to 1 for accurate SNP heritability estimation. We show both theoretically and with simulations that the estimated intercept does not accurately capture population stratification effects, and that it adversely affects the accuracy of the heritability estimate introducing bias and increasing variance. Fixing the intercept to 1 eliminates bias and reduces variance when no population stratification is present. On the other hand, under population stratification, LDSC is biased with both the free and the fixed intercept. Additionally, we show that estimated standard errors in LDSC are underestimated, potentially leading to false-positives in downstream GWAS analyses.

## 1 Introduction

Genetic variations are responsible for the manifestation of traits. Almost all human genomic DNA is identical (about 99% of genomic DNA sequence) and only a small amount of differences are responsible for phenotype variation in a population [2]. Single nucleotide polymorphisms (SNPs) are the simplest form of genetic variation in which a single base-pair in the genome is changed. To date, the 1000 Genomes Project has identified 81 million SNPs in humans [1]. It is of great interest to identify SNPs that influence trait variations in a population. Genome-wide Association Studies (GWAS) simultaneously test the association between SNPs and observed traits.

Population heritability of a trait is not easily calculated, since gathering information directly from an entire population is impossible. Instead, a sample is collected and assumed to be representative of the population, and the sample heritability is used as an estimate of the population heritability. If individual-level data are readily available, that is, genotype data and the observed phenotype measure in the sample, then the SNP heritability can be estimated by genetic restricted maximum likelihood (GREML) using the Genomic Complex Trait Analysis (GCTA) package [39]. However, individual-level data can be computationally expensive to handle as the sample size becomes large and it also constitutes sensitive information pertaining to human subjects. As such, GWAS practitioners instead release statistical test associations between observed phenotype measures and SNPs known as *summary statistics*.

From these summary statistics, heritability can still be estimated with some degree of accuracy compared to using the raw data, but it is not as reliable [22, 26]. The differences between individual-level methods and summary statistics methods arise because individual-level methods are able to adjust for confounding effects within the model [28, 40] while summary statistics methods are less flexible. One such example of confounding is population stratification of subjects which occurs when the population being sampled is a heterogeneous mixture of two or more subpopulations. This adds an unobserved effect to the phenotype that affects the accuracy of the heritability estimation.

The most popular whole-genome method to estimate SNP heritability from summary statistics is Linkage Disequilibrium Score Regression (LDSC) [7]. LDSC estimates SNP heritability as the slope of a weighted regression of phenotype-SNP association test statistics on sum of squared correlations between SNPs (called *LD scores*). Despite the variety of existing methods, LDSC continues to be the most popular method of SNP heritability estimation because of its accessibility and fast computation time. Our work focuses on LDSC due to its popularity in the field with more than 6000 citations as of March 2026.

We also consider GWAS Heritability (GWASH) [29], a method-of-moments whole-genome heritability estimator that uses summary statistics, based on the individual-level data Dicker estimator [12]. When the assumptions for the Dicker estimator hold, GWASH is roughly equivalent to LDSC with the intercept fixed at 1 [29, 4]. The main practical advantage of using GWASH is that it offers an estimate of the SE of the estimator in a closed form that is easy to implement and understand [29].

In [4] it is shown that both GWASH and LDSC are consistent under two important conditions. First, the correlation between the SNPs must be local. Second, all SNPs should have effects that are about the same order of magnitude. The first condition is violated under population stratification (see Section 7.1 in [4]). The second condition is violated if a sizable subset of SNPs have effects that are much larger than the rest. These issues, especially the violation of the first condition, are central to the results obtained in this paper.

In [7], it was claimed that the intercept in LDSC regression distinguishes confounding from polygenicity in GWAS as it measures the contribution of confounding biases, such as cryptic relatedness and population stratification. However, they caution (as has been noticed by others e.g. [22]) that whenever confounding effects are correlated with LD, LDSC estimates heritability poorly. Others have also found that estimating the intercept in an attempt to separate confounding from genetic signal could lead to erroneous heritability estimation [11, 33]. Further, it is stated in the discussion section of [7] that it is preferable to correct for confounding biases directly because adjusting test statistics is no substitute for diligent quality control. Other critics support this approach [11, 22].

The issue of constraining the intercept or not has somewhat been discussed by the LDSC development team on their GitHub Frequently Asked Questions page [14], see Table 1. It is unclear when fixing the intercept is appropriate as the LDSC SNP heritability of most studies are reported with the fitted intercept rather than estimating the heritability with a fixed intercept [38, 35], presumably because fitting a free intercept is the default option in the LDSC software. We will make the argument that the intercept should typically be fixed.

**Table 1:** Questions about when to constrain (fix) the LDSC intercept and the LDSC authors’ guidance [7]. The direct quotations are from [14] while unquoted text is our interpretation of the quotation.

| Question | Answer | Comment |
| --- | --- | --- |
| “When should I constrain the intercept?” | “If you can rule out QC problems, such as inflation from population stratification. For genetic correlation, you should constrain the intercept if you can quantify sample overlap (or you are sure that there is no sample overlap).” | Here the authors state that the intercept should be constrained (fixed) if one can rule out QC (quality control) problems such as inflation from population stratification. |
| “Why is the standard error for my $h^2$ estimate so high?” | “The common culprits are small sample size, small number of regression SNPs, too many partitioned LD scores. Try constraining the regression intercept.” | Here the authors mention constraining (fixing) the intercept as a way to fix the variance issue, but it is unclear when this is appropriate. |

The aim of this work is to identify conditions under which LDSC yields accurate and robust results. This is important because inaccurate heritability estimates can undermine our understanding of the genetic architecture of complex traits, as reliable partitioned heritability estimates require a trustworthy overall heritability benchmark [15, 3]. Furthermore, poor heritability estimates can negatively impact downstream analyses, such as predictive gene expression in transcriptome-wide association studies [19] and genomic structural equation modeling [18]. By clarifying these conditions, we aim to improve the accuracy of heritability estimation and, in turn, enhance the reliability of subsequent genetic analyses.

Here, we establish and recommend that the intercept should be fixed to 1 when estimating heritability with LDSC. We demonstrate both theoretically and with simulations that the free-intercept LDSC estimator is biased in the presence of population stratification. Thus, the claim in [7] that the least squares estimator with free intercept corrects for confounding coming from population stratification is incorrect. Our conclusion is that in the presence of population stratification the estimators are biased and should not be used, and if there is no population stratification, the intercept should be fixed to 1.

We consider the behavior of heritability estimators via simulations with a reference panel that matches the distribution of the individual-level data in two settings: a controlled theoretical setting where SNPs are correlated according to an AR1 process [29] and a realistic setting where SNPs are correlated with LD structure coming from data. The AR1 theoretical setting can be used to understand the precise effect that correlation has on the estimates. Previous works show simulations that are only derived from real genetic LD structure [7, 26, 33, 42, 30]. We demonstrate, with both reference-panel data and individual-level data, that the LDSC estimator is highly variable in non-population stratified regimes while LDSC with fixed intercept and GWASH can successfully estimate heritability in these settings. We also show that in the presence of large confounding effects, such as high population stratification and high genetic stratification, all heritability estimation methods are highly biased.

We also evaluate standard errors computed from the closed-form formula defined in [29] for GWASH and those supplied by block-jackknife in LDSC [7] in both AR1 simulations and realistic simulations. We find that both of these methods underestimate the empirical standard errors in most AR1 simulations and all realistic simulations. This is important because unreliable estimation of variance negatively impacts downstream analyses, especially evaluating the statistical power and reliability of the heritability estimate based on its corresponding z-score.

Finally, we estimate SNP heritability in 89 real GWAS datasets using both GWASH and LDSC, with and without fixing the intercept, to assess how well the behaviors observed in simulations manifest in real data. We show that fixing the intercept results in somewhat higher SNP heritability estimates than with free intercept. GWASH provides similar estimates to LDSC with the fixed intercept. We also computed the z-score of the heritability estimate of each dataset and discuss the reliability of those z-scores.

## 2 Methods

### 2.1 Population Model

We consider a model of observed phenotypes from genotypes for *n* subjects and *m* SNPs, including population stratification effects, as proposed in the original LDSC paper [7, 8]:

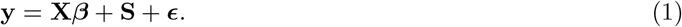

Here **y** is an *n* × 1 vector of phenotype measurements with entries *y*_*i*_, **X** is a *n* × *m* genotype matrix with entries *X*_*ij*_, ***β*** is an *m* × 1 vector of SNP effects with entries *β*_*j*_, **S** is the *n* × 1 population stratification term due to the environment with entries *S*_*i*_, and ***ϵ*** is an *n* × 1 vector of environmental effects not explainable by SNPs with entries *ϵ*_*i*_; *i*∈ {1, …, *n*} and *j* ∈ {1, …, *m*}. Depending on the trait, the response **y** could be continuous or binary. For our investigation in this work, we only consider continuous traits.

We assume that the subjects are i.i.d. From Equation (1), we define SNP-based heritability as the proportion of the total phenotypic variance in *y*_*i*_ that is explained by the additive effects of the SNPs, represented by 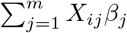:

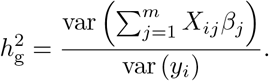

We follow the same setting as in [8] where the “distinction between normalizing and centering in the sample and population is ignored”. Thus, we model the observed genotypes and phenotypes as being normalized in the population to have mean 0 and variance 1, i.e. E (*X*_*ij*_) = 0, var (*X*_*ij*_) = 1, E (*y*_*i*_) = 0, and var (*y*_*i*_) = 1. As part of the data processing, the observations are also normalized to have mean 0 and variance 1 in the sample, but we do not make a distinction in the notation, as this makes little difference to the results when the sample size *n* is large.

We use a simplified population stratification model such that **X** consists of a mixture of two populations where the top half **X**_1_ is an 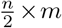 matrix containing subjects from the first subpopulation *P*_1_ and the bottom half **X**_2_ is an 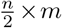 matrix containing subjects from the second subpopulation *P*_2_. To model population stratification effects from genetics, we follow [7] by using the theory in the supplementary file [8] and assume that:

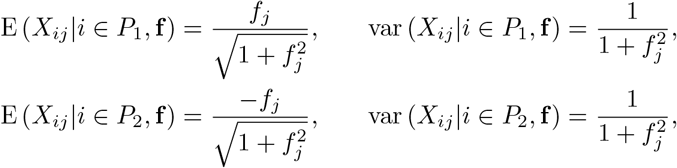

for all *j*, where **f** ~ *N* (0, F_st_**I**) is an *m* × 1 vector, **I** is an identity matrix of *m* × *m*, and F_st_ is the Wright’s constant. This construction ensures that *X*_*ij*_ has mean 0 and variance 1. We use similar F_st_ values to those defined in [8] relying on their heuristic that “F_st_ ≈ 0.1 for populations in different continents, F_st_ ≈ 0.01 for populations on the same continent, and F_st_ *<* 0.01 for subpopulations in the same country” [7].

We assume that every causal genotyped SNP in **X** contributes roughly equally to the heritability of the trait. To reflect this, the effects of the causal SNPs 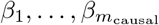 are i.i.d 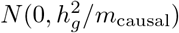, where as before 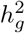 represents the heritability contribution from SNPs and *m*_causal_ is the number of SNPs, out of the total *m*, which are causal to the trait of interest while the remaining non-causal *m* − *m*_causal_ SNPs have a zero effect size.

The environmental population stratification effect of each individual in **X** dependent on population membership is represented as **S**. For each individual *i*:

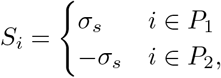

where *σ*_*s*_ is the population stratification effect dependent on population membership. In the original LDSC article [7], *S*_*i*_ is instead defined to take the values 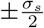. We assert that this is an error because it conflicts with the form of the error variance (see Section S1.3.2 in the Supplementary Material).

The effect from the environment, defined previously as ***ϵ***, has E [*ϵ*_*i*_] = 0 and 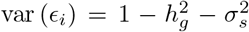 to ensure that var(*y*_*i*_) = 1 for *i* ∈ {1 … *n*}. The variance of *ϵ*_*i*_ is the fraction of variance that cannot be explained by the SNPs.

### 2.2 Data Structure and Processing

#### 2.2.1 Individual-level Data

Individual-level data are typically stored as a *n* × *m* matrix of minor allele counts per *j*th SNP of each *i*th row, where *i* = 1, …, *n* and *j* = 1, …, *m*. As such, the genotype matrix consists of the values 0, 1, or 2. The phenotype data are recorded per individual along with identifying information such as family id or sex. Direct computation of quantities such as the covariance matrix can be computationally difficult because *n* and *m* can be very large. For example, the UK Biobank has around *n* ≈ 500, 000 participants with *m* ≈ 800, 000 SNPs (which increases to *m* ≈ 96, 000, 000 SNPs if imputed SNPs are included) for thousands of phenotypes [9].

#### 2.2.2 LD Scores

The sample-statistics based methods that we will explore in the following sections rely on the LD scores - which are aggregate correlation measures. To define them formally let

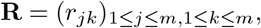

be the SNP pairwise correlation matrix, where *r*_*jk*_ is the Pearson correlation between the genotype data observed at *j*th and *k*th SNP. The population LD Score of the *j*th SNP is derived from **R** as the sum:

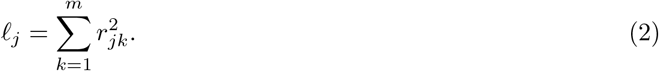

We construct the sample correlation matrix from the observed sample data as:

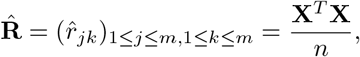

where 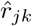 is the Pearson correlation between the *j*th and *k*th SNP in the sample, which is an estimate of the population version *r*_*jk*_. As shown in the equation, the LD matrix can only be computed with individual-level data. As in Equation (2), the sample LD scores are computed in a similar way with the observed sample data:

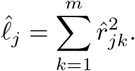

In practice, the sample LD scores are adjusted to account for “upward bias of approximately 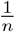” in each 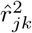 of 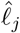 compared to the unobserved *ℓ*_*j*_ [7, 8]. We denote this adjustment with a tilde:

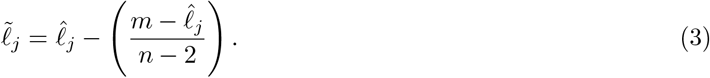

Technical details about the adjusted LD scores are discussed further in Section S1.2.2.

#### 2.2.3 Summary Statistics

We must additionally reduce the full simulation data to summary statistic quantities used in heritability estimation methods - in order to match what is done in practice. For the *j*th SNP, there is a corresponding least-squares *estimated effect size* 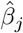 defined as

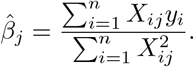

Because of standardization in the sample, the denominator is equal to *n*, yielding the simplified definition

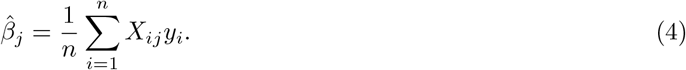

The test statistics used for LDSC are defined as follows:

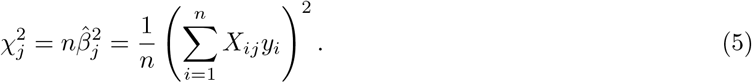

LD scores are computed directly from **X** using Equation (2) and adjusted using Equation (3).

In some datasets, instead of reporting 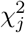 as defined in Equation (5), 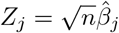 is reported. Out of the 89 datasets that we examine in Section 4, 22 datasets only contain *Z*_*j*_, while the remaining 67 datasets contain both *Z*_*j*_s and 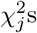. Notice that 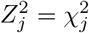, so the *χ*^2^ statistics can still be calculated from the *Z*-statistics.

In the literature, there has been a lack of a standard processing pipeline for summary statistics; the GWAS catalog reported in 2020 that ~7% of 27, 485 GWAS summary statistics considered did not report 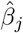 or their estimated standard errors [20]. Only very recently have there been efforts to address such inconsistencies [20, 10].

#### 2.2.4 Simulation Data

Our simulated data is generated according to the model defined in Section 2.1. We assume that each generated sample is independently drawn from the population distribution. The nominal heritability of the simulated phenotype explained by SNPs 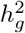 is set to 0.2 as in [29]. We prepare simulated genotype data **X** as a *n* × *m* multivariate normal matrix with mean 0 and variance 1 in the population, then standardize it so that each column of **X** has mean 0 and variance 1 in the sample. The correlation between columns follows either an AR1 process or a realistic LD structure from Chromosome 22 1000 Genomes European reference panel. For the effects we generate a *m* × 1 vector 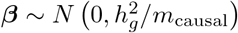. Unless stated otherwise, we assume that all *m* SNPs are causal. In cases where *m*_causal_ *< m*, we take the following steps:

1. Set prop 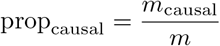 as the expected proportion of causal SNPs.
2. Generate an indicator vector of length *m* as a binomial random variable with probability *p* = prop_causal_ to determine whether each SNP is causal.
3. Set the effect sizes of non-causal SNPs to zero by multiplying ***β*** element-wise with the resulting indicator vector.

We generate the non-genetic environmental effects *n* × 1 term ***ϵ*** from a normal distribution of mean 0 and variance 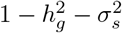. We use these quantities to compute **y** from Equation (1) and standardize it so that it has mean 0 and variance 1 in the sample.

#### 2.2.5 Reference Panel Data

In practice, without individual-level data, key quantities that depend on **X** such as 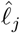 cannot be computed. Instead, a reference panel is used to construct such quantities, with the assumption that it has a population structure similar to that of the sample. LD scores constructed from a reference panel only consider SNPs in-common between **X** and the reference panel. These LD scores can be computed similarly using Equation (2) and adjusted via a modified version of Equation (3):

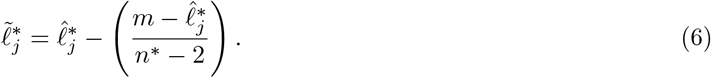

where 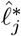 is the unadjusted LD Score from the reference panel and *n*^*\**^ is the sample size of the reference panel genotypes. For simplicity in this study, all simulations have *n*^*\**^ = *n*. However, in real data *n*^*\**^ is often much smaller than *n*.

Two additional quantities 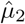 and 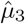 used for GWASH and defined in Section 2.3.3 below must also be computed from a reference panel because they require the trace of the squared LD matrix and the trace of the cubed LD matrix.

To analyze the 89 real summary statistic datasets in Section 4, we computed the necessary quantities for LDSC and GWASH from 1000 Genomes Phase 1 [1] with European ancestry, the same ancestry as the studies. The current release of LDSC has pre-computed adjusted LD scores from this panel [7]. Because the original files used to directly create the released LD scores were not available, we processed 1000 Genomes Phase 1 data to construct LD scores as described in [7].

### 2.3 SNP Heritability Estimation

As explained in Section 1, we consider two representative heritability estimation methods that use GWAS summary statistics: LD Score Regression (LDSC) and GWASH. We compare these summary statistics-based methods with state-of-the-art heritability estimation based on the full data known as Genome-based restricted maximum likelihood (GREML) implemented via GCTA [39].

#### 2.3.1 LD Score Regression (LDSC)

LDSC is the weighted regression of squared trait association statistics 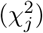 against the adjusted LD scores 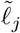. If individual level data are not available, adjusted LD scores from the reference panel are used as discussed in Section 2.2.5.

LDSC is a weighted regression that comprises of two steps. The first step is an initial baseline estimate of the intercept utilizing common SNPs in the dataset with 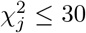 (representing likely non-causal SNPs), LD scores, and an optional set of inverse LD scores used as initial regression weights [6]. It is recommended that the regression SNPs used for weighting be the same as the input LD scores “to save computation time since LDSC is not very sensitive to the precise choice of LD scores [regression SNPs] used for” weights [14]. The heritability and intercept estimates are then used to iteratively refine weights (two iterations by default) [6]. The second step then runs an iteratively reweighted least-squares regression over all SNPs, using the intercept and refined weights from the first step as initial values, to produce the final heritability estimate [6]. LDSC outputs the intercept estimated in the first step and the heritability estimate estimated in the second step. If there are no SNPs with 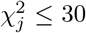, then the heritability estimate and intercept are estimated jointly based on all of the SNPs. Details on the precise algorithm can be found in Section S2.1.2 in the Supplementary Material. We will argue using our simulations that the truncation in the first step produces a bias in the heritability estimates.

Two-hundred blocks of consecutive SNPs with their corresponding 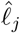 and 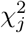 are allocated. Standard errors of heritability and intercept estimates are calculated via pseudovalues, estimated by leave-one-out block jackknife. The sample covariance matrix of the pseudovalues are computed and the square root of its diagonal entries are the reported jackknife standard errors. Details about the block jackknife procedure in LDSC can be found in Section S2.1.2 in the Supplementary Material.

#### 2.3.2 LDSC is biased under population stratification

The formula that motivates the LDSC estimators is given in [7] and in more detail in Proposition 2 of [8]. They argue that under Model (1), 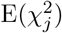 is a linear function of *ℓ*_*j*_ with slope 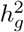. This derivation relies on the unconditional expectation. However, as we argue below, and in more detail in Section S1.3 of the Supplementary material, one should in fact consider the conditional expectation given **X**, rather than the unconditional expectation. Specifically, we show that under Model (1),

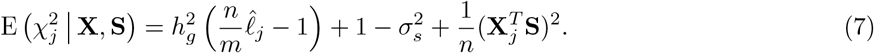

The key point is that the intercept term, 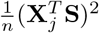, is correlated with 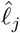 and as such when the 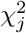’s are regressed on 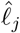’s, bias is introduced as seen in Sections 3.1 and 3.2. (see Sections S1.3.4 and S1.3.5 in the Supplementary Material for technical details).

In homogeneous populations where there is no population stratification, i.e. such that F_st_ = 0, the term 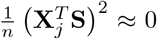 because **X** and **S** are uncorrelated. If there are also no environmental stratification effects such that *σ*_*s*_ = 0, then **S** ≡ 0 and Equation (7) becomes:

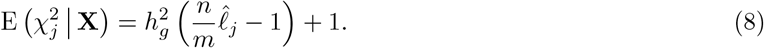

Equation (7) is different from corresponding equations presented in [7]. We prove its validity and correct mistakes in the derivation of [7] in Section S1.3.2 and Section S1.3.4 of the Supplementary Material. This includes a demonstration that the derivation of [7] for the fixed intercept estimator is circular (see Section S1.2.2 in the Supplementary Material).

#### 2.3.3 GWASH

GWASH is an alternative estimator for heritability from summary statistics. In [29] it was originally presented as:

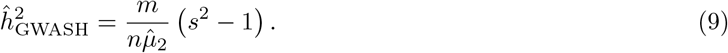

where *s*^2^ is the average of 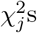 and 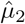 is the estimation of the LD second spectral moment:

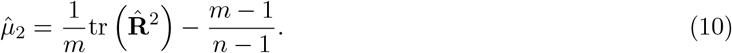

While not immediately apparent, GWASH is very close to LDSC regression with fixed intercept. Indeed it was shown in [29] and further clarified in [4] that Equation (9) can be approximately written as

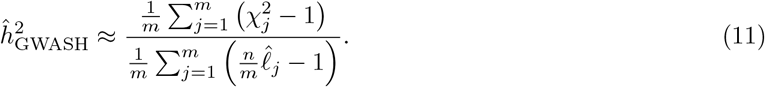

The approximation becomes equality if *m* − 1 and *n* − 1 are replaced by *m* and *n* in (10). In this form, GWASH can be seen as directly obtained from Equation (8) using the method of moments by replacing the expectation by the average and solving for 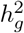.

The estimate of variance of the GWASH estimator is given by the following closed-form as presented in [29]:

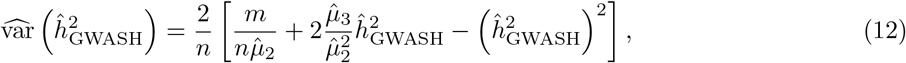

where 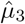 is an estimate of the LD third spectral moment and is given by,

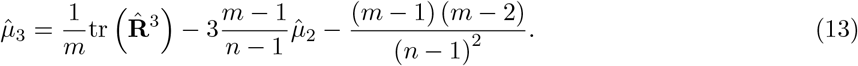

The trace of 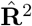 and 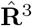, needed for 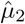 and 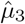, can be efficiently computed via the kinship matrix **K** (see Section S2.3.2 in the Supplementary Material for details). If full genotype and phenotype data are not available, 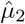 and 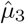 are estimated from a surrogate reference panel on common SNPs between the summary statistics and the surrogate reference panel.

The equations for LDSC and GWASH can be used as written if the SNPs in the reference panel match those of the studied dataset. In our simulations of Section 3, the reference panel and simulated datasets meet this criteria for simplicity. In real data, the SNPs in the dataset do not exactly match the SNPs in the reference panel [7]. To allow comparisons of LDSC with GWASH in Section 4, we computed the necessary quantities for Equation (12) from the 1000 Genomes Phase 1 reference panel that we had used to construct LD scores and slightly modified our implementation (see Section S2.4 in the Supplementary Material for technical details).

#### 2.3.4 GCTA

GCTA utilizes GREML (Genetic Restricted Estimated Maximum Likelihood) to estimate variance components iteratively. Although GCTA is not a summary statistics estimator—as it requires individual-level data—we include it here because summary statistics based estimators are often compared to GCTA as a performance benchmark [7, 33]. We present a simplified version of the mixed linear model presented in [39] in our notation because we do not explore non-genetic covariates. This simplified version of the equation is an alternate representation of Model (1) in variance component terms. Specifically, consider Model (1) with **S** ≡ 0 and suppose that 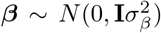. Then, like in the original equation in [39], we rewrite the model with partitioned variance components in terms of total genetic effects across *n* individuals **g** = **X***β* by substituting **XX**^**T**^ with *m***K** and 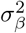 with 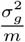. We then take

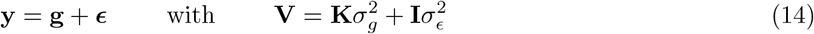

where:

- **g** is an *n* × 1 vector of total genetic effects of individuals with normal distribution 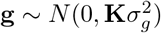.
- ***ϵ*** is an *n* × 1 vector of residuals with normal distribution 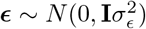.
- **K** represents the genetic relatedness of all subjects computed from all genotyped SNPs. In [39], it is called the genetic relatedness matrix (GRM). This is also known as the *n* × *n* kinship matrix and can be computed as 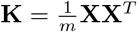.
- **V** is the *n* × *n* variance-covariance matrix of the observed phenotype of all subjects.
- 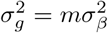 is the variance explained by all SNPs. This component is estimated iteratively.
- 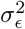 is the variance not explained by SNPs but by the environment. This component is also estimated iteratively.

GCTA can be used to estimate partitioned *h*^2^ such as in chromosomes and aggregate these partitioned heritabilities for whole genome heritability estimation. As such, there would be multiple 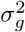 components as shown in [39]. For our purposes here, we are only looking at a simplified problem considering one group.

In brief, the GRM matrix **K** and phenotype **y** are required. An initial guess of 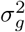 and 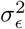 is supplied which is usually 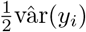 for each component. The first iteration uses Expectation-Maximization REML (EM-REML) to estimate 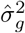 and 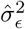 to determine the direction that these components would travel in future iterations. Subsequent iterations use the average-information algorithm [39] until convergence such that the difference between 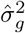 and 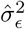 to previous iterations is less than 10^*−*4^. More technical details about the GCTA algorithm can be found in Section S2.2.1. Once the variance components are estimated, the heritability estimate from GCTA is computed as:

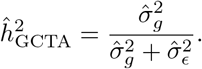

## 3 Simulations

We evaluated the performance of each summary statistic method by observing how close the heritability estimate is to 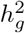 on average across 1000 simulations. We also examined the Monte Carlo (MC) standard error of each heritability estimate and the empirical standard deviation of the estimator across 1000 simulations, used throughout as a proxy for the true standard error. We present two simulation settings: one with AR1 correlation structure and another with a correlation structure derived from real genetic data. We use a sample size of *n* = 5000 because it is documented that heritability estimated from LDSC is very noisy at smaller sample sizes [14] and examine *m* = 10000 SNPs, consistent with the scale of prior heritability estimation simulation studies [29, 36, 17].

Here we illustrate the scenario where both the reference dataset and full data are drawn independently from the same distribution and the sample size of the reference dataset matches the sample size of the full data. We note that this is an ideal condition as the reference panel in real applications typically have a much smaller sample size than the study cohort. We point to Section S4 in the Supplementary Material for individual-level simulations, in which sample LD scores rather than reference LD scores are used. The LD scores in all simulations are adjusted as described in Equation (3).

### 3.1 AR1 Simulations

We identified important parameters that influence heritability estimation: SNP correlation (*ρ*), population stratification due to the environment (*σ*_*s*_), and population stratification due to genetics (F_st_). Parameters range over *ρ* ∈ {[0, 0.995]}, *σ*_*s*_ ∈ {[0, 0.8]}, and F_st_ ∈ {0, 0.05, 0.1}. We constructed 1000 simulations where **X, *β***, and **y** are generated with a set of parameters with the nominal population heritability 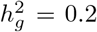. The default parameter set for AR1 simulations is {*ρ* = 0.995, *σ*_*s*_ = 0, F_st_ = 0}. We investigated the effect of parameter increases (*ρ, σ*_*s*_) in varying presence of genetic confounding by taking F_st_ ∈ {0, 0.05, 0.1}. GCTA, which requires individual-level data, serves as a benchmark for evaluating summary statistic methods.

Figure 1 demonstrates the heritability estimates and their variance as these parameters vary. Additional figures exploring the impact of prop_causal_ are shown in the Supplementary Material (see Sections S3.1 and S3.2) and illustrate that prop_causal_ has minimal impact on SNP heritability estimation because the heritability is preserved in *β*_*j*_ regardless of which SNPs are causal.

**Figure 1:**
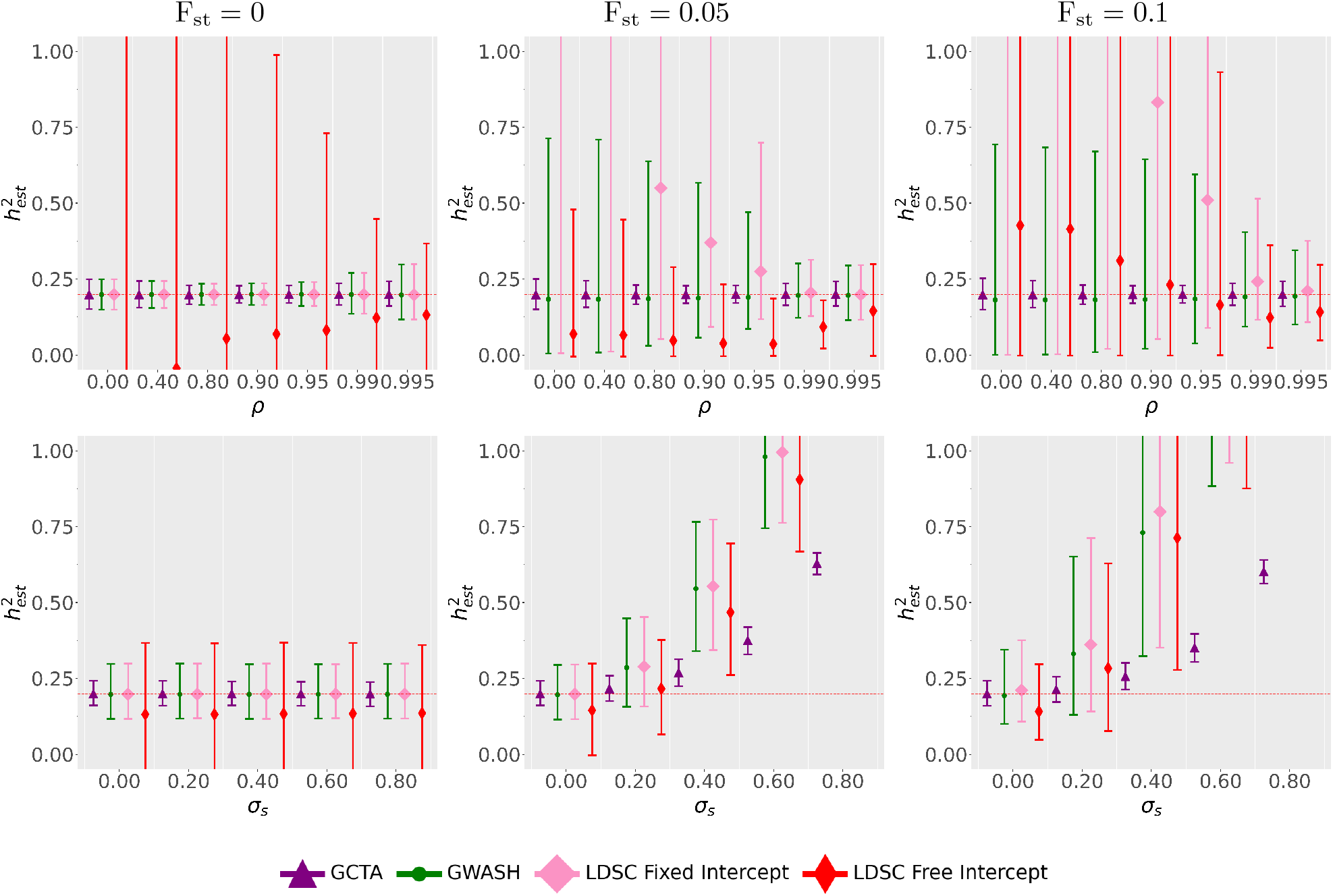
Heritability estimates when changing *ρ* and *σ*_*s*_ while gradually increasing F_st_ from F_st_ = 0 (left column) to F_st_ = 0.05 (middle column) and F_st_ = 0.1 (right column). The lower bar is the 5% quantile and the upper bar is the 95% quantile. The reference panel is computed only once from the same population as **X** and is used in all simulations. The LD scores from the reference panel are used in each heritability estimator and 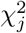 is constructed from the original data. Note that the panels only contain reasonable heritability estimates between 0 and 1 and values outside this range are not plotted.

In the setting with no genetic confounding effects, i.e., F_st_ = 0 (left column of Figure 1), LDSC with free intercept is biased downwards and is more variable than the other estimators. Both the downwards bias and excess variability of LDSC with free intercept decreases with increasing *ρ* but is still pronounced at high *ρ* values. The other estimators considered, including LDSC with the fixed intercept, are unbiased and substantially less variable. Note that in the F_st_ = 0 setting, Figure 1 shows that varying *σ*_*s*_ does not seem to change heritability estimator behavior on average. This is likely because there are an equal number of subjects in *P*_1_ and *P*_2_ so the stratification effect in the mean of **y** cancels out (see Section S1.3.3 in the Supplementary Material).

Under genetic confounding when F_st_ = 0.05 or F_st_ = 0.1 (top middle and right panels of Figure 1), LDSC with free intercept suffers from bias at all values of *ρ* considered. For F_st_ = 0.05, LDSC with free intercept is still biased downwards as in the F_st_ = 0 case. For F_st_ = 0.1, LDSC with free intercept instead exhibits upwards bias when *ρ* ≤ 0.9 and downwards bias when *ρ* ≥ 0.95. When F_st_ = 0.05 or F_st_ = 0.1, constraining the intercept to 1 in LDSC allows for unbiased heritability estimates at high enough values of *ρ*, but the estimate is biased at low correlation values. When *σ*_*s*_ = 0, GWASH appears close to unbiased but for non-zero *σ*_*s*_ and F_st_, GWASH becomes biased.

The MC standard error of LDSC with fixed intercept is close to the MC standard error of GWASH under F_st_ = 0. When setting F_st_ = 0.05, 0.1, LDSC with fixed intercept has larger MC standard error than both GWASH and LDSC with free intercept at *ρ <* 0.99. The variability of LDSC with free intercept is comparable to LDSC with fixed intercept and GWASH when F_st_ = 0.05, 0.1 and *ρ* ≥ 0.99.

The combined presence of the two different types of population stratification, when both F_st_ and *σ*_*s*_ are large (bottom middle and bottom right panels of Figure 1), results in extremely inflated heritability estimates across all methods including GCTA. The reason for this bias for LDSC was discussed in Section 2.3.2. GCTA itself produces biased heritability estimates in these scenarios because it assumes the absence of both environmental and genetic population stratification effects [39, 25].

### 3.2 Realistic LD Structure Simulations

We repeat the simulations in the previous section with a realistic LD structure obtained from real genotype data [37]. The AR1 parameter *ρ* is not varied in these simulations because there is not equivalent parameter in LD structure. Implementation details for generating **X** with a realistic LD structure appear in Section S2.3.3 in the Supplementary Material. Our realistic simulations use the first 10000 SNPs from the chromosome 22 1000 Genomes reference panel [1]. Similar to the AR1 simulations, we benchmarked summary statistics methods against GCTA with individual-level data. We increased *σ*_*s*_ with F_st_ ∈ {0, 0.05, 0.1}.

The results are plotted in Figure 2, which demonstrates that the heritability point estimates in the realistic data setting behave similarly to the AR1 data when altering *σ*_*s*_. When F_st_ = 0 (left panel of Figure 2) LDSC with free intercept has a downwards bias, while the other estimators are unbiased. When F_st_ alone is increased to F_st_ = 0.05 or F_st_ = 0.1 at *σ*_*s*_ = 0 in the middle and right panel of Figure 2, LDSC with free intercept is still biased downwards while LDSC with fixed intercept is now biased upwards; on the other hand, GWASH is unbiased. However, when using individual-level data to estimate LD scores instead of using the reference panel, both LDSC estimates are unbiased (see Figure S11 in the Supplementary Material) indicating that the observed upwards bias of LDSC with fixed intercept is driven by the mismatch between reference panel and sample. This is discussed below in Section 5.3.

**Figure 2:**
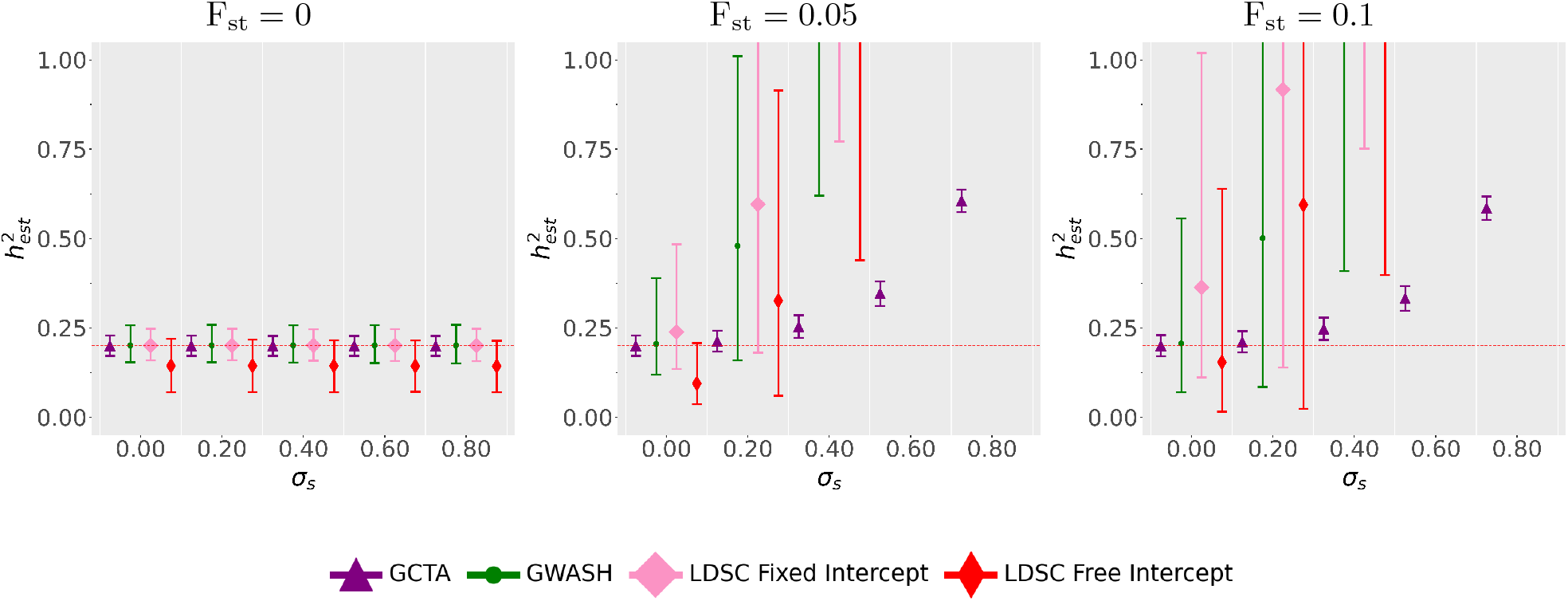
Heritability estimates on simulated data across 1000 simulations where a parameter of interest is gradually increased from F_st_ = 0 (left panel) to F_st_ = 0.05 (middle panel) and F_st_ = 0.1 (right panel). The lower bar is the 5% quantile and the upper bar is the 95% quantile. The reference panel is computed only once from the same population as **X** and is used in all simulations. The LD scores from the reference panel are used in each heritability estimator and **y** is constructed from the original data. Note that the panels only contain reasonable heritability estimates between 0 and 1 and values outside are not plotted.

Similar to the AR1 structure, LDSC with free intercept is more variable than the other estimators when F_st_ = 0. However, the magnitude of this difference is less than previously observed in AR1 simulations. All estimates are upward biased when both F_st_ and *σ*_*s*_ differ from zero. The summary statistics estimates (all estimates besides GCTA) have increased variability as F_st_ increases. Interestingly, the variance of LDSC with free intercept is less than both LDSC with fixed intercept and GWASH when F_st_ = 0.05 and *σ*_*s*_ = 0. At F_st_ = 0.1 and *σ*_*s*_ = 0, the MC standard error of LDSC with free intercept is smaller than that of LDSC with fixed intercept but is now larger than GWASH’s.

### 3.3 Estimation of standard error in AR1 Simulations

We first consider estimating standard errors in the previous settings explored in Section 3.1. We examined standard error estimates for all three heritability estimators across these 1000 simulations by comparing them with their MC standard errors, our best approximation of the true standard error, using log_10_fold change. A log_10_fold change of 0 indicates that the estimated standard error is exactly the MC standard error, negative values indicate that the estimated standard error underestimates the MC standard error, and positive values indicate that it overestimates the MC standard error.

LDSC, both fixed and free, uses a leave-one-out block jackknife to estimate standard errors (see Section 2.3.1). GWASH instead uses a closed-form estimate for the standard error (see Equation (12)). We do not consider the standard error estimate from GCTA as our focus is on summary statistics methods. As in the previous section, we examined the effect of a single parameter change on the estimated standard errors with and without genetic confounding. The goal of this analysis is to assess the accuracy of the standard error estimates. We point to Section S5.1 in the Supplementary Material for standard error estimation in individual-level simulations.

Figure 3 compares the estimated standard error for each estimator against its MC standard errors. At *ρ* ≤ 0.9 with F_st_ = 0 (top left panel), the log_10_fold change between estimated standard errors and MC standard errors of GWASH and LDSC with fixed intercept is close to zero. For *ρ >* 0.9, the standard error of GWASH is underestimated; for LDSC with fixed intercept, underestimation occurs only at *ρ* ≥ 0.99. At high *ρ*, the log_10_fold change quantile error bars of LDSC with free intercept estimated standard errors are wider than those from GWASH and LDSC with fixed intercept. When setting F_st_ = 0.05, 0.1 (top middle and right panels), all estimators except LDSC with free intercept exhibit a clear negative log_10_fold change as *ρ* increases. At *ρ* = 0.99, 0.995, the upper bound of the log_10_fold change quantile error bars for LDSC with free intercept overlaps 0.

**Figure 3:**
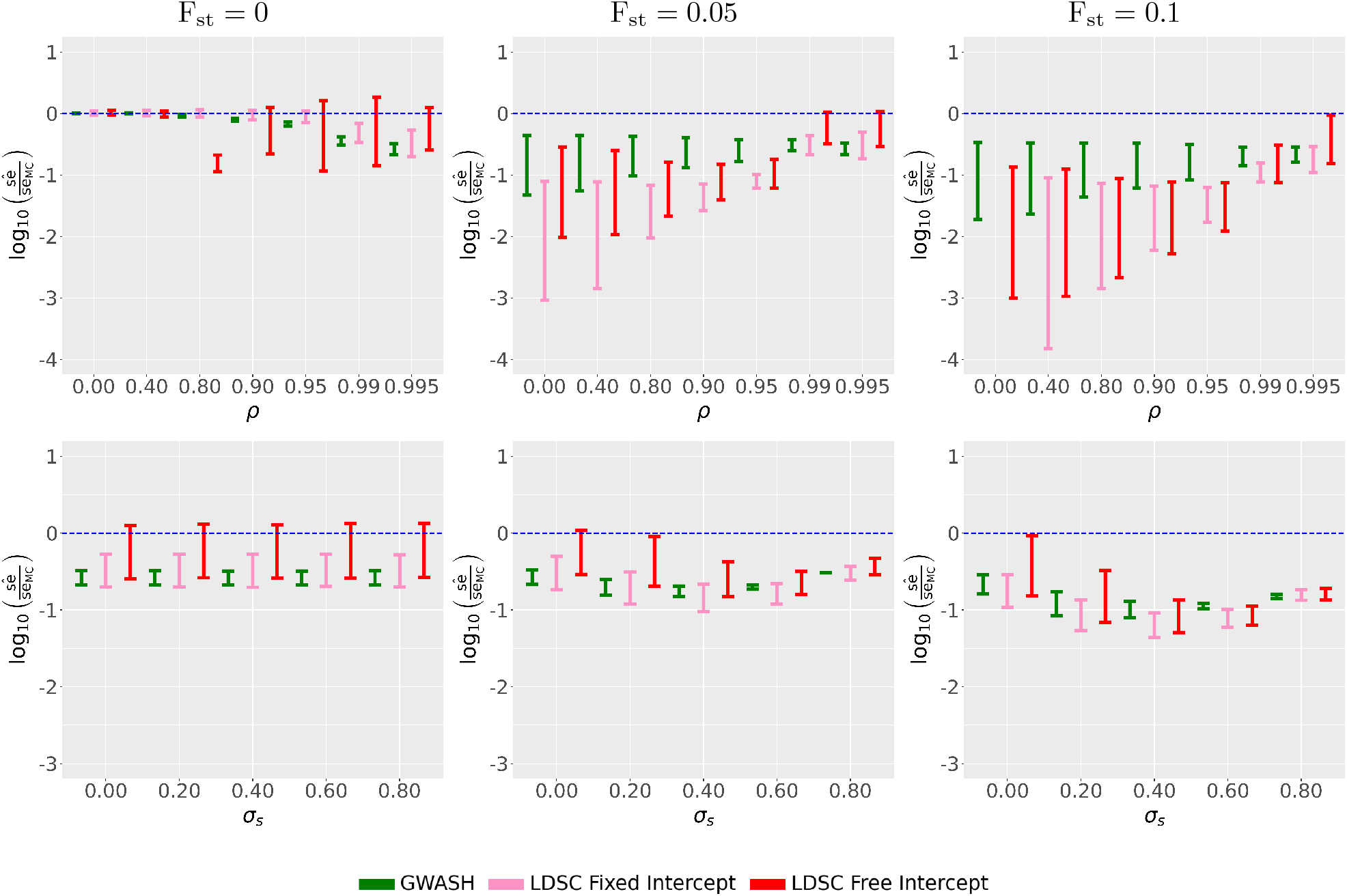
Evaluation of sample standard errors in AR1 simulated data with genetic confounding across 1000 replicates by log_10_fold change between the estimated standard error and their respective MC standard error (se_MC_) is computed. The setting is the same as in Figure 1. A log_10_fold change of 0 (blue line) indicates that 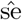 and se_MC_ are equivalent while negative values indicate underestimation and positive values indicate overestimation. The lower bar is the 5% quantile and the upper bar is the 95% quantile of the log_10_fold change. Note that the y-axis is the same across columns, but different across rows.

The bottom row of Figure 3 plots the log_10_fold change as a function of *σ*_*s*_ for F_st_ ∈ {0, 0.05, 0.1}. When F_st_ = 0 (bottom left panel), the log_10_fold change of GWASH and LDSC with fixed intercept lies below 0 and the log_10_fold change of LDSC with free intercept covers 0 for all *σ*_*s*_. For all values of F_st_ and *σ*_*s*_ considered, the GWASH fold change quantile bars are small compared to both LDSC fold change quantile bars. LDSC with free intercept has the widest error bars compared to both GWASH and LDSC with fixed intercept. At high F_st_ and *σ*_*s*_ (bottom middle and right panels), GWASH quantile error bars still remain the smallest while both LDSC quantile error bars become close. The log_10_fold changes of LDSC with free intercept becomes more negative as F_st_ increases, across all values of *σ*_*s*_.

### 3.4 Estimation of standard error in Realistic LD Structure Simulations

We also evaluated the estimated standard errors of GWASH, LDSC with fixed intercept, and LDSC with free intercept as done in the previous section under the realistic structure simulation considered in Section 3.2. We point to Section S5.2 in the Supplementary Material for standard error estimation in individual-level simulations.

The results are plotted in Figure 4. All estimators exhibit negative log_10_fold change of the estimated standard errors. At F_st_ = 0 (left panel) the error bars of the log_10_fold change are quite close across all estimators unlike in the AR1 setting. When F_st_ ∈ {0.05, 0.1} increasing *σ*_*s*_ causes the error bars to become increasingly negative until *σ*_*s*_ = 0.6. At *σ*_*s*_ ≥ 0.6, they rise slightly but still remain negative. The pattern of log_10_fold change error bar widths is similar to that observed under the AR1 setting with GWASH having the narrowest log_10_fold change error bar followed by LDSC with fixed intercept and then LDSC with free intercept.

**Figure 4:**
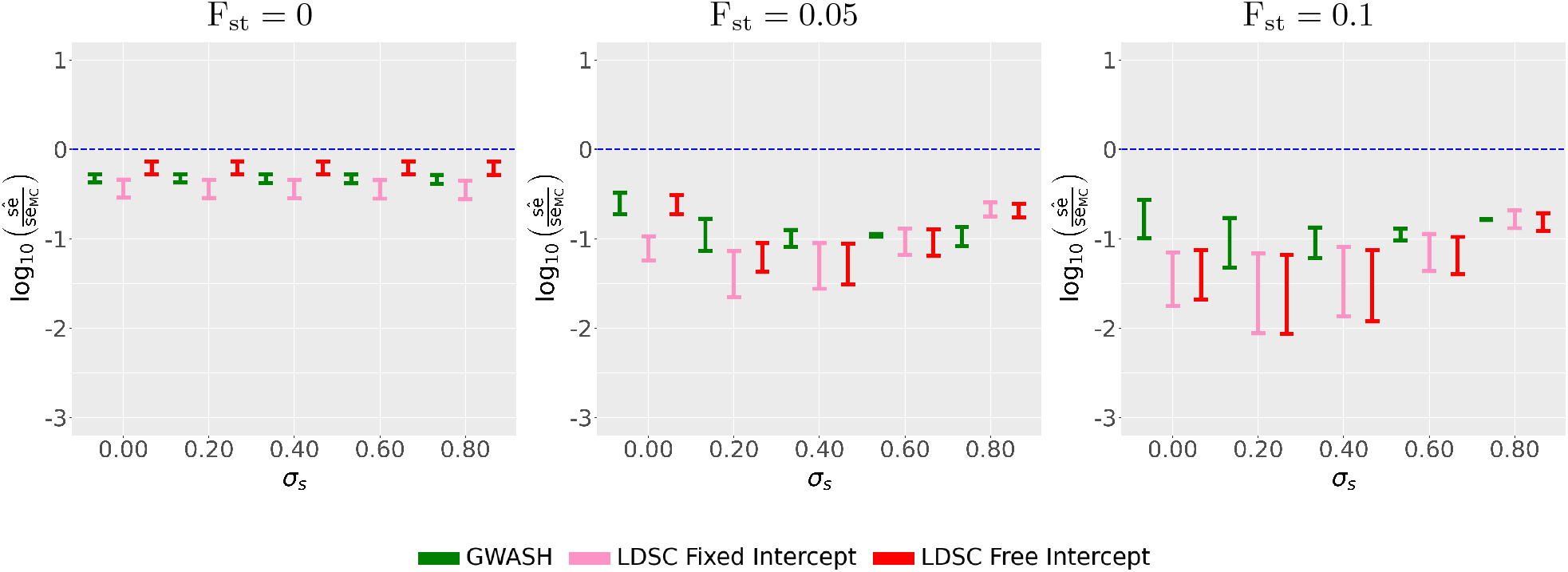
Evaluation of sample standard errors in realistic LD simulated data with genetic confounding across 1000 replicates. The setting is the same as in Figure 2. The plotting elements in each panel are the same as described in Figure 3.

### 3.5 Impact of standard error estimates on Z-Scores in AR1 Simulations

Heritability estimates and their estimated standard errors can be used in *SNP Heritability z-scores*, a metric to roughly assess the statistical power and reliability of the estimate. For a given method, the z-score is defined as:

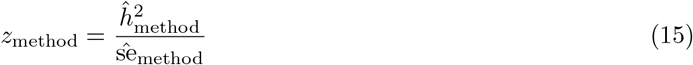

Where 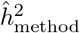 is the heritability estimate and 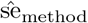 is the estimated standard error. Many papers consider a GWAS to be reliable if the heritability estimate is reasonable (i.e. 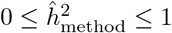) and *z*_method_ is between 4 and 7 [41, 24, 15, 21, 16, 23, 31, 3]. We consider each simulation as a “study”, and we adopt a threshold of *z*_method_ *>* 6 as a middle ground. A simulation study is “passing” for a given method if these two criteria are met: 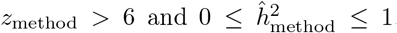. We gauge the impact of heritability estimation from the studied methods by evaluating the percentage of our simulation studies that follow the above guidelines. To emphasize the impact of the underestimated standard errors that we have observed in Section 3.3 and Section 3.4, we compute *z*_method_ with both the estimated standard error (transparent bar) and the MC standard error (solid bar) in the figures below. We first start with the AR1 simulations in Figure 5.

**Figure 5:**
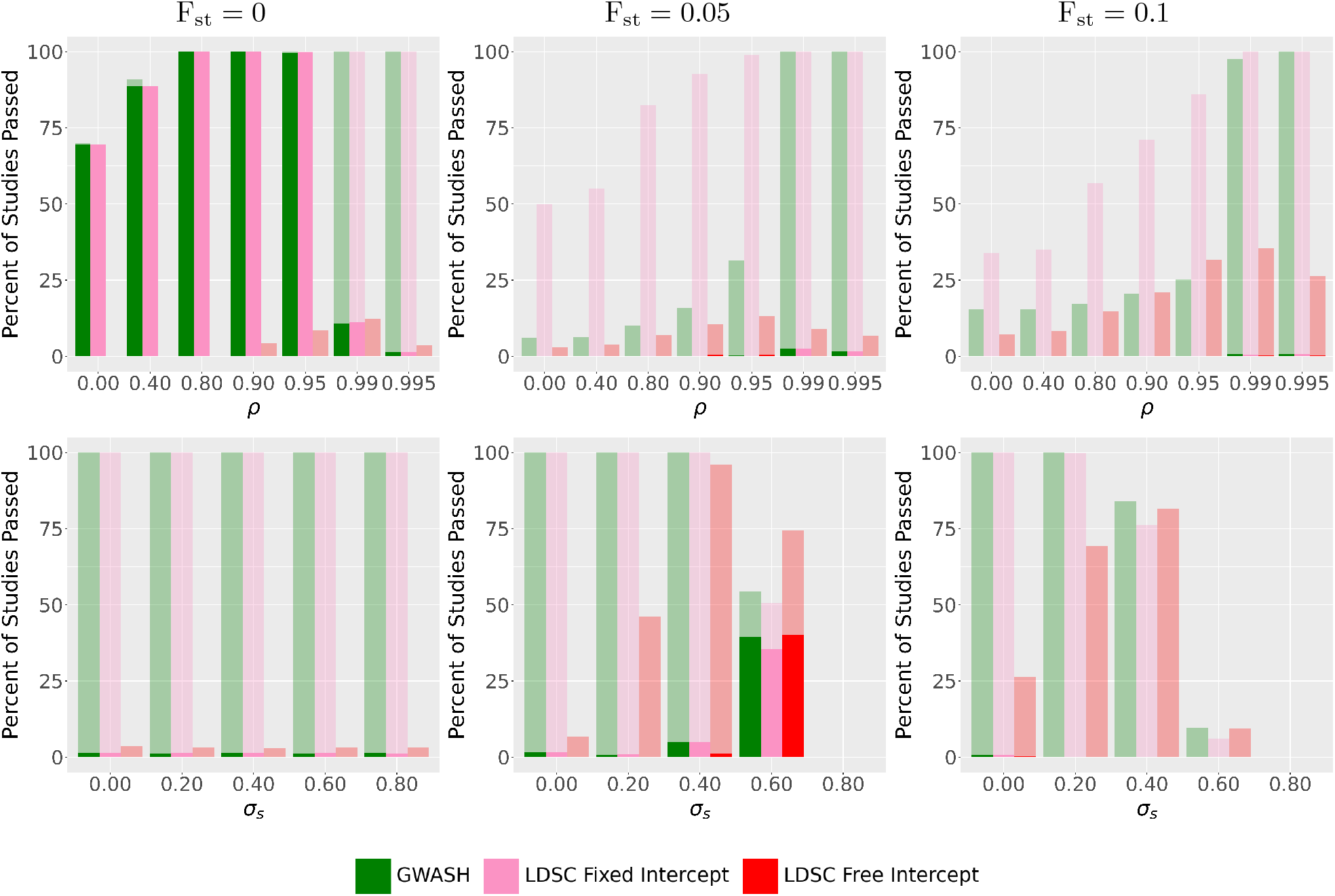
Evaluation of studies passed based on heritability z-scores in AR1 Simulations changing *ρ* or *σ*_*s*_ after F_st_ is increased. The left column shows no genetic confounding (F_st_ = 0.0), the middle column is with moderate genetic confounding (F_st_ = 0.05) and the right column is with high genetic confounding (F_st_ = 0.1). The transparent bars show the number of studies passed with a heritability z-score *>* 6 with the estimated standard errors while the solid bars show the number of studies passed with a heritability z-score *>* 6 with the MC standard errors.

We observe in Section 3.1 that, when there is no genetic confounding, the heritability estimates for LDSC with fixed intercept and GWASH seem unbiased with small simulation variance while LDSC with the free intercept is biased downwards with a highly variable heritability estimate. We also observe in Section 3.3 that the estimated standard errors underestimate the MC standard errors. Z-scores computed with the (larger) MC standard errors are smaller which heavily reduces the number of studies passed for all heritability estimators. All of these factors result in a much lower percentage of passed studies for LDSC with free intercept than the other heritability estimators. This observation is more apparent when F_st_ = 0 and *ρ* increases, as shown in the top left panel of Figure 5. In contrast to LDSC with free intercept, many studies passed under LDSC with fixed intercept and GWASH estimators. In lower correlation regimes (*ρ* ≤ 0.4), approximately 70 − 80% studies pass while all studies passed when *ρ* is increased higher (0.8 ≤ *ρ* ≤ 0.95) even with the MC standard error. At the highest possible correlation regimes (*ρ* ≥ 0.99), all studies pass with the estimated standard errors, yet fewer than ~ 10% of studies pass with the MC standard errors.

The percentage of studies passed is static across all *σ*_*s*_ when F_st_ = 0 (left side of bottom left panel in Figure 5) because *σ*_*s*_ does not greatly influence heritability estimates, see Section 3.1. Only ~ 4% of studies passed for LDSC with free intercept under the estimated standard error while no studies pass under the MC standard error. All studies pass for GWASH and LDSC with fixed intercept under the estimated standard errors. However, when using MC standard errors, only ~ 1% studies pass for each estimator.

Under genetic confounding (F_st_ = 0.05 and F_st_ = 0.1) while *σ*_*s*_ = 0(left side of bottom middle and bottom right panels in Figure 5), the percentage of studies passed for LDSC with free intercept under the estimated standard error increases from ~ 7% with F_st_ = 0.05 to ~ 26% with F_st_ = 0.1. This is because of the improved regression as previously discussed in Section 3.1. In contrast, all studies for GWASH and LDSC with fixed intercept pass under the estimated standard error in both F_st_ = 0.05 and F_st_ = 0.1. We note that little to no studies pass for all three heritability estimators when using the MC standard error.

Under genetic confounding with environmental confounding (*σ*_*s*_ ≠ 0 in the bottom middle and bottom right panel in Figure 5), there is a trend that more studies pass in LDSC with free intercept until reaching a peak around a particular *σ*_*s*_ value. After this point, the percentage of studies passed for all three estimators gradually decreases to 0 with larger *σ*_*s*_. This peak occurs with F_st_ = 0.05 around *σ*_*s*_ = 0.4 when using the estimated standard error, and it occurs around *σ*_*s*_ = 0.6 when using the MC standard error. This occurs because, around these *σ*_*s*_ values, some heritability estimates (bottom middle panel in Figure 1), although still biased upwards, start going above 1 and thus fail to satisfy the reasonable heritability estimate condition. Additionally, some MC standard errors around these *σ*_*s*_ values (bottom middle panel of Figure 3) are small enough to yield large z-scores which makes the studies look well-powered. This is most prominently seen at *σ*_*s*_ = 0.6. There are no studies passed at *σ*_*s*_ = 0.8 with F_st_ = 0.05 (right of bottom middle panel in Figure 1) because all heritability estimates are above 1.

In contrast, with F_st_ = 0.1, the peak occurs around *σ*_*s*_ = 0.2 only when using the estimated standard error. It does not appear when using the MC standard error for each estimator because the error is not small enough (bottom right panel of Figure 3) to push the z-scores above the *z*_method_ *>* 6 threshold. Fewer heritability estimates (bottom right panel of Figure 1) are below 1 at *σ*_*s*_ = 0.6 with F_st_ = 0.1 which results in fewer studies passing in this regime than at *σ*_*s*_ = 0.6 with F_st_ = 0.05. As when F_st_ = 0.05, there are no studies which pass at *σ*_*s*_ = 0.8 with F_st_ = 0.1 (right of bottom right panel in Figure 1) because all heritability estimates there are above 1.

### 3.6 Impact of standard error estimates on Z-scores in Realistic LD Simulations

We repeat the analysis performed in Section 3.5, for the realistic LD simulations when F_st_ = 0, and display the results in the leftmost panel of Figure 6. As with the AR1 simulations, the estimated standard errors inflate the heritability z-scores which causes all studies to pass for GWASH and LDSC with fixed intercept. There is also a noticeable increase in the percentage of studies passing for LDSC with free intercept in the realistic setting compared to the AR1 setting with the estimated standard errors: ~24% of studies pass in the realistic setting versus ~4% in the AR1 setting (left panel of Figure 6). When using MC standard errors, more studies for GWASH and LDSC with fixed intercept pass in the realistic setting than in the AR1 setting: ~ 50% of studies pass for GWASH and ~90% of studies pass for LDSC with fixed intercept in the realistic setting compared to ~1% for both estimators in the AR1 setting. For LDSC with free intercept, using MC standard errors in the realistic setting yields ~0.5% studies passing, an increase from none in the AR1 setting.

**Figure 6:**
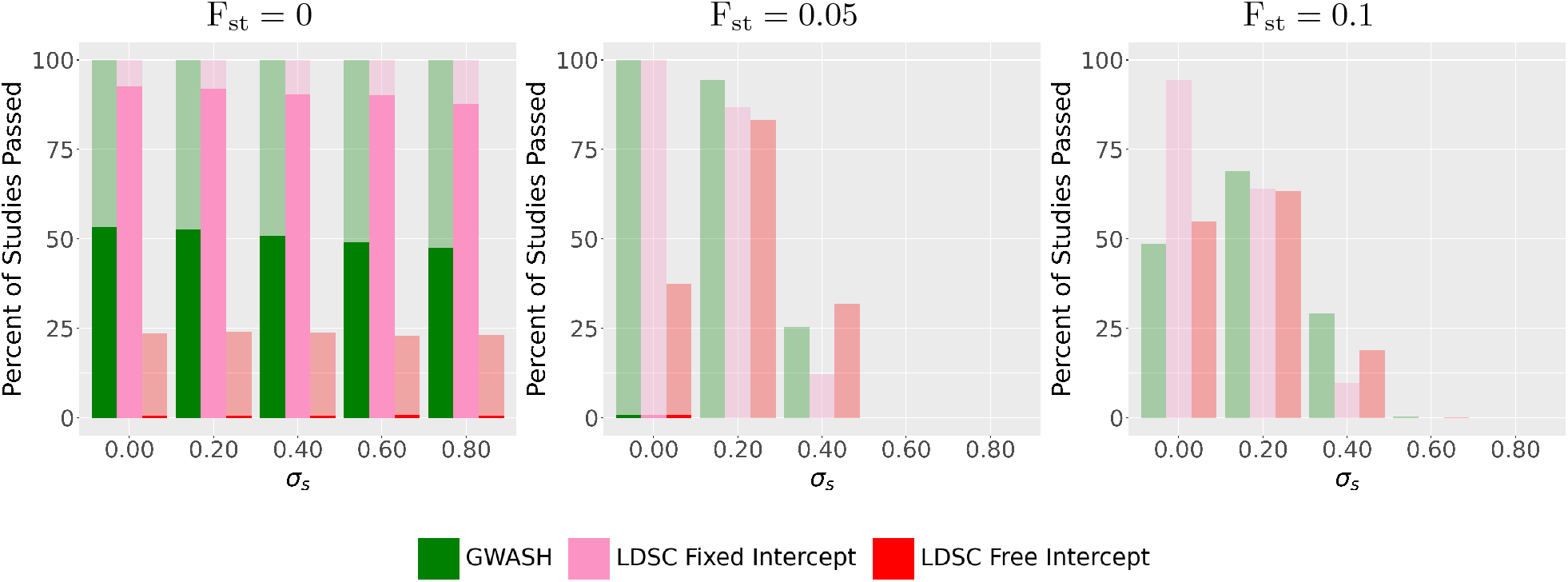
Evaluation of studies passed based on heritability z-scores in Realistic Simulations changing a parameter after F_st_ is increased from F_st_ = 0 (left panel) to F_st_ = 0.05 (middle panel) and F_st_ = 0.1 (right panel). This is the same setting as described in Figures 2 and 4. The plotting elements are the same as in Figure 5.

The trend of studies passing for the estimators in the realistic setting under genetic confounding at *σ*_*s*_ = 0 is similar to what was observed from AR1 simulations in Figure 5. Like in the AR1 setting, all studies pass for GWASH and LDSC with fixed intercept with F_st_ = 0.05 when using estimated standard errors. There is also an increase for LDSC with free intercept of percentage of studies passed in the realistic setting: from ~40% with F_st_ = 0.05 to ~55% with F_st_ = 0.1. This percent increase when F_st_ changes is close to what was observed in the AR1 setting. However, a stark difference in the realistic setting compared to the AR1 setting is that, at *σ*_*s*_ = 0, the percentage of passing studies decreases with F_st_ = 0.1 for GWASH and LDSC with fixed intercept: from 100% for both estimators to ~50% for GWASH and ~95% for LDSC with fixed intercept. This is due to the high variability of the heritability estimates (right panel in Figure 2) together with their smaller estimated standard errors (right panel in Figure 4), which yields fewer heritability z-scores that exceed the *z*_method_ *>* 6 threshold.

In the realistic setting under genetic confounding at *σ*_*s*_ ≠ 0, the downwards trend of passed studies for all estimators is more pronounced than in the AR1 setting. The downwards peak of the percentage of passed studies for all estimators when using the estimated standard error starts at *σ*_*s*_ = 0.2 for both F_st_ = 0.05 and F_st_ = 0.1 (middle and right panel in Figure 2). At increasing *σ*_*s*_, the percentage of passed studies for all estimators decrases to 0. This is because most heritability estimates are inflated over 1 at *σ*_*s*_ *>* 0.2 in the realistic setting (bottom right panel in Figure 2) whereas most heritability estimates are inflated over 1 at *σ*_*s*_ *>* 0.4 in the AR1 setting (bottom middle and bottom right panel in Figure 1). As a result, substantially fewer studies pass the *z*_method_ *>* 6 threshold in the realistic setting than in the AR1 setting at *σ*_*s*_ ≥ 0.4 under either standard error.

## 4 Real Data Application

### 4.1 Implementation details

We compared heritability estimates from three summary-statistic based methods: GWASH, LDSC with fixed intercept, and LDSC with free intercept, in 89 GWAS publicly available studies. The summary statistics datasets for these studies are available at https://github.com/TiffanyAmariuta/TCSC/tree/main/sumstats.

To compute GWASH for real summary statistic datasets, we multiplied 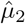 and 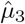 by a factor of (*n* −1)*/*(*n* − 2) to carry out the same bias correction as in 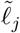. This term has a noticeable effect for smaller *n* such as that of the 1000 Genomes Phase 1 Reference Panel (see Section S2.4.2 in the Supplementary Material). We also replaced *m* − 1 with *m* in 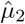 and 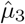 for Equation (9) (See Section S2.4.2 in the Supplementary Material). We computed 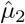 and 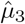 with a 1 centimorgan window of SNPs similar to LD Score computation in [7] (see Section S2.4.4 in the Supplementary Material for technical details) to directly compare to LDSC. We verify that our constructed LD scores are very close to those released by the LDSC team (see Section S2.1.3 in the Supplementary Material).

### 4.2 Heritability Estimation in Real Data

We begin by estimating heritability and corresponding standard errors using GWASH, LDSC with fixed intercept, and LDSC with free intercept across all 89 datasets. Because we do not have individual-level data available for all datasets, we cannot compare against GCTA as we did in the simulations. In Figure 7 we compare the estimators pairwise: LDSC with fixed intercept against GWASH (left panel) and against LDSC with free intercept (right panel). We omit GWASH against LDSC with free intercept because simulations (and prior work [29]) show GWASH and LDSC with fixed intercept yield very close results and hence it is enough to compare to one of them.

**Figure 7:**
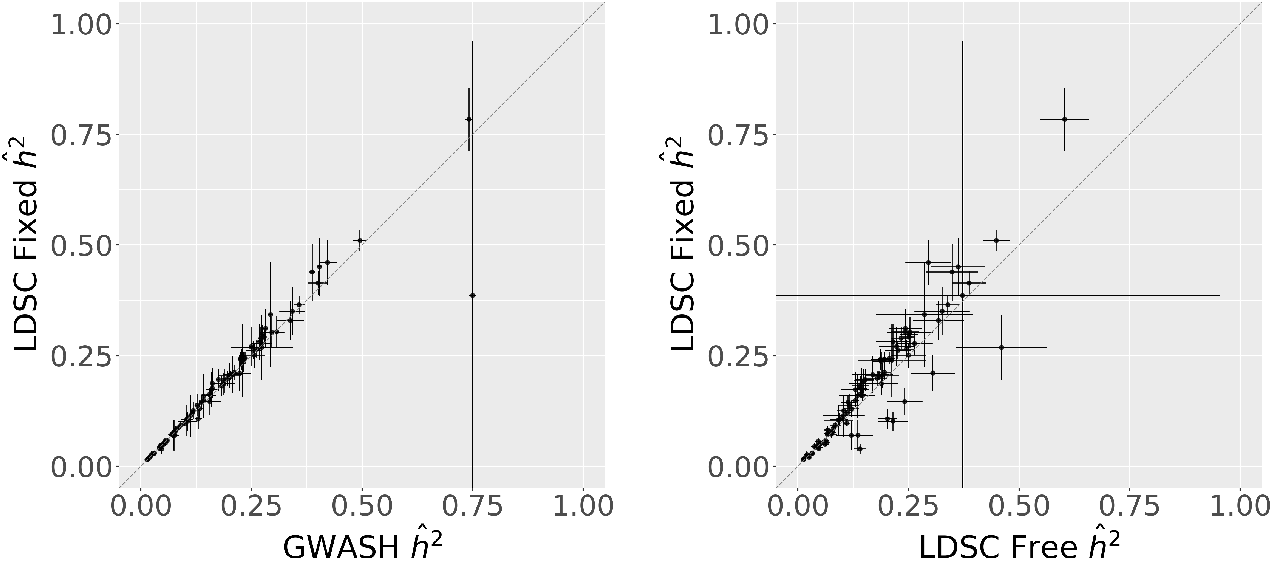
Comparison of *ĥ*^2^ and 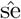 in 89 summary statistics. Each point represents a dataset. The error bars are 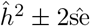. The grey dashed line has intercept 0 and slope 1, so that datasets on this line have the same value of *ĥ*^2^ in both methods. In the left panel, 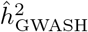 is comparable to 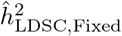 and the estimated standard error from GWASH is smaller than LDSC with fixed intercept across most datasets. In the right panel, most datasets exhibit 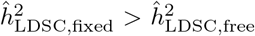 and the estimated standard error in LDSC with fixed intercept is smaller than LDSC with free intercept across most datasets.

The left panel of Figure 7 shows that heritability estimates from GWASH and LDSC with fixed intercept are mostly aligned in the 89 datasets, which is consistent with our realistic LD simulations. As also seen in our realistic simulations with genetic confounding, there are some datasets that have a higher heritability estimate from LDSC with fixed intercept than GWASH (see Figure 2).

In one dataset (Total Bilirubin from UK Biobank 460k), the heritability estimate from LDSC with fixed intercept is much smaller than that from GWASH. It was reported that the heritability of this trait comes from a small region of SNPs [32]. Because this violates the LDSC model that the polygenetic effects are evenly spread in SNPs across the whole genome [34, 7], LDSC downweights these SNPs in its regression, and it fails to fully capture the genetic signal. On the other hand, GWASH does not perform downweighting and can better capture the heritability across the genome.

In contrast to what we observe in our realistic LD simulations without genetic confounding (see left panel of Figure 4), the estimated standard errors from GWASH are smaller than those from LDSC with fixed intercept. Since we demonstrated that the estimated standard errors of all estimators underestimate the true standard error (see Figure 3 and Figure 4), we emphasize that these estimated standard errors in the real datasets are likely not reliable and cannot make a conclusion on performance.

The right panel of Figure 7 shows that the majority of heritability estimates with fixed intercept are higher than with free intercept. This is consistent with Figure 2, which shows that the free intercept causes a downward bias. The discrepancy between the two estimates also gets larger as the heritability itself increases. This is confirmed in realistic simulations with varying 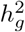 (see Figure A6 in the Appendix).

The estimated standard errors in the right panel of Figure 7 are generally larger for the free intercept estimator, which is consistent with what we observed in the realistic LD simulations without genetic confounding (see left panel of Figure 4). Some datasets exhibit similar behavior found in realistic LD simulations under genetic confounding where the estimated standard errors of LDSC with fixed intercept are larger than those of LDSC with free intercept (see middle and right panel of Figure 4).

### 4.3 Z-score differences across SNP Heritability methods in real data

We evaluated, for all of the 89 datasets, the z-scores, as defined in Equation 15. The results are plotted in Figure 8. The z-scores are far under the diagonal in the left panel of Figure 8 indicating that the z-scores of the datasets under GWASH are much larger than those of LDSC with fixed intercept. As seen in the left panel of Figure 7, the heritability estimates are similar between the two methods, so the inflation comes from the GWASH estimated standard errors being much smaller than those from LDSC with fixed intercept. This observation is consistent with our realistic simulation results in Figure 4. All 89 datasets exceed the z threshold *>* 6 under GWASH, but there are 7 datasets that fail to do so under LDSC with free intercept.

**Figure 8:**
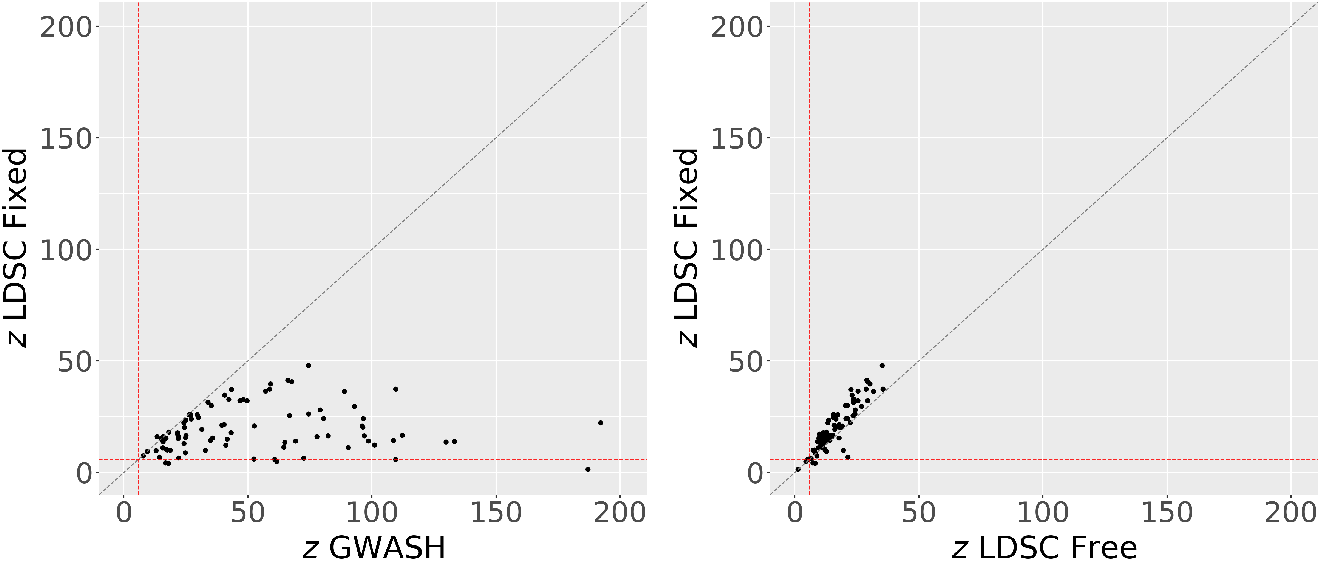
Z-scores of 89 datasets computed with *ĥ* and 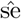 for LDSC with fixed intercept, GWASH, and LDSC with free intercept from Figure 7. The dashed red lines represent the z-score threshold of 6 to consider a dataset as passing for each method as discussed in Section 3.5. Z-scores in the upper right quadrant would represent studies that pass under both methods while z-scores in the lower-left quadrant would represent studies that would not pass in both methods. The upper-left quadrant would represent z-scores that pass in LDSC with fixed intercept only while the bottom-right quadrant would represent studies that pass under the other method (GWASH in the left panel or LDSC with free intercept in the right panel).

The z-scores lie slightly above the diagonal in the right panel of Figure 8, indicating that the z-scores of LDSC with fixed intercept are slightly higher than those from LDSC with free intercept. This is because the heritability estimates of LDSC with free intercept are slightly larger than with fixed intercept as seen in Figure A6 in the Appendix; see the discussion below in Section 5.1. The estimated standard errors are somewhat similar as shown with equidistant bars in Figure 7. While the z-scores under LDSC with free and fixed intercept are mostly similar, there are 2 datasets that have a z-score *>* 6 under LDSC with free intercept but do not under LDSC with fixed intercept. There are 5 datasets that have a z-score ≤6 under both LDSC with fixed intercept and LDSC with free intercept, and in the rest of the datasets, the z-scores are larger than 6 under both methods. Based on the simulation results in Figure 4, which show that all estimators underestimate their standard errors, we suspect that the fact that many studies appear well-powered, may be because their estimated standard errors are underestimated.

## 5 Summary and Discussion

### 5.1 The LDSC free intercept does not fix bias from population stratification and increases variance

Our results show that LDSC with free intercept as proposed in [7] is ineffective in decreasing the bias of the heritability estimate in scenarios with population stratification. Moreover, in scenarios without population stratification, using the free intercept needlessly introduces a downward bias and results in higher variance of the heritability estimate, when compared to LDSC with fixed intercept and GWASH. This downwards bias is also present across different heritability values 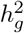 in realistic simulations (see Figure A6 in the Appendix). To understand the downward bias, recall that the LDSC intercept is first estimated from a regression of SNPs with 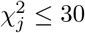. This is the default setting in LDSC from [6] where the authors assume chi-squared scores smaller than this threshold are likely to be null SNPs. This truncation, however, distorts the data distribution and results in both a higher value of the intercept and a lower value of the slope than would be otherwise obtained. We illustrate this in further detail in Section S2.1.4 of the Supplementary Material. Because the initial estimate of the intercept tends to be biased upwards, the estimate of the slope corresponding to the SNP heritability tends to be biased down in the second step after the intercept is fixed to the estimate computed in step 1.

The bias in LDSC under population stratification comes from not adequately accounting for the term 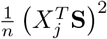 that appears in the conditional expectation of 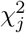 (see Equation (7)), which cannot be computed from summary statistics alone. The intercept is not a good estimator for this term, as the authors from [7] originally suggested, because fitting an intercept implies that each *j*th SNP is inflated in the same way rather than the inflation being SNP specific as it is in practice. To estimate this term, the reference panel would need to be stratified in the same way as the individual-level data which would require its corresponding environmental population stratification **S** effect to be known or approximated.

In both the AR1 setting and the realistic simulations we saw that, even under no population stratification, LDSC with free intercept has a higher variance than the other summary statistic estimators. This variance increase is more extreme in the AR1 setting, but still substantial in the realistic simulations. The variance in the AR1 setting is larger because AR1 data generation process produces a narrower distribution of LD scores than in the realistic simulations (see left column of Figure A2 in the Appendix).

Again in both settings, the variance of LDSC with free intercept is not affected by increasing *σ*_*s*_ when there is no genetic population stratification (F_st_ = 0). However, it increases substantially when both genetic and environmental population stratification effects are present (F_st_ *>* 0 or *σ*_*s*_ *>* 0) as either F_st_ or *σ*_*s*_ increase (see middle and right panel of Figure 1 and Figure 2). This is consistent with the theoretical analysis in [4].

Because the estimated intercept is typically overestimated, even when there is no population stratification, the estimated intercept can falsely indicate population stratification effects even though there is no population stratification in actuality. We thus further caution against using the LDSC intercept, as recommended by others [7, 35, 22], as a diagnostic tool for estimating the presence of population stratification in GWAS summary statistic datasets.

### 5.2 Fixing the intercept to 1 in LDSC reduces bias and variance, when there is genetic or environmental population stratification, but not both

LDSC with fixed intercept is unbiased in scenarios without population stratification. This was justified theoretically in Equation (7), which shows that, in the absence of population stratification, the intercept is indeed equal to 1.

In general, the higher variance of the heritability estimate with free intercept is the result of the regression estimating two parameters, intercept and slope, as opposed to estimating only one parameter, the slope, when the intercept is fixed. The increase in variance by unnecessarily estimating the intercept is substantial. On the other hand, constraining the intercept to 1 reduces the standard error of the heritability estimate across simulations because the regression model has one less parameter to estimate, as already observed in [5].

As we showed, LDSC with fixed intercept is still unbiased in the presence of genetic or environmental population stratification alone, but not both at the same time. When only F_st_ is increased and *σ*_*s*_ = 0, Figure 1 (top row) seems to indicate that constraining the intercept introduces upwards bias. However, this effect can be attributed to variability of the reference panel. LDSC is sensitive to mismatches between the LD scores in the data analyzed and the reference panel [7]. Even though both are constructed from the same distribution, the additional variability due to genetic population stratification introduces differences in the LD scores. In individual-level simulations (see Section S4 in the Supplementary Material), LDSC with fixed intercept heritability estimates do not exhibit upwards bias. When *σ*_*s*_ increases and F_st_ = 0, there is no mismatch in the distribution of LD scores between the reference and the original dataset and LDSC with fixed intercept performs well (see left panel of Figure 1 and Figure 2).

When both genetic and environmental population stratification are present, when either or both F_st_ and *σ*_*s*_ are increased (see middle and right panel of Figure 1 and Figure 2), all of the considered estimators are biased due to the term 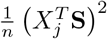 that was discussed above. Neither free nor fixed intercept solve this problem. Instead when genetic population stratification is not present LDSC with fixed intercept is unbiased and has lower variance than LDSC with free intercept, as discussed above. As such we recommend using the fixed intercept as the default approach when estimating heritability using LDSC.

### 5.3 GWASH is close to LDSC with fixed intercept but offers robustness to reference panel heterogeneity

As we have seen in Section 3, the point estimates and estimated variances of GWASH and LDSC with fixed intercept are quite similar in most scenarios. However, in the realistic simulations, LDSC with fixed intercept exhibits higher bias as F_st_ and *σ*_*s*_ increase (see Figure 2). This discrepancy is due to the mismatch between LD scores computed from the reference panel and LD scores computed from the dataset. We show in Figure S11 in the Supplementary Material that this extra bias disappears when the reference panel matches the individual-level sample data.

These results indicate that GWASH offers some robustness to reference panel heterogeneity. This can be partially explained by the fact that GWASH depends on the mean of the LD scores (see Equation (9)) - meaning that the effect of using a reference panel averages out. We further illustrate this point in the Appendix (see Figure A5), where we show that LDSC is sensitive to adding and subtracting a constant to individual LD scores, while GWASH remains the same if the average of the LD score does not change.

### 5.4 Standard errors for both LDSC and GWASH are underestimated

Standard errors for all three estimators are estimated correctly in AR1 scenarios with weak correlation (*ρ* ≤ 0.8) as shown in the top left panel of Figure 3, but heavily underestimate in regimes of stronger correlation. In the case of GWASH, this problem was not noticed in [29] because that work never considered AR1 correlations higher than *ρ* = 0.8. We argue that this is not simply a problem of estimation of the parameters in the GWASH standard error formula, but a problem with the formula itself, as it underestimates even if the true population parameters are used (see Figure S3 in Section S2.5 of the Supplementary Material).

Standard errors for all three estimators are underestimated in all other scenarios with higher correlation and/or in combination with increased F_st_ and *σ*_*s*_ (Figure 3), including in realistic LD (see Figure 4). This indicates that the block-jackknife in LDSC and the closed-form formula from GWASH are not suitable methods for standard error estimation.

It may seem surprising that the block-jackknife underestimates the error, since the classical jackknife estimator is conservative at least in the case of iid data [13]. However, SNPs are correlated and the iid assumption does not hold. The block-jackknife is an attempt to solve this issue by partitioning SNPs into blocks and then carrying out the jackknife procedure. There could be correlations between SNPs at the boundary of the blocks or long range correlations due to genetic population stratification, which again would violate the iid sampling assumption. The size of the blocks can also impact the estimate of the standard error. In our study, the small standard error could be from the small number of SNPs in each block: LDSC splits the 10000 SNPs into 200 blocks by default so each block will contain 50 SNPs. A similar result is also shown informally in [27] where a smaller block size results in smaller standard error because the impact of removing such a block is small. If a block is too large (containing many SNPs), then removing a single block would have a very large impact which results in very unstable standard error estimates.

As mentioned above, we use the same default setting of 200 blocks to reflect what is done in practice. Although it is simply a parameter to change the number of blocks, there is no guidance on choosing the optimal number of blocks. Many LDSC practitioners and tools that inherit the LDSC framework use this default setting without further exploration [6, 31, 3].

### 5.5 Inflated heritability z-scores can cause false positive GWAS results

We have shown that both LDSC (with fixed and free intercept) and GWASH can produce inflated heritability estimates, especially when using a reference panel and in the presence of population stratification (Sections 3.1 and 3.2). Moreover, their standard errors are underestimated (Section 3.4). As a consequence, heritability z-scores tend to be inflated, often highly (Section 3.5 and 3.6). Inflated heritability z-scores imply potential false positives in identifying important studies. This can result in reaching erroneous conclusions when utilizing these summary statistic studies in further downstream analyses.

As mentioned in Section 3.5, it is often recommended to trust the heritability estimates if the corresponding z-score is above a certain threshold. We found that the number of studies that pass according to this criterion is comparable across the three methods when there is a moderate amount of population stratification because the heritability estimates are inflated just enough such that the z-scores are high, while still within the reasonable heritability upper-bound of 1 (see bottom row of Figure 5). If this upper-bound criterion is loosened, as some publicly available repositories do by reporting heritability estimates that lie outside reasonable heritability estimate ranges such as in [38] or in (https://nealelab.github.io/UKBB_ldsc/index.html), then all studies would pass under population stratification because the inflated heritability estimate dominates.

In the real data analysis of 89 datasets, we observed reasonable heritability estimates. Most of these datasets have GWASH heritability estimates that are close to LDSC with fixed intercept and LDSC with fixed intercept heritability estimates are higher than LDSC with free intercept heritability estimates (see Figure 7). Because we found in our realistic simulations that standard errors are underestimated, it is reasonable to expect that underestimation of standard errors also applies to these 89 datasets.

### 5.6 Limitations of the current study

There are some limitations in our study. We model **X** as a multivariate-normal random variable for simplicity, but real genotype data consist of strictly 0, 1, 2 values. We do not expect this to have a large effect, as [29] showed that similar results are obtained under both scenarios. Our simulations use a relatively smaller number of SNPs than in practice due to computational constraints based on the **X** generating process. The LD structure is still preserved across the predictors as long as both *m* and *n* are large and the data is high dimensional (*m > n*). There exists heritability methods that are based on simulations with similar sample-size and/or predictor amounts [36, 29, 17].

While the AR1 correlation structure allows control of the strength of the correlation, it is simplistic compared to actual LD correlation patterns and does not account for allele frequencies. The realistic correlation structure instead implicitly accounts for allele frequencies and provides the closest approximation to sampling genotypes from an actual population. This ensures that our observations are from the properties of LDSC rather than from artifacts in the generated data. In this study, the correlation structure was limited to a segment of chromosome 22, which was chosen, due to being the shortest, for computational purposes. However, it may not be fully representative of the entire genome.

Previous studies fix **X** [33, 7] rather than generating **X** in each simulation. Using a fixed **X** in simulations is computationally more efficient, allowing for the direct use of real LD structure, realistic sample size, and large number of predictors. However, conclusions drawn from simulations constructed with a fixed **X** might be specific to that realization rather than to the average realizations from the population. Simulating **X** each time, as we have done in this study, captures realistic variability of sampling **X** on average from the population, but the simulation genotypes are more susceptible to randomness from the generating process.

### 5.7 Conclusions and future directions

Our findings demonstrate that the free intercept in LDSC fails to estimate confounding effects in a variety of settings and thus does not estimate heritability well. It follows that other estimators in the LDSC framework that use a free intercept [6, 31, 3] to account for confounding effects might also share this issue. A direct example is in the estimation of genetic correlations, which is a bivariate LDSC regression and uses an intercept term that attempts to indicate population stratification [6]. An extension of our work could be to investigate such methods considering a constrained intercept and study the use of the intercept term in this context.

Our work also introduced a population stratification term in the corrected LDSC equation (Equation (7)) following [4]. If this term is successfully estimated, it would allow control for stratification directly without reliance on external quality control in the sample data. In practice, estimation of this term from summary statistics alone is currently not feasible. However, it might be possible to estimate this term if partial individual-level information is available such as population membership labels. Future work could explore this possibility.

Our work also highlights the need for improved standard error estimation in both GWASH and LDSC. Works that utilize LDSC-like frameworks still use the block-jackknife procedure to estimate standard error [15, 31, 3] and therefore their results may also suffer from the same variance underestimation problem. As possible remedies, the choice of block size and number of blocks in the block-jackknife could be further explored to examine its effect on standard error estimation in different correlation structures. Variance estimation could also be approached theoretically along the lines of [4].

## 7 Data and code availability

All simulation results and code used for this study can be found at: https://github.com/b-k-pham/when_snp_h2_estimates_reliable. The 1000 Genomes Phase 1 reference panel is publicly accessible at https://www.cog-genomics.org/plink/1.9/resources. The 89 GWAS summary statistics datasets are also publicly accessible and can be retrieved from https://github.com/TiffanyAmariuta/TCSC/tree/main/sumstats/. The details of these implementations can be found in Section S2.3.2 in the Supplementary Material.

## 8 Acknowledgments

We would like to thank Tiffany Amariuta (U. of California, San Diego) for helpful discussions on LD Score Regression and critique on the draft of the paper. We would also like to thank Chun Chieh Fan (U. of California, San Diego) for his critique. This work was partially supported by NIH grant R01MH128923.

## 9 Contributions

BP: Conceptualization, Methodology, Software, Formal Analysis, Writing - Review and Editing, Writing - Original Draft

SD: Conceptualization, Methodology, Formal Analysis, Writing - Review and Editing, Supervision

DA: Conceptualization, Methodology, Formal Analysis, Writing - Review and Editing, Supervision

AS: Conceptualization, Methodology, Formal Analysis, Writing - Review and Editing, Project Administration, Funding Acquisition.

## Appendix

### A1 Effects of Parameter Changes on LDSC Regression

#### A1.1 Effect of *ρ*, F_st_, and *σ*_*s*_ on 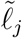 and 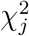

In Figure A1 we compare the 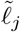’s of **X** constructed from AR1 correlation structure against 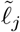’s of **X** constructed from Chromosome 22 1000 Genomes realistic LD Structure across 100 simulations. Like with the simulations in the main text, **X** has *n* = 5000 and *m* = 10000.

**Figure A1:**
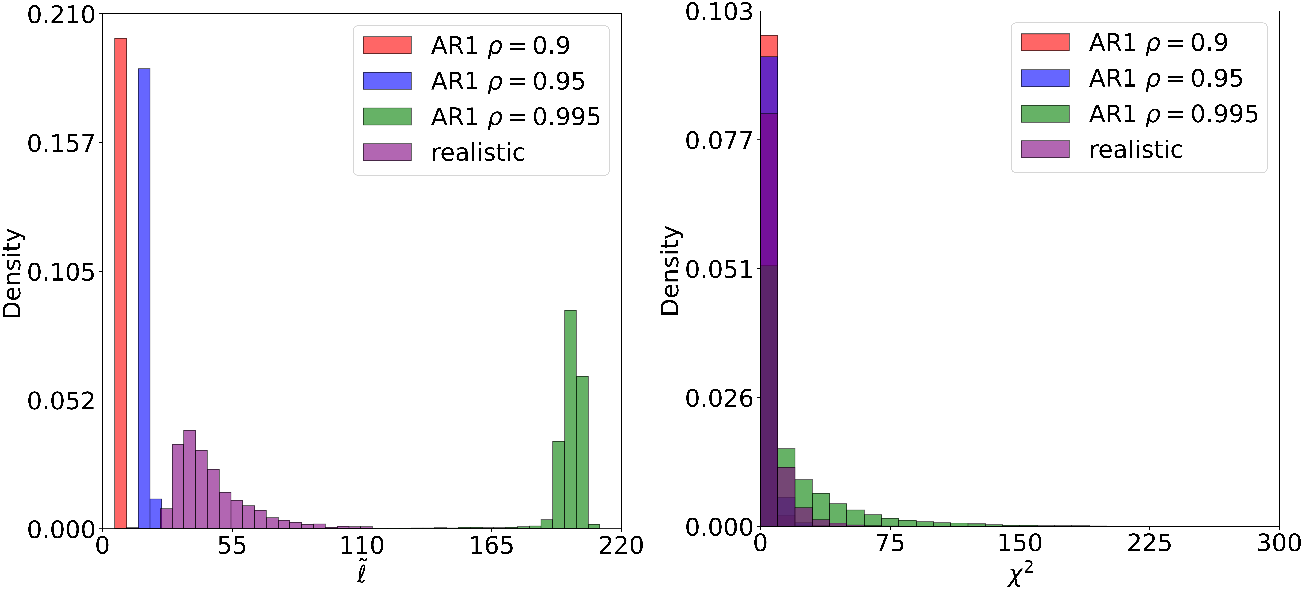
Histograms comparing 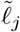(left) and 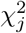(right) constructed from AR1 and realistic LD structure in **X** with F_st_ = 0 and *σ*_*s*_ = 0. Increasing *ρ* increases the magnitude of the 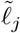’s. The 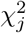’s match up across the two correlation structures in high *ρ*. The 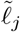’s from the realistic correlation structure have more variance than those from the AR1 correlation structure.

Heritability estimates on data with AR1 structure have wider error bars than on data with realistic LD structure (see Figure 1 for AR1 setting and Figure 2 for realistic setting) because the AR1 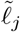’s have less variance. When predictors have low variance, regression estimates become unstable because they provide little information to explain the variation in the observed data. In LDSC, this affects stability of both the slope and intercept estimates.

**Figure A2:**
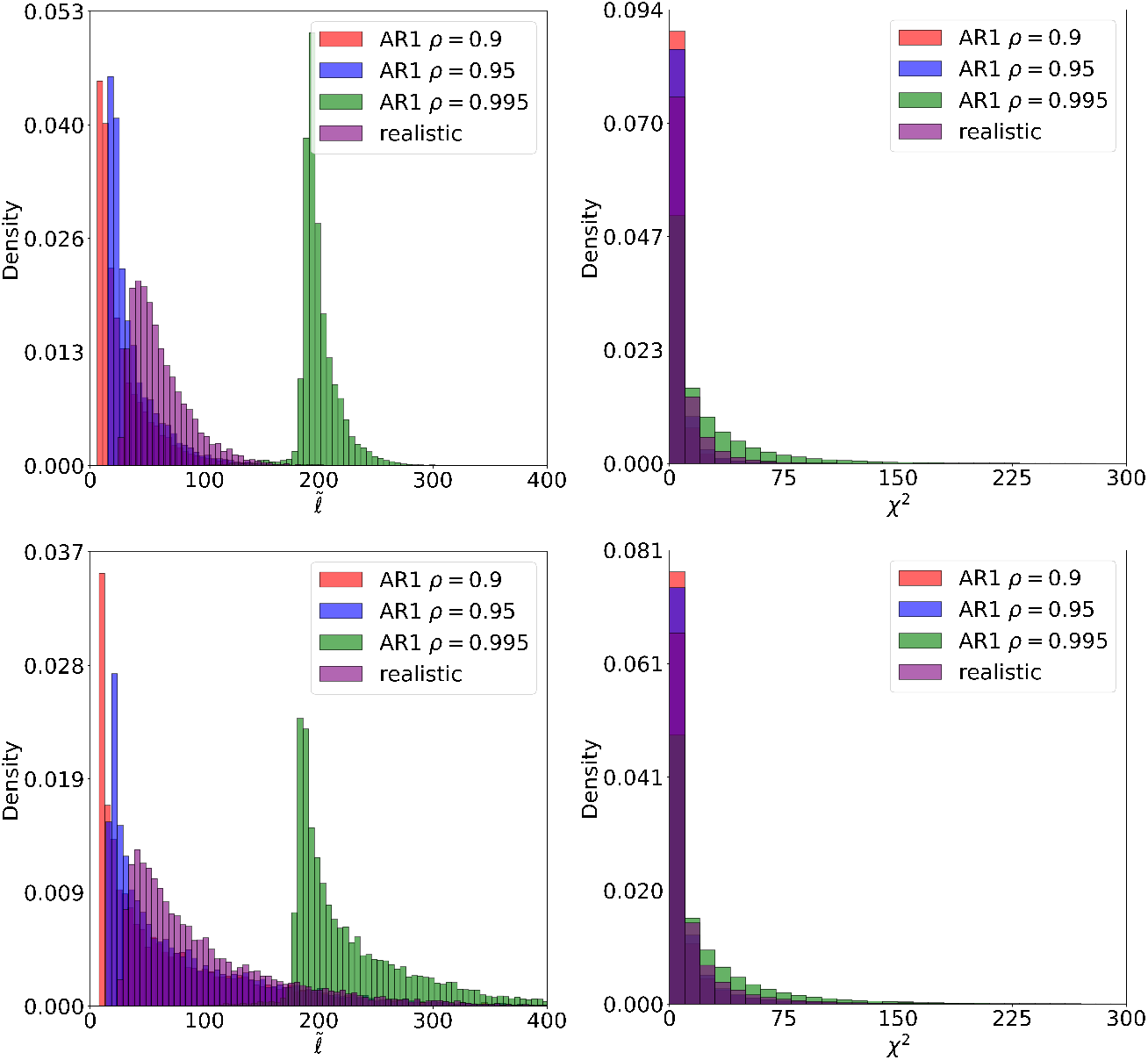
Histograms of 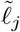 and 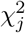 after increasing F_st_ in both non-realistic and realistic data. The top row has histograms with F_st_ = 0.05 and the bottom row has histograms with F_st_ = 0.1. Increasing F_st_ increases the variability of the 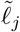’s. The 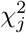’s increase a little bit, but all *χ*^2^ histograms still overlap.

We perturb F_st_ and *σ*_*s*_ in **X** in both AR1 and realistic simulations. We first change these parameters separately to see their separate effects and then together to explain why there is inaccurate heritability estimation in LDSC. Figure A2 shows that increasing F_st_ is an effective way to make the variability of AR1 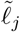’s closer to that from the realistic structure. However, they still do not align with each other.

**Figure A3:**
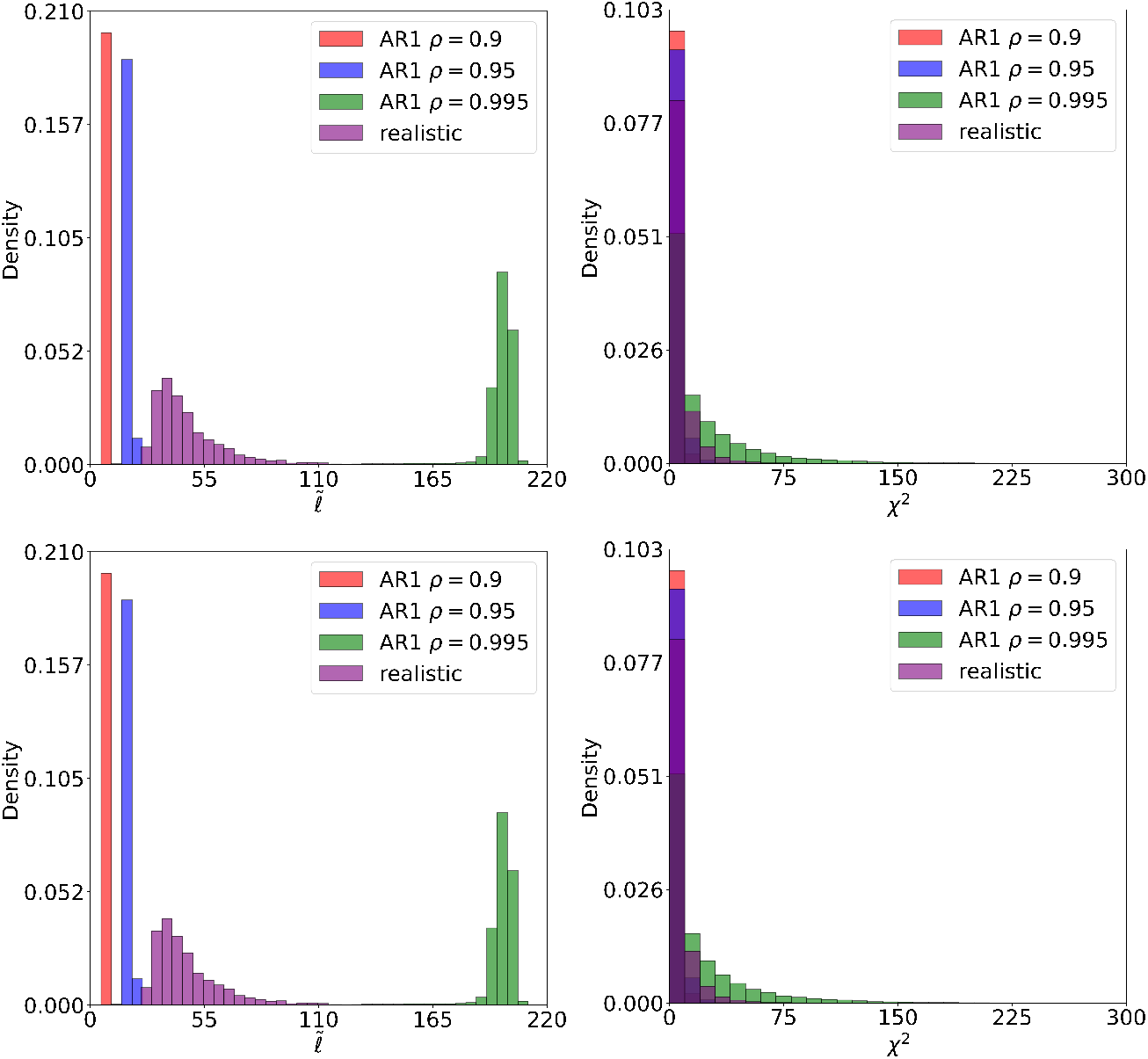
Histograms of 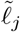 and 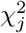 after increasing *σ*_*s*_ in both non-realistic and realistic data. The top row represents *σ*_*s*_ = 0.4 and the bottom row represents *σ*_*s*_ = 0.8. The 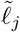 remain unchanged. The 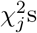 change a little bit as seen by the movement of the x-axis, but all *χ*^2^ histograms still overlap.

Figure A3 shows that increasing *σ*_*s*_ only affects 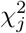’s since changing *σ*_*s*_ only affects *y* computation. LDSC can still estimate heritability reasonably well with just increased *σ*_*s*_ because the 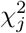’s are still within its expected values.

**Figure A4:**
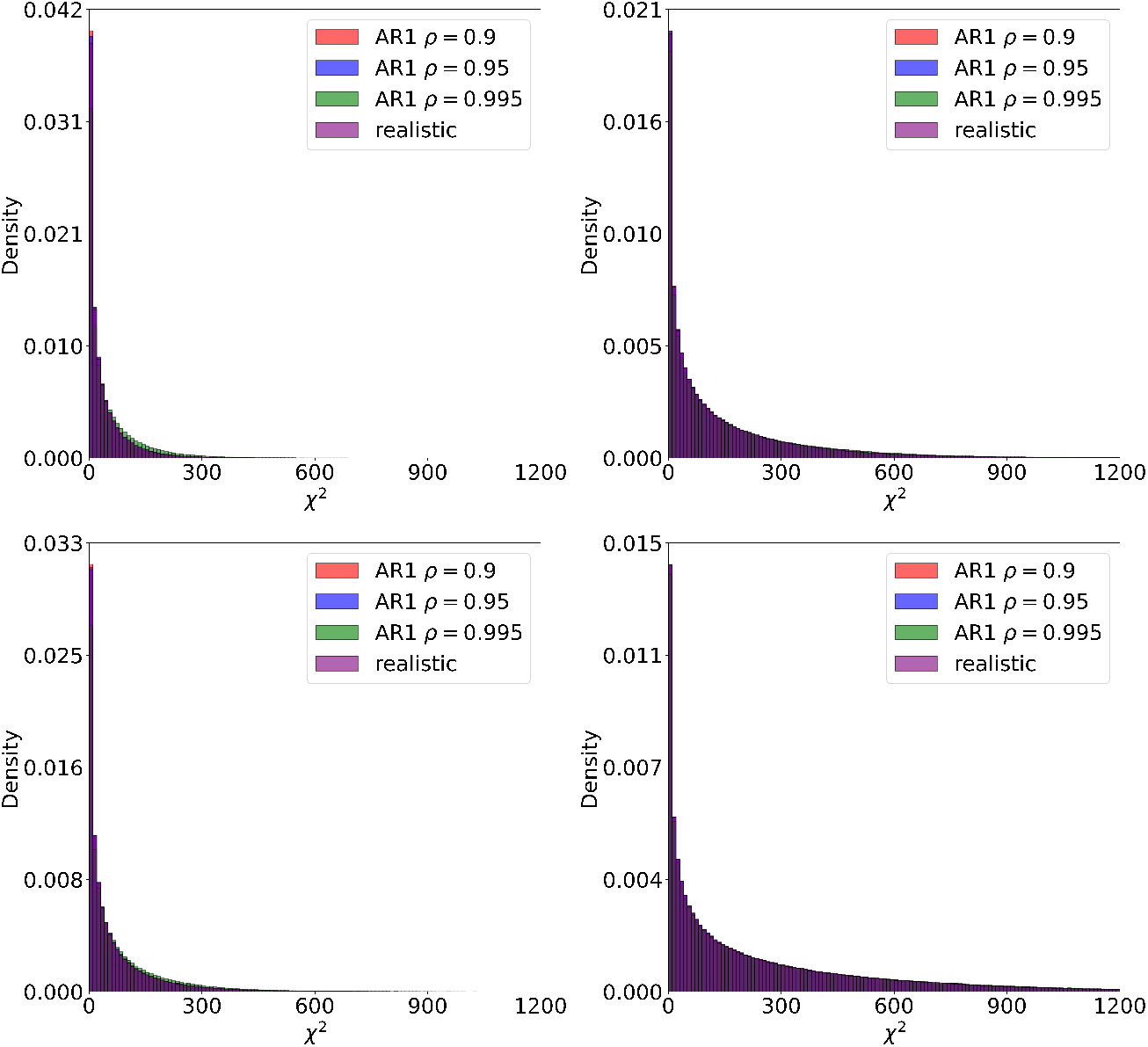
Histograms of 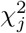 after increasing F_st_ and *σ*_*s*_. The top row represents F_st_ = 0.05 and the bottom row represents F_st_ = 0.1. The left column represents *σ*_*s*_ = 0.4 and the right column represents *σ*_*s*_ = 0.8. Increasing both parameters massively inflates 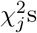 more than what was shown in Figure A3. The 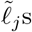 are exactly the same as in Figure A2 so they are not shown. Note that the scale of the y-axis of each panel is different.

We explore how 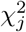’s might change when combining both F_st_ and *σ*_*s*_ effects. We do not examine the 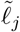’s in this setting because we see from Figure A3 that they remain unchanged when *σ*_*s*_ changes. It is observed in Figure A4 that increasing both F_st_ and *σ*_*s*_ inflates 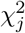’s more than the 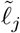’s. Because the 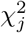’s are higher than expected in the regression model, the estimated slope is inflated resulting in inaccurate heritability estimation.

#### A1.2 Miscellaneous Simulations

**Figure A5:**
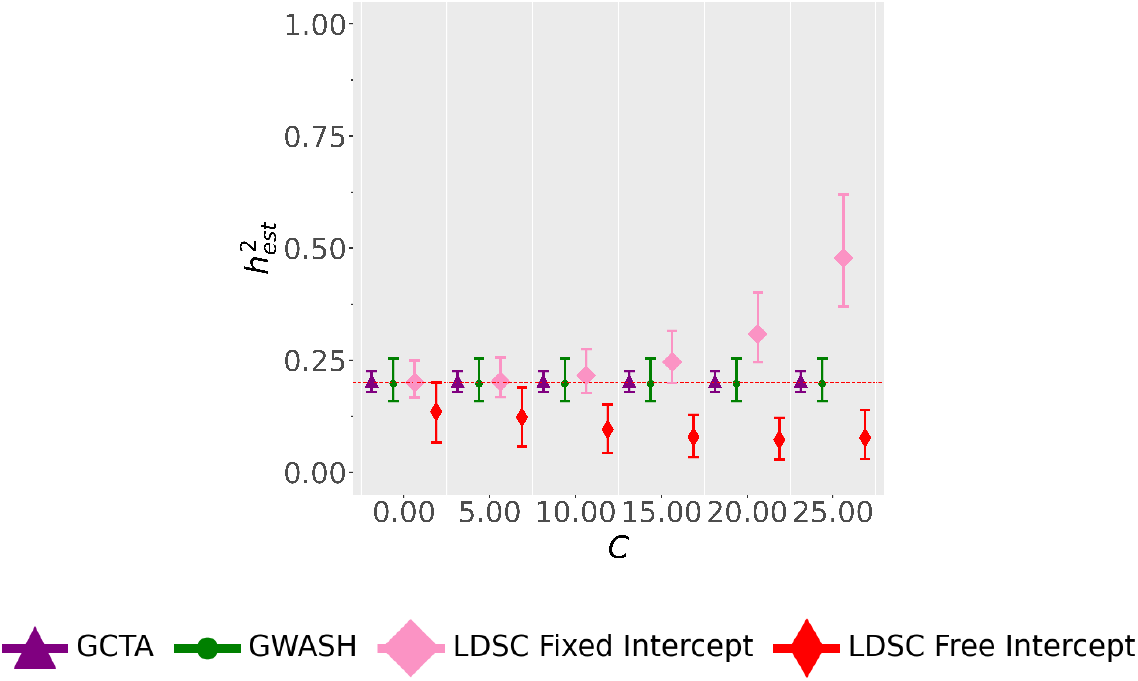
Heritability estimates when introducing ±*C* in LD scores of the reference panel 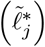. The first 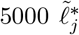 have *C* added while the remaining 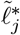 have *C* subtracted. Here, **X** is generated with realistic LD with *n* = 5000 and *m* = 10000 under 100 simulations. The 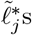 have the same distribution as 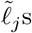.

We explore the effect of adding and subtracting a constant *C* to each half of 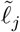’s. It is observed in Figure A5, that at *C* ≤ 5, LDSC with fixed intercept is robust to increases in LD scores. At larger values of *C*, we observe that LDSC with fixed intercept exhibits an upwards bias similar to that observed with an increase in F_st_. GWASH stays at the nominal value because the mean of the 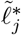’s remains the same. This illustrates that LDSC is sensitive to per-LD score changes due to regression-based fitting while GWASH is unaffected since it only depends on the mean.

**Figure A6:**
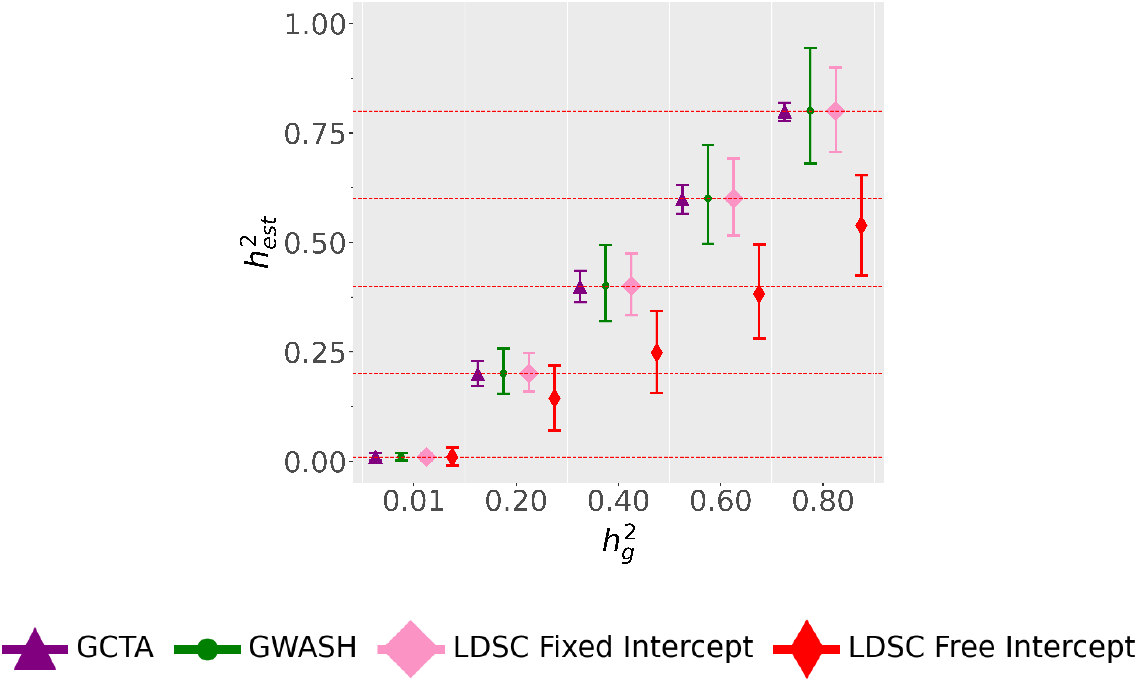
Heritability estimates when increasing 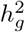 in the usual realistic simulation setting of *n* = 5000, *m* = 10000, no population stratification effects (*σ*_*s*_ = 0 and F_st_ = 0), and with realistic LD from Chromosome 22 1000 Genomes. There are 1000 simulation replicates and a reference panel matching the distribution of **X** is used. The red horizontal dotted lines represent the 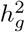 that the estimators are supposed to capture. LDSC with free intercept has an increased downwards bias as 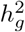 increases.

We explore the effect of changing the 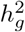 in realistic simulations with no population stratification effects. The results are plotted in Figure A6. At all levels of 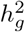, GCTA, GWASH, and LDSC with fixed intercept remains unbiased while LDSC with free intercept is biased down (expect for 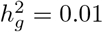).

## Supplementary

### S1 Corrections to the LD Score Regression Derivation

In this section we seek to clarify the LD score regression derivation. We derive corrected versions of the equations and discuss issues with the original derivation of [8], particularly under population stratification. In order to make this supplementary self-contained we repeat some of the definitions that appear in the main text.

#### S1.1 Notation and Conditioning

##### S1.1.1 Notation

We start our derivation by defining our notation, following that of [4]. In particular suppose that we observe data from *m* SNPs from each of *n* participants. Define the sample genotype matrix to be **X** ∈ ℝ^*n×m*^ where each row of **X** represents the genetic data for a given subject and let **y** ∈ ℝ^*n*^ be a vector of sample phenotypes.

We begin, for simplicity, by considering the model (See Equation (1) from Section 2.1 in the main text) without population stratification. In this case the relationship between sample genotype and phenotype can be represented in a linear model as follows:

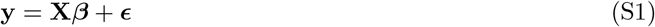

where ***β*** = (*β*_1_, …, *β*_*m*_)^*T*^ ∈ ℝ^*m*^ is the SNP effect size and *ϵ* = (*ϵ*_1_, …, *ϵ*_*n*_)^*T*^ ∈ ℝ^*n*^ is the error term. For each 1 ≤ *j* ≤ *m*, let **X**_*j*_ ∈ ℝ^*n*^ be the *j*th column of **X**. We make the following model assumptions (as in [4]).

1. For each 1 ≤ *j* ≤ *m*, **X**_*j*_ is normalized to be mean zero and variance one (note that as in [4] we ignore the distinction between normalizing and centering in our sample and in the population)
2. The rows of **X** - corresponding to the genotype data for each subject - are independent and identically distributed.
3. *ϵ*_1_, …, *ϵ*_*n*_ are iid with E (*ϵ*_*i*_) = 0 and 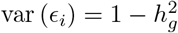.
4. *β*_1_, …, *β*_*m*_ are iid with E (*β*_*j*_) = 0 and 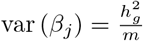.
5. **X, *β, ϵ*** are independent.

Define the estimated effect-size of the *j*th SNP 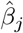 as:

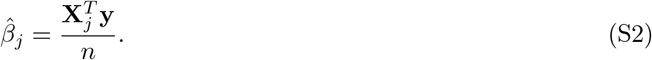

Define 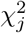 as:

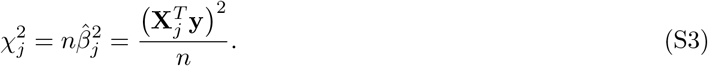

We also define the LD scores as follows. Let 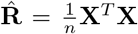 denote the correlation matrix with entries 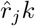. Then the sample LD scores are computed as:

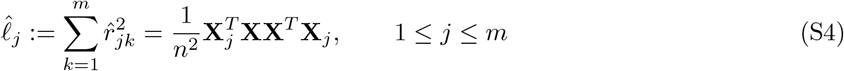

Letting *r*_*jk*_ = cov(*X*_1*j*_, *X*_1*k*_), population LD scores can be also defined as

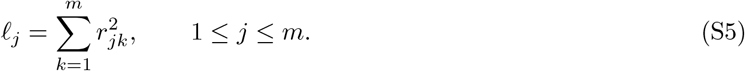

##### S1.1.2 Conditioning

Before continuing with our explanation of the LD score derivation we first make an observation regarding conditioning which in our view is approached incorrectly in [3]. The paper as a whole targets the unconditional expectation: 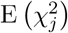 instead of the conditional expectation 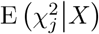 (see for instance the statements of Propositions 1 and 2 in [4]). This is problematic and means that the derivation of their estimator without population stratification is circular (as we demonstrate in Section S1.2). Furthermore it causes their derivation under population stratification to be incorrect, as we show in Section S1.3. This in turn means that using the original LD score regression equations leads to bias under population stratification, as we demonstrated in AR1 and Realistic LD simulations from Section 3 in the main text.

The reason that 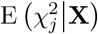 should be targeted rather than the unconditional expectation is that the summary statistics 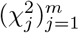 are calculated based on the sample **X**. There is thus not enough information in the system to infer on 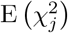 as that would require many samples of 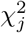 from different **X** matrices. Furthermore, regression that is based on the unconditional expectation leads to bias under population stratification as explained below in Section S1.3.1.

#### S1.2 LDSC derivation without population stratification

##### S1.2.1 LDSC Derivation

In what follows we re-derive the LDSC regression equation with the correct conditioning resulting in Theorem S1.1. This result is the analogue of Proposition 1 of [4], with conditioning correctly accounted for.

###### Theorem S1.1.

*Under Model* (S1),

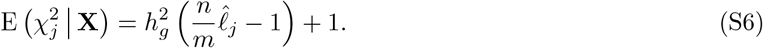

*Proof*. We first note that

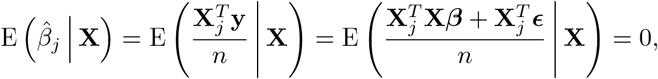

since E (***β***) = **0** and E (***ϵ***|**X**) = **0**. As such,

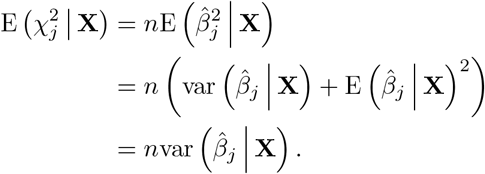

Now,

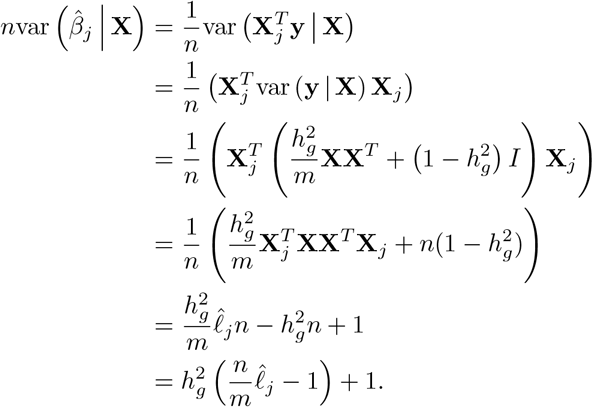

The proof follows the begining of Proposition 1 of [4]. However the end result is is exact rather than an approximation. The authors use an approximation of the Olkin and Pratt estimator for population squared correlation [13] combined with a *δ* method approximation in order to estimate the unconditional expectation. In particular they derive the following unconditional approximation.

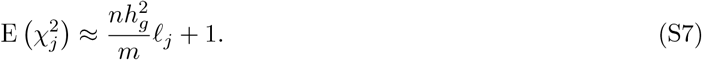

However, as we show in the next section, using the unadjusted LD scores in (S7) (as in done in practice in the implementation of [3]) actually recovers (S6) - meaning that the use of the approximation and (S7) is unnecessary.

##### S1.2.2 The circular use of approximations

The LD score equation derived in [4] (displayed in (S7)), depends on the population LD scores which are unknown. In order to get around this [3] propose taking

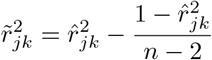

as this is an unbiased estimator of 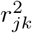. They then use these to compute adjusted LD scores,

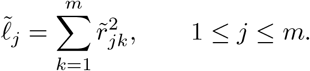

It follows that

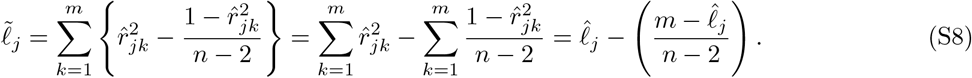

In the LDSC software these adjusted LD scores are then plugged into (S7) and used when performing LD score regression. As such, when 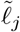 is plugged into the right hand side of (S7) in place of *ℓ*_*j*_ we obtain

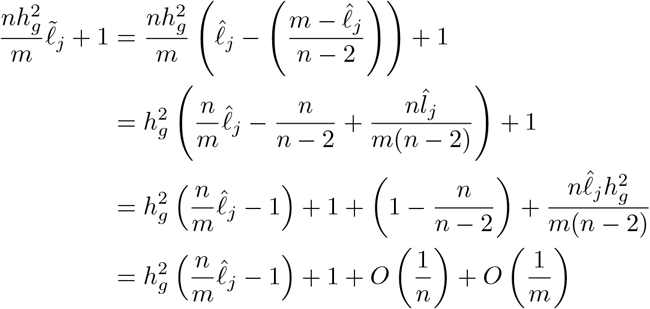

so we recover the right hand side of (S6) up to terms of a low order. This explains why using the unconditional equation, as done in [3] is okay in this case - since in practice the conditional equation is recovered.

###### Remark S1.2.

*The original proof of Proposition 1 of [4] starts off as in the proof of Theorem S1.1 then uses an approximation to derive* (S7) *and then uses a second approximation to estimate the LD scores. As we have shown doing all of this in fact returns us to the result of Theorem S1.1 and so the combination of the two approximations is circular. Since the conditional expression is the one that should be used, as we argued in Section S1.1.2, using these approximations are unnecessary*.

###### Remark S1.3.

*In this setting, using and then undoing the approximation recovers the conditional equation. As such in this simple setting where there is no population stratification the original formula is thus conditionally correct. However, as we shall show in Section S1.3, under the presence of population stratification the equation changes. Unfortunately this means that using the original formula can lead to bias, as demonstrated in Section 3 of the main text*.

#### S1.3 LDSC regression derivation under Population Stratification

##### S1.3.1 Modeling Population Stratification

To account for population stratification we follow the model of [4]. We assume that each subject *i* belongs with equal probability to one of two populations *P*_1_, *P*_2_, and

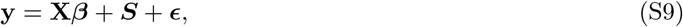

where *S*_*i*_|*i* ∈ *P*_1_ = *σ*_*s*_ and *S*_*i*_|*i* ∈ *P*_2_ = −*σ*_*s*_ for a constant 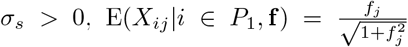 and 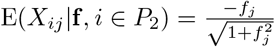, and 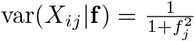, where the vector **f** ∈ R^*m*^ is iid with mean 0 and variance 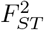. Instead of 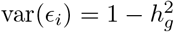 of Assumption 4 of Model (S1) we now assume that 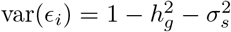 in order to keep var(*y*_*i*_) to be 1. The rest of the Assumptions of Model (S1) holds also here.

##### S1.3.2 A comment about the definition of S

The population stratification term **S** was originally defined in [4] as 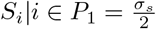 and 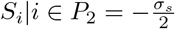, while in our definition we do not divide by 2. On the other hand, they assumed, as we do, that 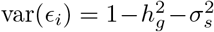, which implicitly requires that 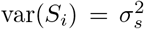 nbecause var(*y*_*i*_) = 1. However, according to their definition, 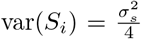, which makes their definitions inconsistent. To fix this mistake we define *S*_*i*_ as ±*σ*_*s*_ without dividing by 2.

##### S1.3.3 Varying *σ*_*s*_ in S does not change heritability estimation on average in even subpopulations

Varying *σ*_*s*_ does not change heritability estimator behavior on average in our simulations because there are an equal number of subjects in *P*_1_ and *P*_2_. This shifts **y** by ±*σ*_*s*_ over all *i* individuals which aggregates as 0:

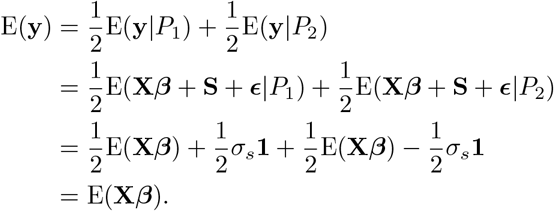

##### S1.3.4 A mistake in the derivation of [4]

The derivation [4] contains a critical mistake, which, as explained below and demonstrated in Sections 3.1 and 3.2 of the main text, causes their estimate to be severely biased when *σ*_*s*_ and 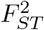 is large. In [4] on page 4 it is written that “we compute 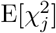 with the expectation taken over random **X, *β, ϵ*, f** but with **S** fixed to ensure population stratification.” However, Eq. (2.11) of [4] reads 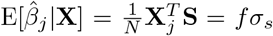 (in our notation *n* = *N*); the first equality is correct, but the second is wrong as **X** is conditioned upon and **S** is fixed. Hence, 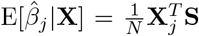. As we shall see below, this mistake results in bias of the suggested estimator of [4].

##### S1.3.5 The conditional expectation of 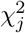 and the resulting bias of the estimator

The equivalent result of Theorem S1.1 under Model (S9) is given now. To make the argument clear, the conditioning on **S** is explicit in the notation.

###### Theorem S1.4.

*Under Model* (S9),

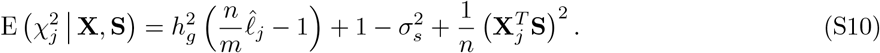

*Proof*. We proceed as in the proof of Theorem S1.1. As mentioned in Section S1.3.4,

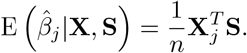

The computation of *n*var 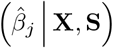 is almost the same as in the proof of Theorem S1.1, besides that now 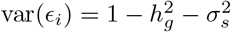 (while under Model (S1), 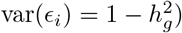). It follows that

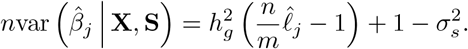

Putting the terms together in

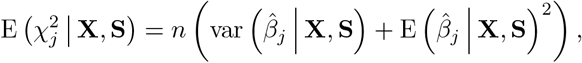

yields (S10).

The key term in the conditional expectation (S10) is the last term 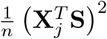, which was not calculated properly in [4] as mentioned earlier. This term is correlated with 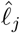, which leads to bias estimator as explained next.

Theorem S1.4 demonstrates a conceptual mistake in the derivation of [4]. As mentioned above they calculate the unconditional expectation of 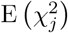. However writing,

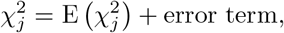

Theorem S1.4 implies that the error term is correlated with 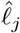. Therefore, when 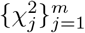 is regressed on the LD-scores 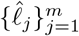, the resulting estimator will be biased due to this correlation. On the other hand, when one considers the conditional expectation 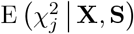 the error term is not correlated with **X** and **S** by definition. Thus, one should base the regression estimator on the conditional expectation rather the unconditional one - not doing so can lead to bias.

###### Remark S1.5.

*As we observed in Section 3 and Figures 1 and 2 from the main text, LDSC is biased under population stratification despite [3] claiming that adding the bias correction term should resolve this. We have now explained this as an issue with the derivation and fact that the error is correlated with the LD scores. The bias of the LDSC estimator is calculated explicitly in Theorem 9 of [1], where it is shown that it is asymptotically equal to* 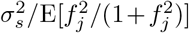. *Note that this issue does not affect the model without population stratification because the error in the conditional expectation* (S6) *is not correlated with the LD scores (and indeed because as argued in Section S1.2.2 the actual implementation of the LDSC regression estimator with the fixed intercept reduces to the conditional version)*.

### S2 Implementation Details

#### S2.1 LDSC Implementation

##### S2.1.1 LDSC Equation in Practice

The current implementation of LDSC is built with functional annotations in mind [7]. While *m* in Equation (8) in the main text is meant to be interpreted as a scalar value of all SNPs in the summary statistic, *M* in the current release of the LDSC code is actually a *c* × 1 vector which sums to *m* where *c* is the number of functional annotations and each element of *M* is the number of SNPs per *c* annotations. Each SNP can have different sample sizes due to genotyping or quality control [9]. This is represented as *N*, a *m* × 1 vector of sample-size per genotyped SNP. LDSC uses the average of *N* as *n* in Equation (8).

##### S2.1.2 LDSC Method Overview

LD Score Regression heritability is computed in two steps (three steps in actuality). For this explanation, we assume that 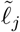 is observed. If not, then it can be substituted with 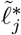 as described in the main text.

0. A “crude” estimate of heritability is calculated by a ratio of means: The mean of 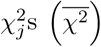 on the mean of LD scores 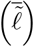. This is known as the “aggregate” heritability estimate and is given by the following expression:

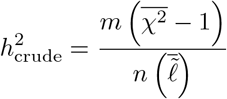

This heritability estimate is susceptible to population stratification effects and is used to compute initial weights. The weighting stage consists of the following:
  a. Heritability is bound between 0 or 1.
  b. The minimum value *ℓ*_*j*_ can take is 1 and is set to 1 if below it.
  c. Initial weights are computed:

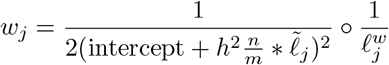

Where 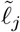 represents the LD scores of specific SNPs of interest. 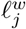 are LD scores of the same SNPs from potentially another trusted panel (such as HapMap3 SNP LD scores). When the intercept is not yet estimated, intercept = 1 in *w*_*j*_. In LDSC, weighting is done for the following reasons:
    i. The HapMap “regression SNPs” LD scores come from LD scores computed from JUST HapMap3 SNPs. This is because *χ*^2^ statistics used in the regression are not independent.
    ii. Heteroskedasity in SNPs meaning that *χ*^2^ of SNPs with high LD have higher variance than *χ*^2^ if SNPs with lower LD so SNPs with high LD scores are down-weighted.
    In our scenarios, we do not explore the inclusion of a trusted reference panel and we use all SNPs so 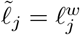. Further, LDSC “is not sensitive to the precise choice of” 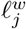[6].
1. Starting with the inital weights, iterative weighted least squares is performed on SNPs with 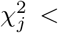 twostep (default: twostep = 30).
  a. Do weighted least squares by weighting 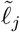 by *w*_*j*_ and 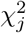 by *w*_*j*_ and fitting the regression. In this step, 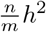 and an intercept is estimated.
  b. From the weighted least squares results, the new *h*^2^ and intercept are used to compute new weights. This occurs twice.
  c. The 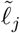 and 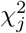 are weighted by the final weights from the above step and a final weighted least squares regression is fit.
2. Using the intercept from the previous step, do iterative weighted least squares on the whole dataset (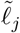 and 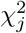 unfiltered) with the starting initial weights (steps a through c in step 1).

From the results of the iterative weighted least squares, the “jackknife process” is conducted. Jackknife is usually used to get standard errors by leaving-one-out. The implementation outputs the estimate during this process as well. We follow typical linear regression notation where X represents the design matrix and y represents the response, not genotype and phenotype respectively. In a very simple case, X is a *m* × 2 matrix with a column of ones and 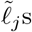 and y is an *m* × 1 vector of 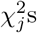. The following steps occur:

1. Blocks (default: 200) of `X` and `y` are created. For each *k*th block, calculate (`X`^*T*^ `X`)_*k*_ and (`X`^*T*^ `y`) _*k*_.
2. Sum across all (`X`^*T*^ `X`) _*k*_ s and (`X`^*T*^ `y`) _*k*_ s to make total `X`^*T*^ `X` and total `X`^*T*^ `y`. Calculate 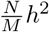 from the OLS Normal Equation:

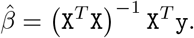
3. Create delete values. The *k*th delete value 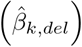 is the 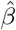 calculated after subtracting the *j*th `X`^*T*^ `X` from the total `X`^*T*^ `X` and subtracting the *k*th `X`^*T*^ `y` from the total `X`^*T*^ `y`.
4. Create pseudovalues. The *k*th pseudovalue (*P*) is the impact of removing the *k*th block from the total calculations:

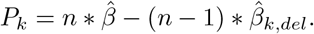

The Pseudovalue matrix (**P**) is a *k* × 2 matrix since two estimates are computed: the intercept and the heritability estimate. The covariance matrix is computed as:

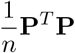

and the diagonals are the block-jackknifed variance of each parameter. The reported standard error in the LDSC output is the square-root of the diagonals.

##### S2.1.3 LD Score Comparisons

We constructed LD scores from 1000 Genomes Phase I data as described in [3] directly using the LDSC package. The individual that was excluded “from a pair of cousins” in [3] was not identified so we created LD scores of chromosome 22 where each individual was excluded iteratively and compared the output against the released LD scores of chromosome 22 via MSE. The individuals in these LD scores with the lowest MSE of 0.003844 were used to construct the LD scores for the remaining chromosomes. The heritability estimates from our constructed LD scores with the lowest MSE closely matches those estimated with the released LD scores.

##### S2.1.4 Effect of the twostep Parameter in LDSC

The intercept is first estimated by fitting a regression with SNPs that have 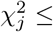 twostep, where twostep is a predetermined threshold. The rationale for this is that SNPs with small 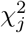 under this threshold are less likely to be causal [2]. The twostep parameter is set by default in LDSC as 30.

**Figure S1:**
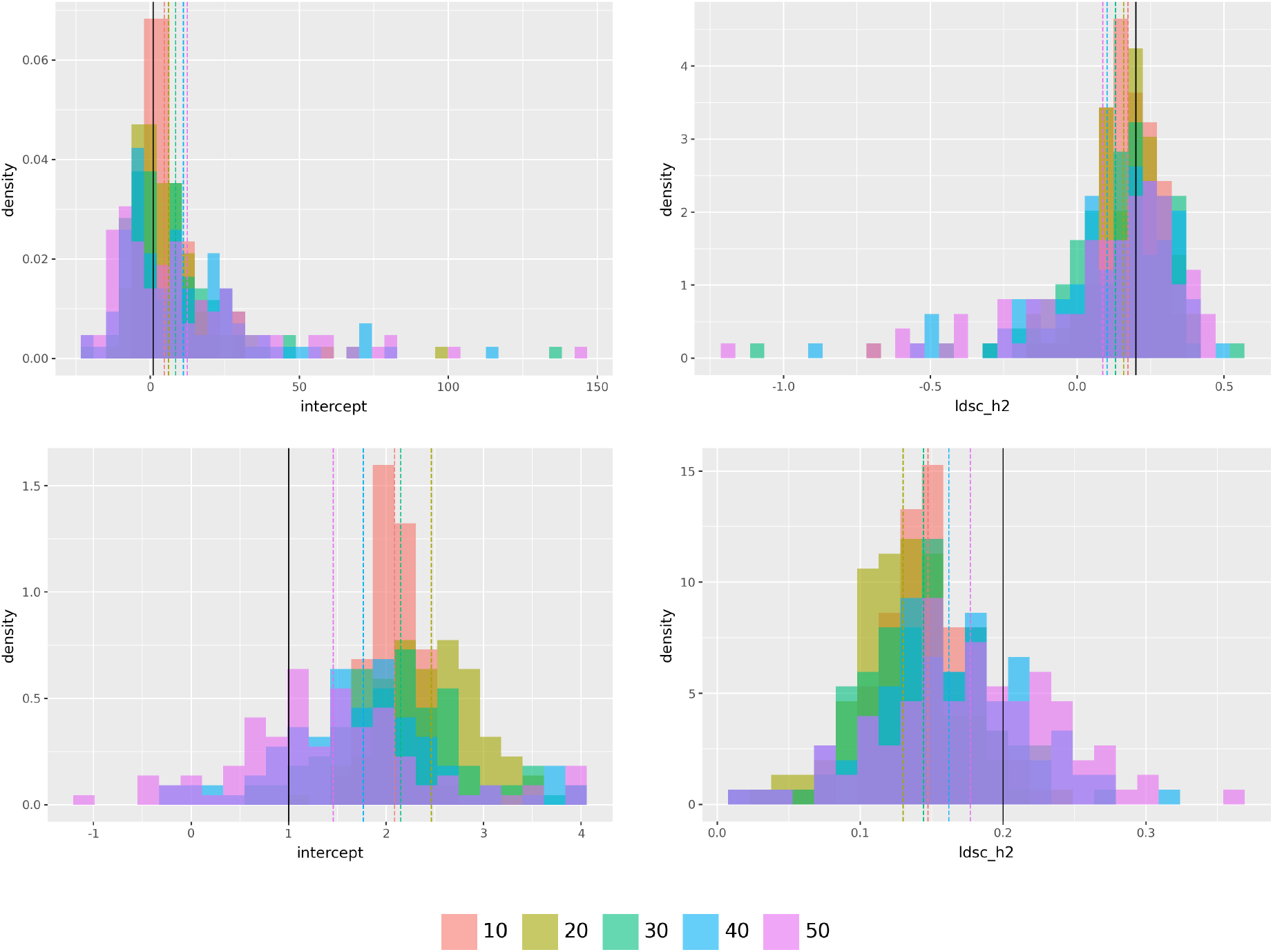
Histograms of LDSC estimated intercepts and *ĥ*^2^ in AR1 *ρ* = 0.995 (top row) and Realistic (bottom row) settings of 100 simulations with varying twostep (colors) and no population stratification. The black line in the intercept histogram represents an intercept of 1 and the black line in the heritability histogram represents 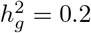. The dashed lines are the average quantity across the 100 simulation replicates. Note that the density scale across the histograms are different.

In the AR1 setting, increasing twostep results in a larger intercept on average. In the realistic setting, a large twostep results in a smaller intercept. The intercept is estimated to be larger than 1 under the default setting of twostep = 30 resulting in a downwards bias of *ĥ*^2^ in both settings. Increasing the twostep parameter in the AR1 setting increases the intercept away from 1 and biases the heritability estimate downwards away from 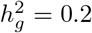, on average. Unlike the AR1 setting, a clear trend is not present in the realistic setting with respect to the twostep parameter. We do note that a twostep = 50 results in an intercept closest to 1 and a heritability estimate closest to 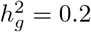, on average.

#### S2.2 GCTA Implementation

We compare our implementation (Python GCTA) to the released implementation (Native GCTA) from [12]. We measured the computation time of each algorithm and its heritability estimate across 10 simulations with *n* = 5000, *m* = 10000 1000G chr 22 SNP correlation. On an Intel computational platform, the computation time is comparable:

| Implementation | $\hat{h}^2$ | Iterations to Converge | Computation Time (s) |
| --- | --- | --- | --- |
| Native GCTA | 0.1950 (0.0279) | 10.000 (0) | 51.4160 (0.6205) |
| Python GCTA | 0.1950 (0.0279) | 8.6000 (0.5477) | 66.6420 (4.2146) |

On an AMD platform, the time needed for the Python implementation is ~ 2.7 times worse:

On an AMD platform, the time needed for the Python implementation is $\sim 2.7$ times worse:
| Implementation | $\hat{h}^2$ | Iterations to Converge | Computation Time (s) |
| --- | --- | --- | --- |
| Native GCTA | 0.1950 (0.0279) | 10.0000 (0) | 39.5220 (0.2372) |
| Python GCTA | 0.1950 (0.0279) | 8.6000 (0.5477) | 107.6062 (10.7435) |

We saw that despite having heritability estimates that are really close ~10^*−*7^, there is a difference in computation time between the two implementations favoring Native GCTA. The advantage of using the

Python GCTA implementation is that it integrates seamlessly in our framework, and that we do not need to consider computational overhead of writing/formating the simulated data as typical input files for Native GCTA (ie: GRM, .pheno files) and reading on the spot.

##### S2.2.1 GCTA Algorithm

The inputs for this are:

- *y*, the phenotype individual-level data, an *n* × 1 vector.
- **K** = **XX**^*T*^, the kinship matrix (also called the GRM in [12]), an *n* × *n* matrix constructed from individual-level genetic data **X**.

The GCTA algorithm estimates 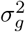 and 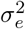 iteratively using the Expectation Maximization (EM) algorithm and Average Information (AI) algorithm. The EM algorithm as the initial step to determine the direction of iteration updates and AI algorithm for the remaining steps until convergence [12]. The original implementation of GCTA considers multiple variance components per region of interest (ie: chromosomes) [12], but for our study we decided to look at only one region. Therefore, our scheme here only considers 2 variance components whereas the original paper has a generalized derivation for multiple variance components.

We start with the initial EM algorithm in the following steps:

1. Start with guesses: 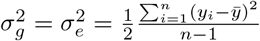.
2. Construct 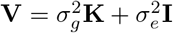. This is an *n* × *n* matrix.
3. Construct projection matrix **P** = **V**^*−*1^ − (**V**^*−*1^**A**)**A**^*T*^ **V**^*−*1^**A**^*−*1^**A**^*T*^ **V**^*−*1^. This is an *n* × *n* matrix. The original GCTA implementation examines *c* non-genetic covariates for each *n* subject. This is represented as **A**, a *n* × *c* matrix. Since we do not examine these in this work, we omit explicit mention from the main text and this is just an intercept (ie: a column of ones).
4. Calculate the log-likelihood (*L*) given **V** and **y**. This is:

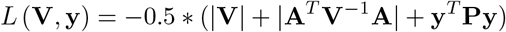

where |**V**| is the determinant of **V** and the other symbols are defined as before.
5. Compute the initial update of 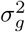 and 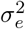.

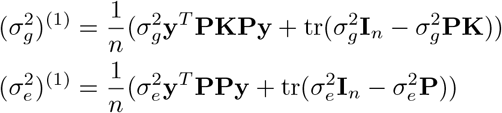

The 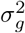 and 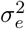 from the EM algorithm are used as the starting points for the AI algorithm. The AI algorithm runs until abs ((*L* (**V**, *y*))^(*t*)^ − (*L* (**V**, *y*))^(*t−*1)^) *<* 10^*−*4^. For each *t* iteration:

1. Check if 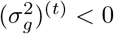. If it is, then set 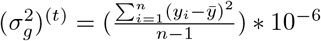
2. Check if 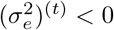. If it is, then set 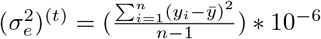
3. Compute 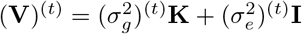
4. Compute (**P**)^(*t*)^ = (**V**^*−*1^)^(*t*)^ − ((**V**^*−*1^)^(*t*)^**A**)**A**^*T*^ (**V**^*−*1^)^(*t*)^**A**^*−*1^**A**^*T*^ (**V**^*−*1^)^(*t*)^.
5. Compute *L* (**V**, *y*))^(*t*)^ = −0.5 * (|(**V**)^(*t*)^| + |**A**^*T*^ (**V**^*−*1^)^(*t*)^**A**| + *y*^*T*^ (**P**)^(*t*)^*y*).
6. Check if abs (*L* (**V**, *y*))^(*t*)^ − (*L* (**V**, *y*))^(*t−*1)^ *<* 10^*−*4^. If this statement is true, stop; otherwise, go to the next steps.
7. Compute AI matrix:

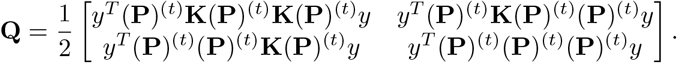
8. Take the derivative of the log-likelihood function with respect to 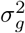 and 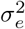:

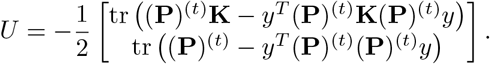
9. Update parameters:

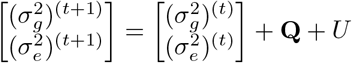

Go back to step 1 of the AI algorithm with (*t* + 1) parameters until the stopping rule is met.

#### S2.3 Computational Efficiency

##### S2.3.1 Efficient Computation of LD scores from Non-scaled Kinship Matrix

The correlation matrix is represented as 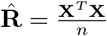. Recall that the LD Score of each *j*th SNP is the sum of all correlations to other *k* SNPs around it and is computed by squaring each element 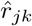 in 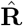 and summing across rows:

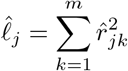

This is equivalent to computing the diagonal of 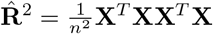:

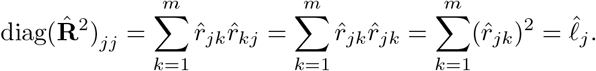

Since only the diagonal of 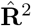 is needed, the operation is simplified using the Kinship Matrix: 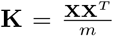 and the *j*th column of **X** instead of computing the full matrix:

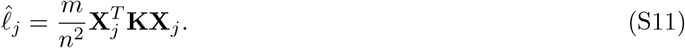

##### S2.3.2 Efficient Computation of 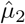 and 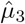

As shown in [10] and in our work here, 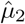 and 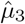 from Section 2.3.3 require the computation of 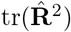 and 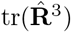 respectively where 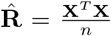 is the *m* × *m* correlation matrix and **X** is the usual *n* × *m* genotype matrix.

We start with a general case to compute the trace of 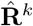:

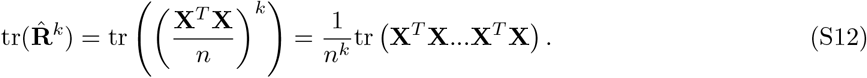

We use the cyclic property of traces, where for any conformable matrices **A** and **B**, tr(**AB**) = tr(**BA**) (see Equation 14 in [8]). Moving **X** at the end to the front, Equation (S12) becomes:

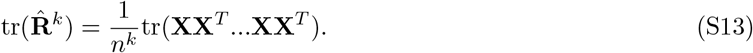

The kinship matrix is defined as 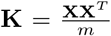 and is *n* × *n*. It can be seen from this that **XX**^*T*^ = *m***K**. Then, (S13) becomes:

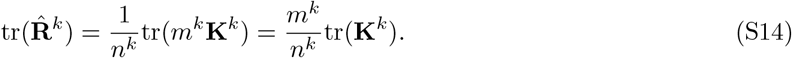

Computation of **K**^*k*^ is much more manageable as *n << m*. To compute 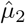 and 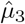 is set to 2 and 3 respectively.

The following table summarizes the average computational advantage of utilizing this trick over a direct computation across 100 simulations of **X** with AR(*ρ* = 0.2) correlation structure.

**Table 2:** Comparison of kinship method and direct computation for computing 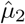 and 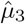. The kinship method offers a near 64 fold improvement for 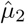 and a near 61 fold improvement for 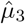 compared to the direct calculation. The absolute differences between the estimates are extremely small.

| Estimator | Computation Time Direct $\hat{\mu}_2$ (s) | Computation Time Kinship $\hat{\mu}_2$ (s) | Absolute Difference |
| --- | --- | --- | --- |
| $\hat{\mu}_2$ | 0.1888 | 0.0029 | 5.4401e-16 |
| $\hat{\mu}_3$ | 0.3140 | 0.0051 | 3.0531e-15 |

##### S2.3.3 Efficient Computation of Realistic X

Given a real genotype *n* × *m* matrix **X**^*\**^ that acts as a reference panel, we want a desired **X** simulated with similar LD structure that has dimensions *k* × *m*. Here, *n* is the sample size of the real genotype matrix **X**^*\**^ and *k* is the sample size of the desired simulated genotype matrix. Both **X** and **X**^*\**^ must share the same *m* SNPs.

In twas sim [11], **X**^*\**^ undergoes pre-processing to ensure that 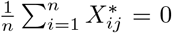 and 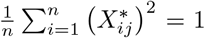 for all *j*. Then, **X** is generated from a lower-triangular of (**X**^*\**^)^*T*^ **X**^*\**^, which is denoted by **L**, and a matrix **Z** of dimensions *k* × *m* with iid standard normal entries independent of **X**^*\**^:

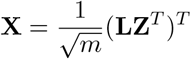

This induces correlation between columns following the LD structure in **L**. It is easy to see that in expectation **X**^*T*^ **X** is equal to (**X**^*\**^)^*T*^ **X**^*\**^:

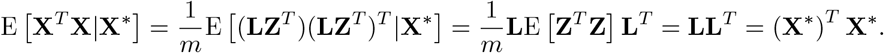

A well-known requirement of using the Cholesky Decomposition method is that the matrix must be positive definite. To ensure that (**X**^*\**^)^*T*^ **X**^*\**^ matches such a condition, a diagonal matrix of some arbitrary coefficient ld_ridge_ is added as a correction matrix [5] and the matrix (**X**^*\**^)^*T*^ **X**^*\**^ is standardized by 1 + ld_ridge_ to ensure that the diagonal of the new corrected LD matrix is 1. If ld_ridge_ is small, the off diagonals should be offset by a near-negligible factor. To do Cholesky Decomposition in twas sim, ld_ridge_ is set to 0.1, and this value can significantly offset the off-diagonal values. Further, **X** is standardized such that 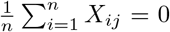 and 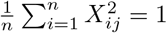 for all *j*.

We show that the same objective can be met without doing Cholesky Decomposition and eliminating this correction cost. Specifically, let **Z** be a *k* × *n* matrix with iid standard normal entries independent of **X**^*\**^, and define

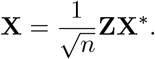

Repeating a similar calculation as above we have

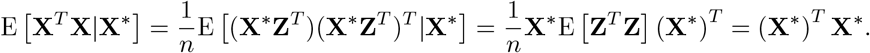

Then, the matrix **X** undergoes the same standardization as that from twas sim to ensure 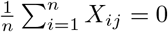 and 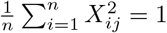 for all *j*.

#### S2.4 Modifications to GWASH to match LDSC in Real Data

The goal of this section is to clarify implementation details to enable a direct comparison between LDSC and GWASH. This direct comparison is possible because GWASH should be close to LDSC with fixed intercept as discussed in [10]. These implementation details are the following:

1. To mirror the analysis done in the LDSC package, in the LDSC calculation we set 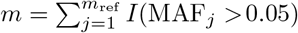, where *m*_ref_ is the number of SNPs in the reference panel and MAF_*j*_ is the minor allele frequency for the *j*th SNP. Note that this *m* refers to the number of SNPs used in the calculation of LD scores, and is therefore different from the number of SNPs that overlap between the reference panel and dataset. This is only done in the real data analysis as we assume in simulations that all SNPs in the reference panel match those in the simulated dataset and meet the MAF_*j*_ *>* 0.05 criteria.
2. We multiply 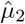 (see Equation (10)) and 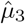 (see Equation (13)) by a factor of 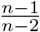. We denote these quantities as 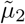 and 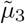 which consider the bias correction discussed in [3]. This is done in both real data and simulations.
3. In 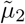 and 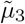, *m* is used instead of *m* − 1 because of how 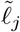 is computed. This is only done in the real data analysis.
4. The quantities 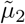 and 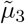 are computed with a 1 Centimorgan window similarly to how this window is used in LDSC. We then compute 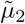 and 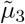 by chromosome as specified in [10]. This is only done in the real data analysis.

We discuss each point in a respective section below.

#### S2.4.1 Rationale of using 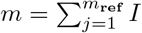 (MAF_*j*_ *>* 0.05) in LDSC

In the LDSC software package [3], 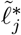 from a reference panel must be prepared. Each 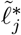 is computed with all SNPs found in the reference panel which may not include SNPs found in the dataset. Then, *m* is set to 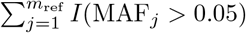 (MAF_*j*_ *>* 0.05) from the reference panel to account for the “missing” SNPs not found in the dataset that contribute LD in 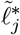. The value of *m* in the LDSC calculation refers to the number of SNPs used in the calculation of LD scores, not the number of SNPs that overlap between the reference panel and dataset. If *m* was to be set to the number of SNPs that overlap the reference panel and dataset, this might not accurately represent 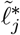(as this considers the mismatched SNPs). It could also inflate *ĥ*^2^ because the dominator is smaller since *m*_ref_ *> m*. In simulations, we assume the ideal case that all SNPs in the reference panel are found in the dataset and that all SNPs have MAF_*j*_ *>* 0.05.

#### S2.4.2 Computation of 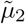 and 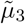

We make the deliberate choice of adjusting the GWASH equations in the implementation by a factor of 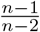. This is because the LD scores 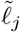 (as defined in Equation (3)) have a slightly different bias correction term than that presented in [10]. At low *n*, such as with the 1000 Genomes Phase 1 reference panel with an *n* = 378, this discrepancy is notable. We show this by examining the relative difference across different *n* sample sizes in simulation.

**Figure S2:**
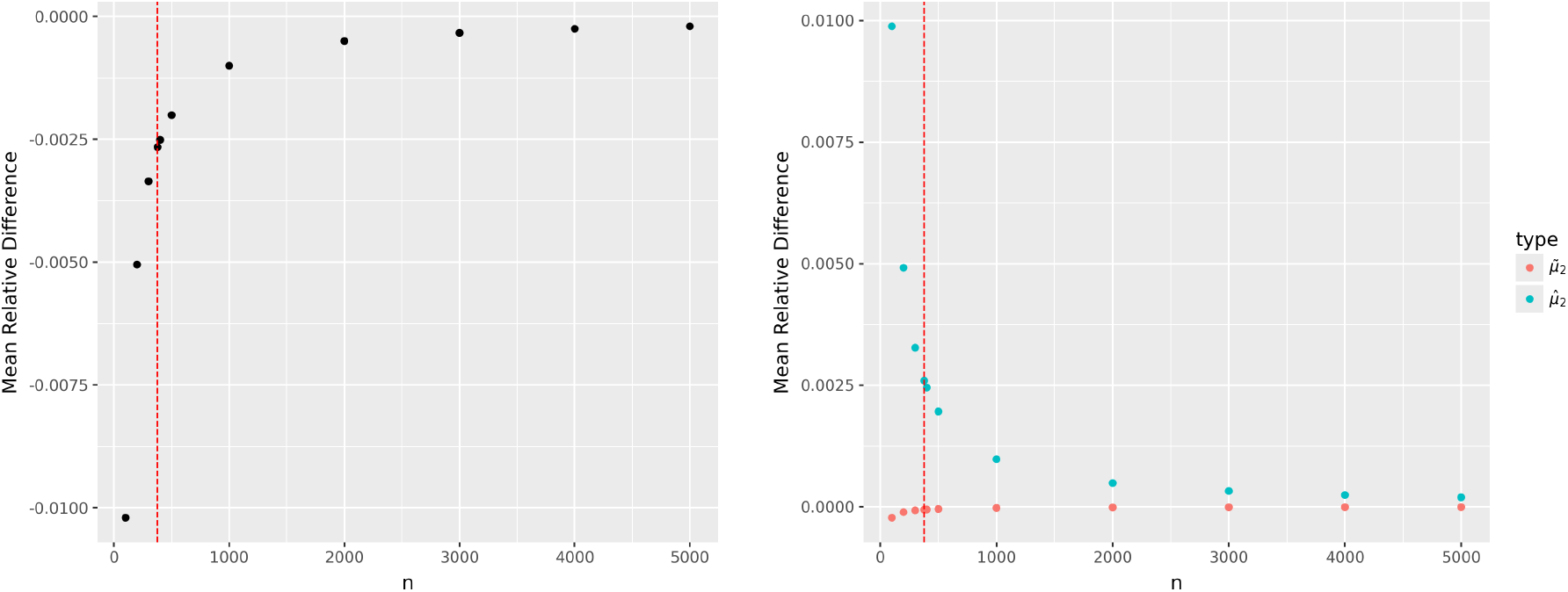
Mean relative difference between 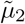 and 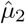 across 100 realistic LD simulations of *m* = 10000 across varying *n* (left panel) and mean relative difference between these quantities and the average of 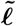. As *n* increases, the difference of the mean of 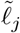 with 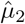 of the 2019 equation decays exponentially. From the right panel, we can see that the mean relative difference of both 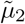 and 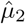 against the average of 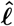 decreases towards 0 as *n* increases. Note that the y-axis for both panels is not shared. The red dotted line in both panels represents the sample size of the actual 1000G Phase 1 reference panel.

Recall that 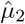 is written as the average of 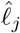 ‘s and we define 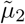 as the average of 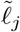 ‘s, i.e., 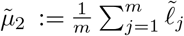. In the left panel of Figure S2, the relative difference between 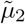 and 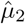 is computed as 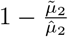. In the right panel, the relative differences with respect to 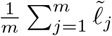 are computed as

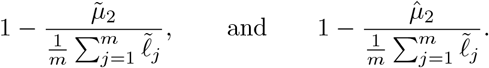

The difference between 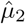 from the implementation in [10] and the mean 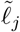 is quite substantial at *n* = 378. If [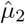 is not adjusted, LDSC and GWASH with real data are not comparable.

Below, we show that 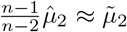. We use the result from the middle part of Equation (S8) in Section S1.2.2:

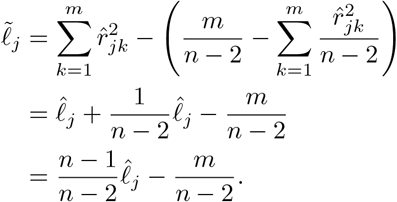

Now,

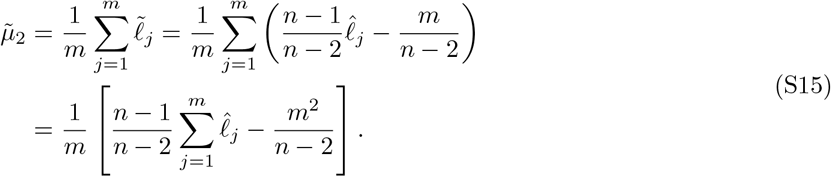

We have that 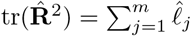 because

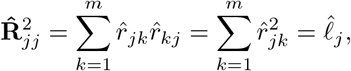

and therefore 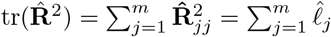. Equation (S15) becomes

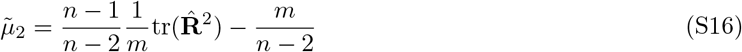

and one can see that Equation (S16) is approximately Equation (10) scaled by a factor of 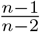:

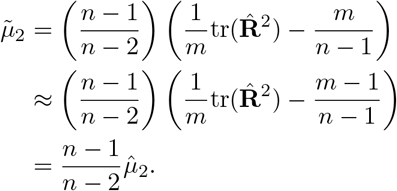

The 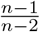 term is also multiplied to 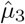 in our implementation:

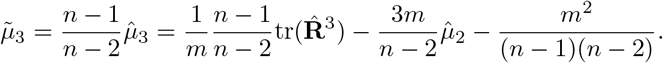

Notice that *m* is used instead of *m* − 1 as written in Equation (10) in the main text. This comes from the usage of 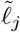. We address this in the next section.

##### S2.4.3 Use of *m* instead of *m* − 1 in 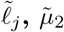, and 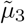

We also note that the equations for 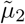 and 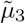 presented in this implementation and [10] have *m* − 1 instead of *m* as shown in 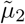 and 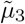. We justify using *m* for the real dataset analysis to directly compare GWASH and LDSC since we use the released precomputed LD scores while we use *m* − 1 as written in the main text equations for the simulations.

If *m* is replaced by *m* − 1 in the original equation for 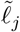 (S2.4.2), then the computation of 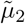 from individual-level data above will equal the average of all 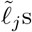 within a numerical accuracy of ~ 10^*−*15^. We show this empirically when varying *n* and *m* of 1000 simulations. We assume that the reference panel and dataset have the same sample size and number of SNPs.

**Table 3:** Comparison of 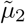 computed from taking the mean of 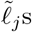 against 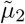 computed directly from **X** across different *n* and *m* using *m* as done in [3] or *m* − 1 as done in [10]. This analysis was conducted over 1000 simulations.

| n | m | use $m-1$ | $\tilde{\mu}_2$ from $\tilde{\ell}_j$ | $\tilde{\mu}_2$ from reference $\mathbf{X}$ | absolute difference |
| --- | --- | --- | --- | --- | --- |
| 50 | 100 | True | 1.9211e+01 (5.2815e+00) | 1.9211e+01 (5.2815e+00) | 2.9061e-15 (2.8765e-15) |
| 50 | 100 | False | 1.9190e+01 (5.2815e+00) | 1.9211e+01 (5.2815e+00) | 2.0833e-02 (4.1715e-15) |
| 500 | 1000 | True | 1.9480e+01 (5.8048e-01) | 1.9480e+01 (5.8048e-01) | 2.5224e-15 (2.4777e-15) |
| 500 | 1000 | False | 1.9478e+01 (5.8048e-01) | 1.9480e+01 (5.8048e-01) | 2.0080e-03 (3.1034e-15) |
| 5000 | 10000 | True | 1.9510e+01 (5.8228e-02) | 1.9510e+01 (5.8228e-02) | 2.4798e-15 (2.2782e-15) |
| 5000 | 10000 | False | 1.9510e+01 (5.8228e-02) | 1.9510e+01 (5.8228e-02) | 2.0008e-04 (3.2764e-15) |

Across 1000 simulations, the average absolute difference of 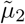 decreases by a factor of 10. This is because *m* − 1 ≈ *m* as *m* increases. We emphasize that *m* − 1 in the equations of the main text is replaced with *m* for the real data analysis but is as written for simulations.

##### S2.4.4 Computation of 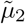 and 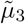 by chromosome on 1000 Genomes Data with 1 Centimorgan Window

To directly compare GWASH and LDSC on real datasets, the quantities that are used in each estimator must be preprocessed in the same way. We follow the strategy outlined in [10] to compute the LD contributions within each chromosome and then combine the results across them. As also done in LDSC, for each *j*th SNP, we compute 1 Centimorgan SNP correlation blocks, apply the bias correction as discussed previously in Section S2.4.2, and create 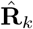 for each *k*th chromosome. We then compute the contribution of each *j*th SNP to 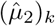 for each *k*th chromosome:

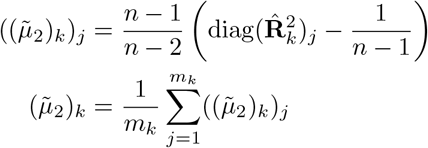

where 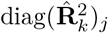 is the *j*th entry of the diagonal of 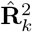, the covariance matrix of SNPs in chromosome *k*. We also do a similar operation for 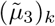:

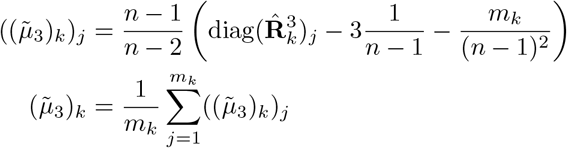

where *m*_*k*_ is the number of SNPs on the *k*th chromosome. We note that *m*_*k*_ is used here instead of *m*_*k*_ − 1 as written in Equation (10) and Equation (13) to facilitate direct comparisons to LDSC. We store 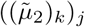 and 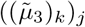 in separate tables that can be readily loaded. We then compute 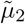 by computing the weighted average of all 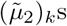 across the *k* chromosomes:

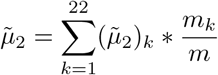

and 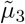 as a weighted average of 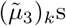:

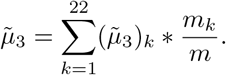

The code to compute all of these quantities described can be found on our GitHub repository (see Section 7).

#### S2.5 GWASH SE formula breaks at high *ρ*

The standard error formula for GWASH from [10] works when *ρ <* 0.8. At higher *ρ* values, we see that the GWASH standard error formula underestimates. The true GWASH se is calculated from the exact AR1 *ρ* correlation matrix and 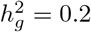.

**Figure S3:**
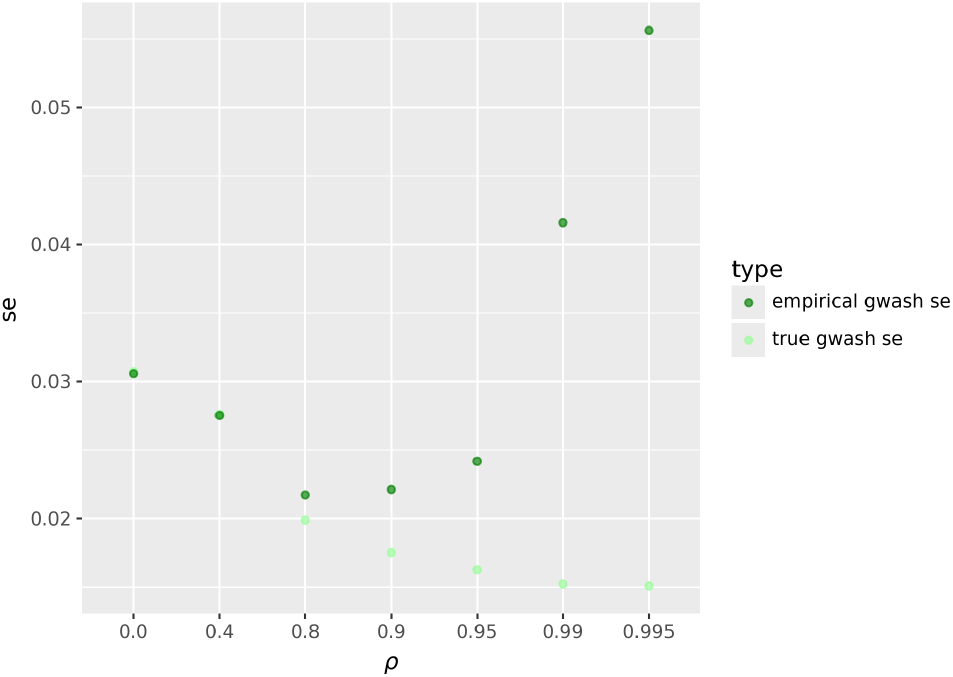
Comparison of empirical GWASH se from simulation against GWASH se computed with the true AR1 correlation matrix and 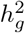 = 0.2. The previous study [10] only looked at *ρ* ≤ 0.8 where the formula and empirical se mostly aligns. At higher *ρ* regimes, the formula underestimates the empirical se from simulations.

The observed AR1 simulation standard errors are underestimated because the default *ρ* for the AR1 simulations is 0.995. Although the Realistic Simulations do not have AR1 correlation structure, we see the same underestimation effect in the estimated standard errors.

### S3 prop_causal_ Simulations with Reference Panel

We observed that changing prop_causal_ has no effect on heritability estimation in both AR1 and realistic LD simulations. These figures are shown here.

#### S3.1 AR1 prop_causal_ Simulations with reference panel

**Figure S4:**
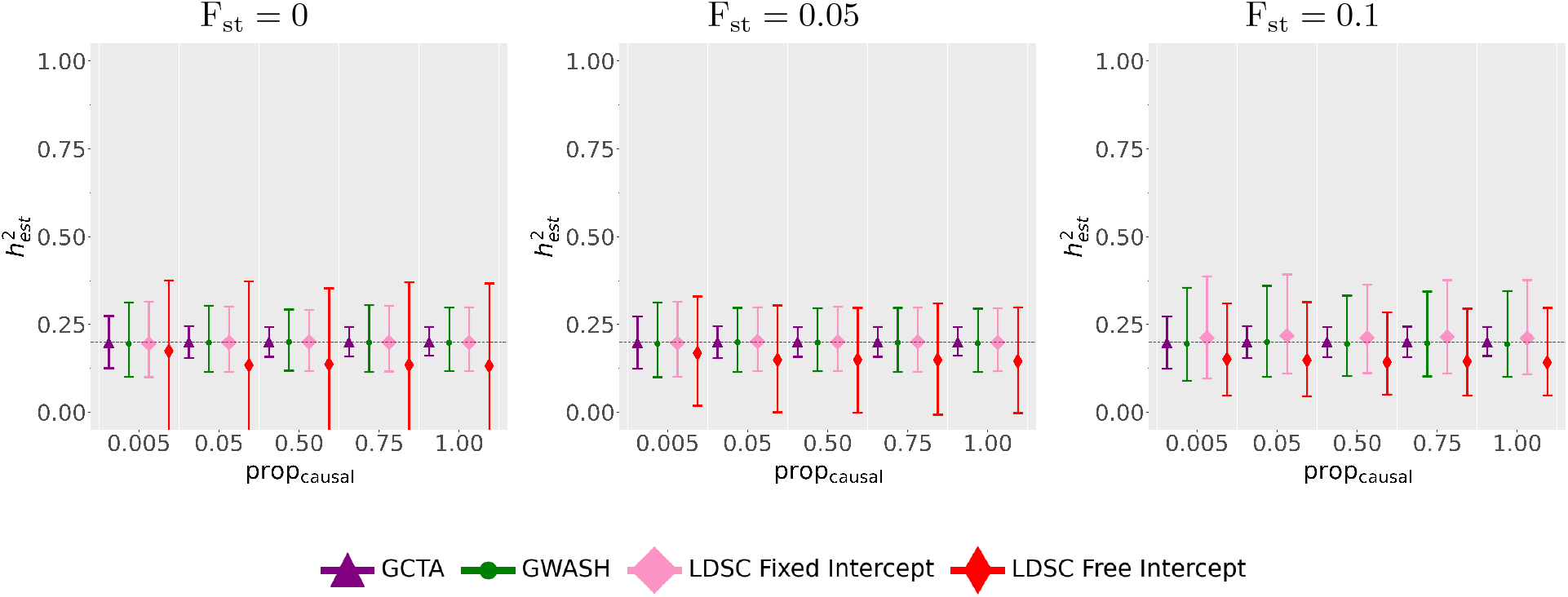
Heritability estimates on simulated data across 1000 simulations where a parameter of interest is gradually increased from F_st_ = 0 (left panel) to F_st_ = 0.05 (middle panel) and F_st_ = 0.1 (right panel). The setting is the same as in Figure 1.

#### S3.2 Realistic prop_causal_ Simulations with reference panel

Like with the AR1 simulations, we observe that prop_causal_ has no effect on heritability estimation.

**Figure S5:**
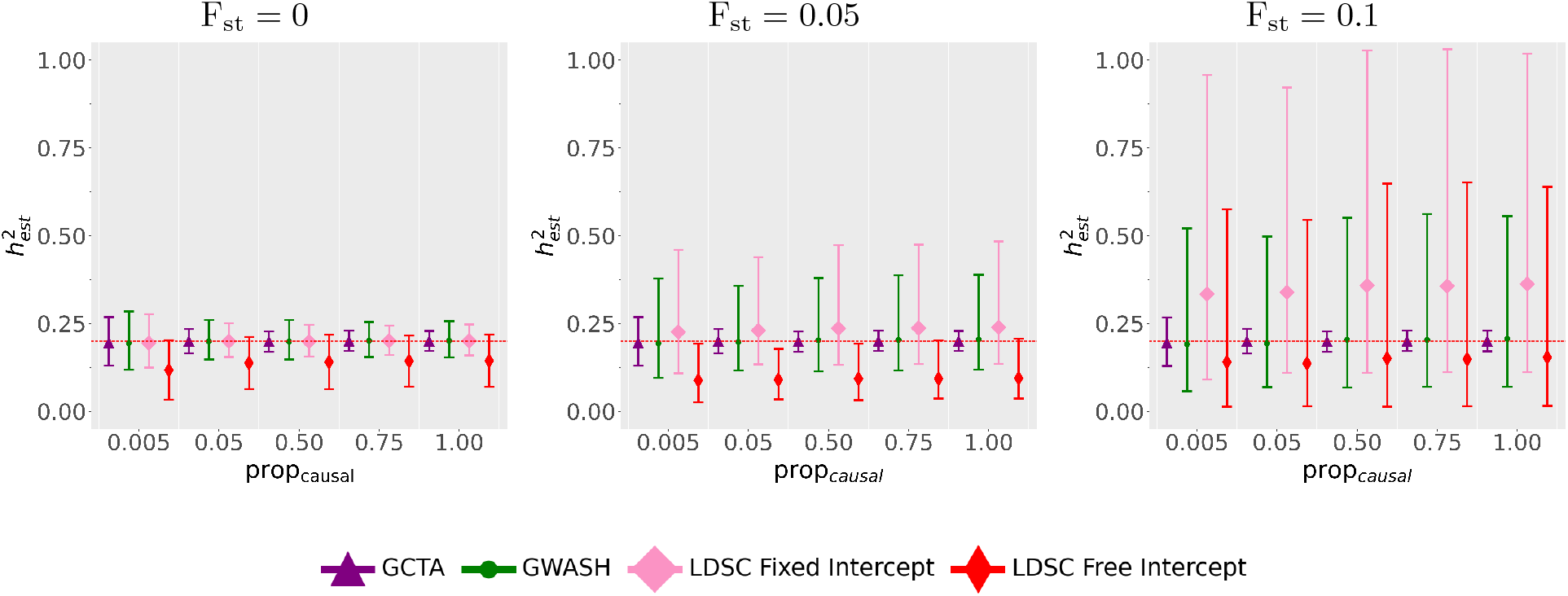
Heritability estimates on simulated data across 1000 simulations where a parameter of interest is gradually increased from F_st_ = 0 (left panel) to F_st_ = 0.05 (middle panel) and F_st_ = 0.1 (right panel). The setting is the same as in Figure 2.

#### S3.3 AR1 prop_causal_ Standard Error Estimation

**Figure S6:**
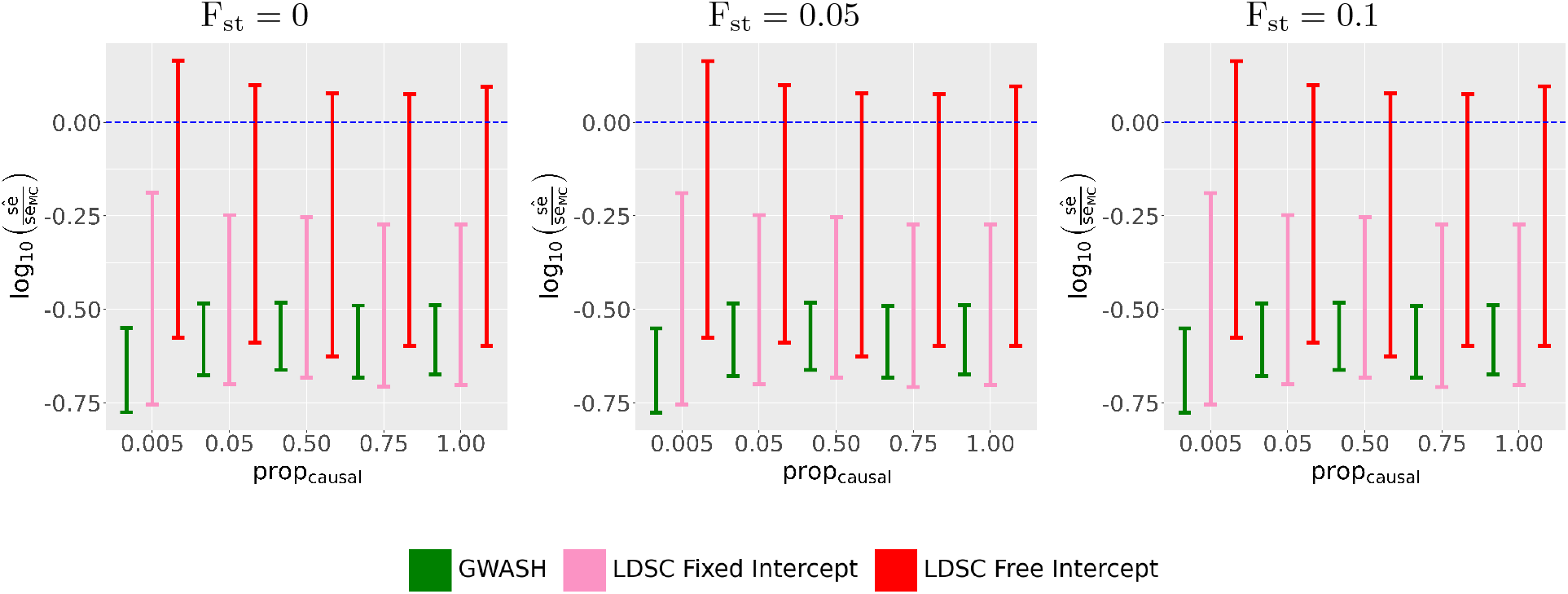
Heritability estimates on simulated data across 1000 simulations where a parameter of interest is gradually increased from F_st_ = 0 (left panel) to F_st_ = 0.05 (middle panel) and F_st_ = 0.1 (right panel). The setting is the same as in Figure 3.

#### S3.4 Realistic prop_causal_ Standard Error Estimation

**Figure S7:**
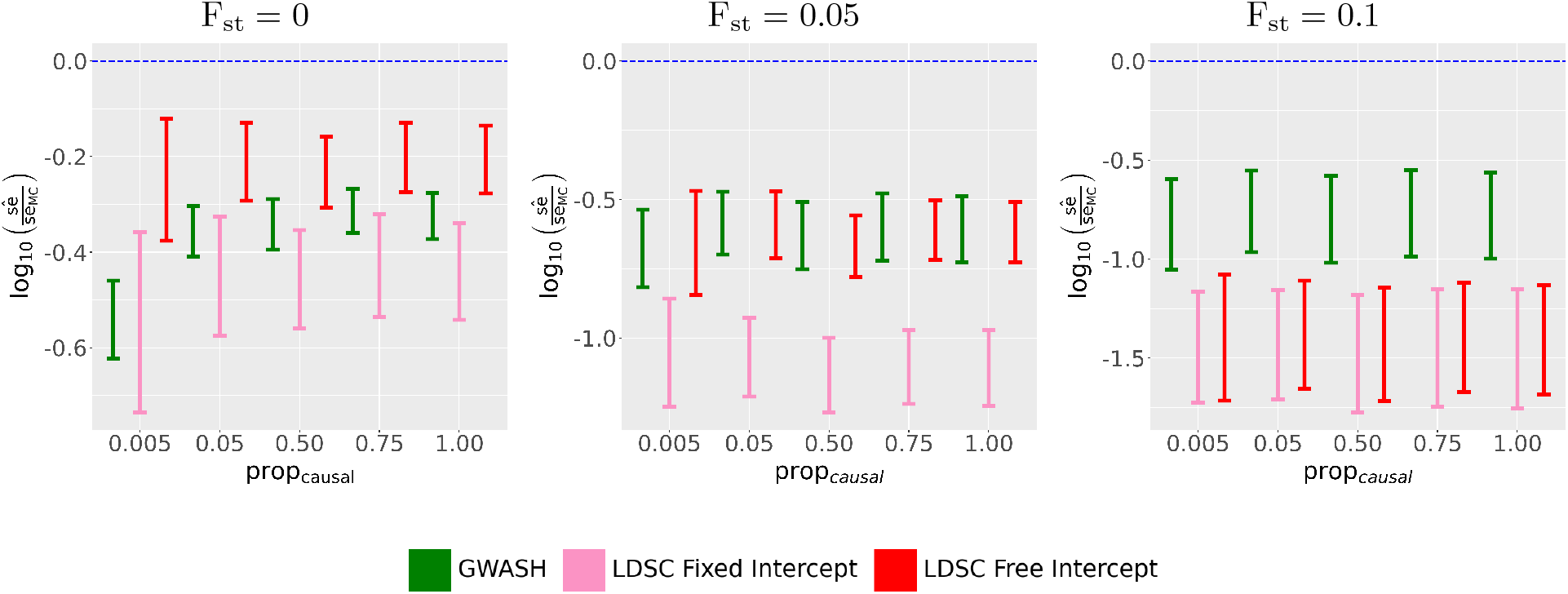
Heritability estimates on simulated data across 1000 simulations where a parameter of interest is gradually increased from F_st_ = 0 (left panel) to F_st_ = 0.05 (middle panel) and F_st_ = 0.1 (right panel). The setting is the same as in Figure 4.

#### S3.5 Impact of standard error estimates on Z-Scores in AR1 Simulations when changing prop_causal_

**Figure S8:**
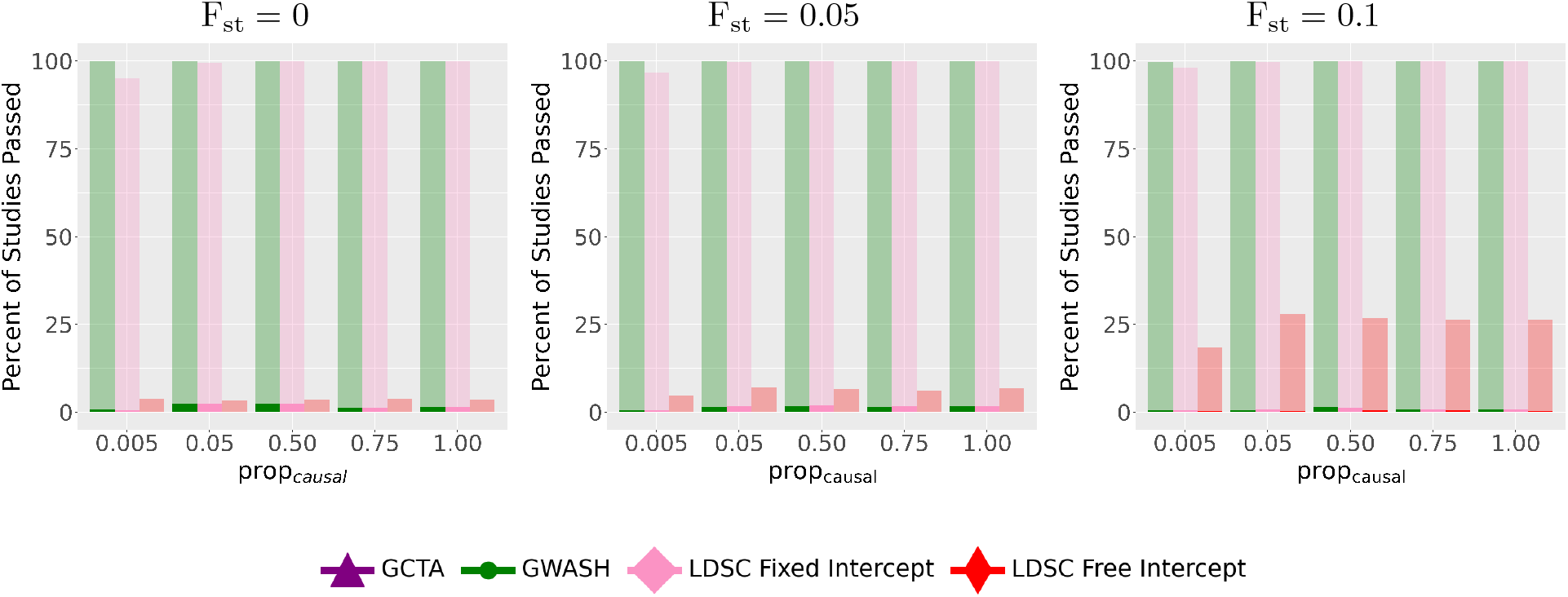
Heritability estimates on simulated data across 1000 simulations where a parameter of interest is gradually increased from F_st_ = 0 (left panel) to F_st_ = 0.05 (middle panel) and F_st_ = 0.1 (right panel). The setting is the same to that in Figure 1.

#### S3.6 Impact of standard error estimates on Z-Scores in realistic Simulations when changing prop_causal_

**Figure S9:**
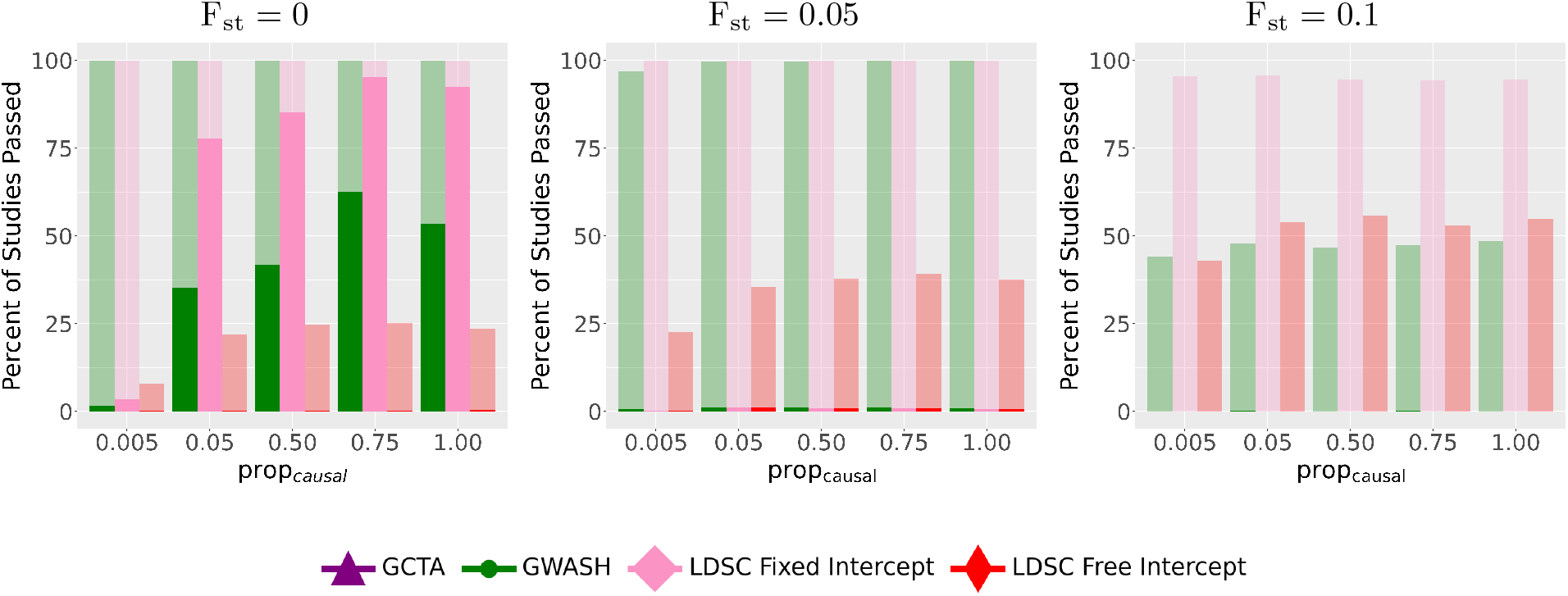
Heritability estimates on simulated data across 1000 simulations where a parameter of interest is gradually increased from F_st_ = 0 (left panel) to F_st_ = 0.05 (middle panel) and F_st_ = 0.1 (right panel). The setting is the same to that in Figure 2.

### S4 Simulations with Individual-Level Data

#### S4.1 AR1 Supplementary Simulations with Individual Level Data

These AR1 simulations have the same setting as those in the main text, using the same reference panel as the individual-level data. This is the most optimistic case where the reference panel matches exactly the individual level data.

**Figure S10:**
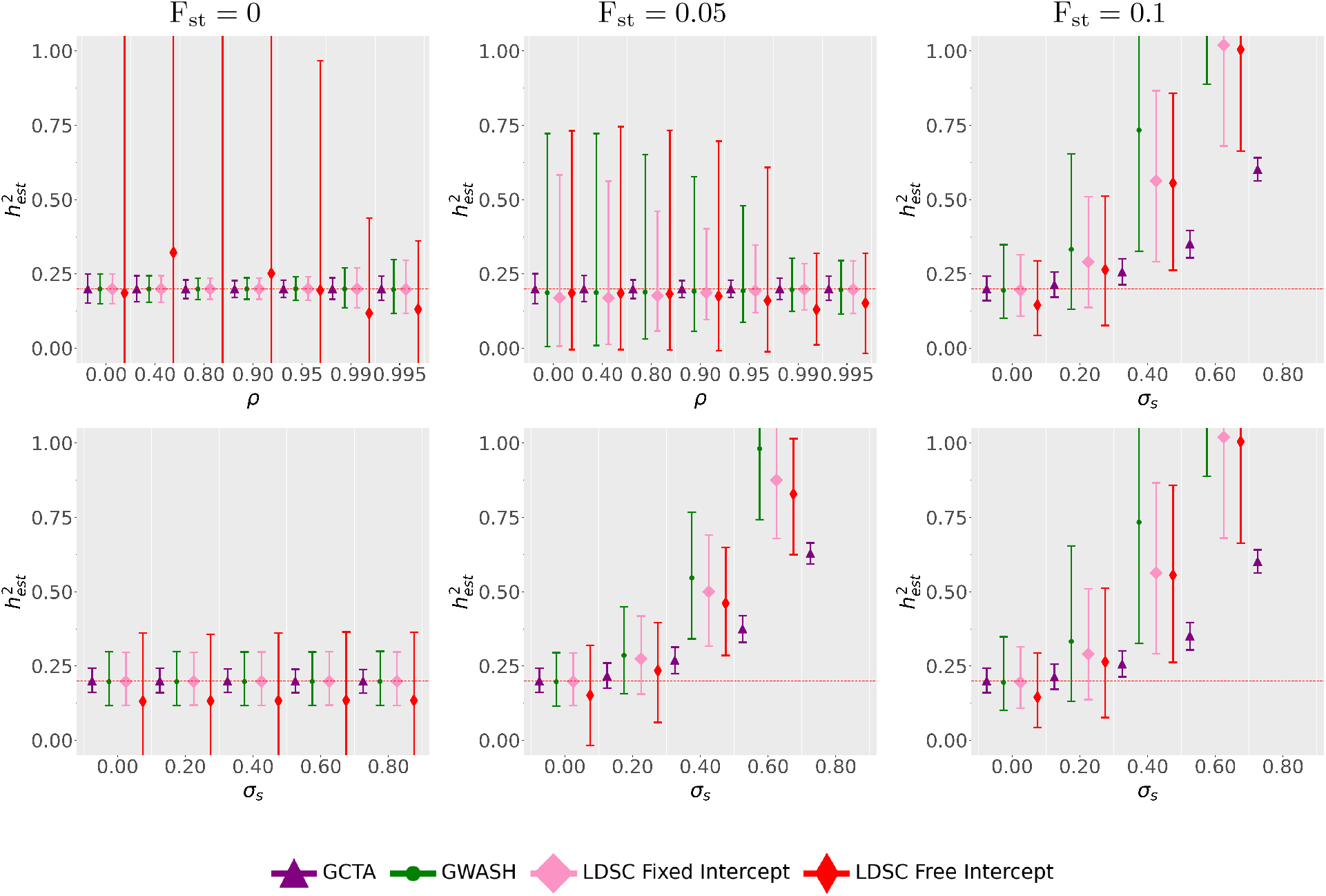
Heritability estimates when changing *ρ* and *σ*_*s*_ while gradually increasing F_st_ from F_st_ = 0 (left column) to F_st_ = 0.05 (middle column) and F_st_ = 0.1 (right column). The setting is similar to that in Figure 1 except that the sample data is observed and can thus be used as the reference panel.

#### S4.2 Realistic Supplementary Simulations with Individual Level Data

These Realistic simulations have the same setting as those in the main text, using the same reference panel as the individual-level data. This is the most optimistic case where the reference panel matches exactly the individual level data.

**Figure S11:**
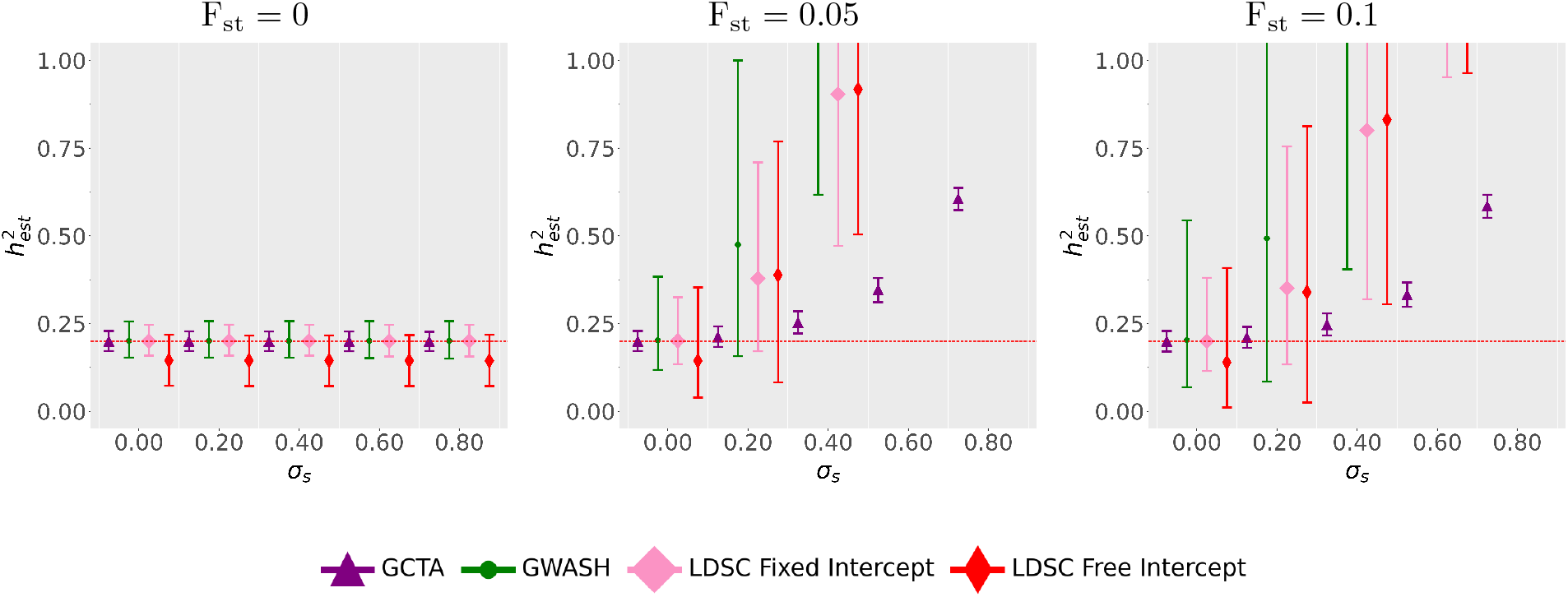
Heritability estimates on simulated data across 1000 simulations where a parameter of interest is gradually increased from F_st_ = 0 (left panel) to F_st_ = 0.05 (middle panel) and F_st_ = 0.1 (right panel). The setting is similar to that in Figure 2 except that the sample data is observed and can thus be used as the reference panel.

### S5 Estimation of Standard Error

#### S5.1 Standard Error AR1 Supplementary Simulations with Individual-Level Data

These Standard Error evaluations correspond to the AR1 Supplementary Figures with full individual-level data.

**Figure S12:**
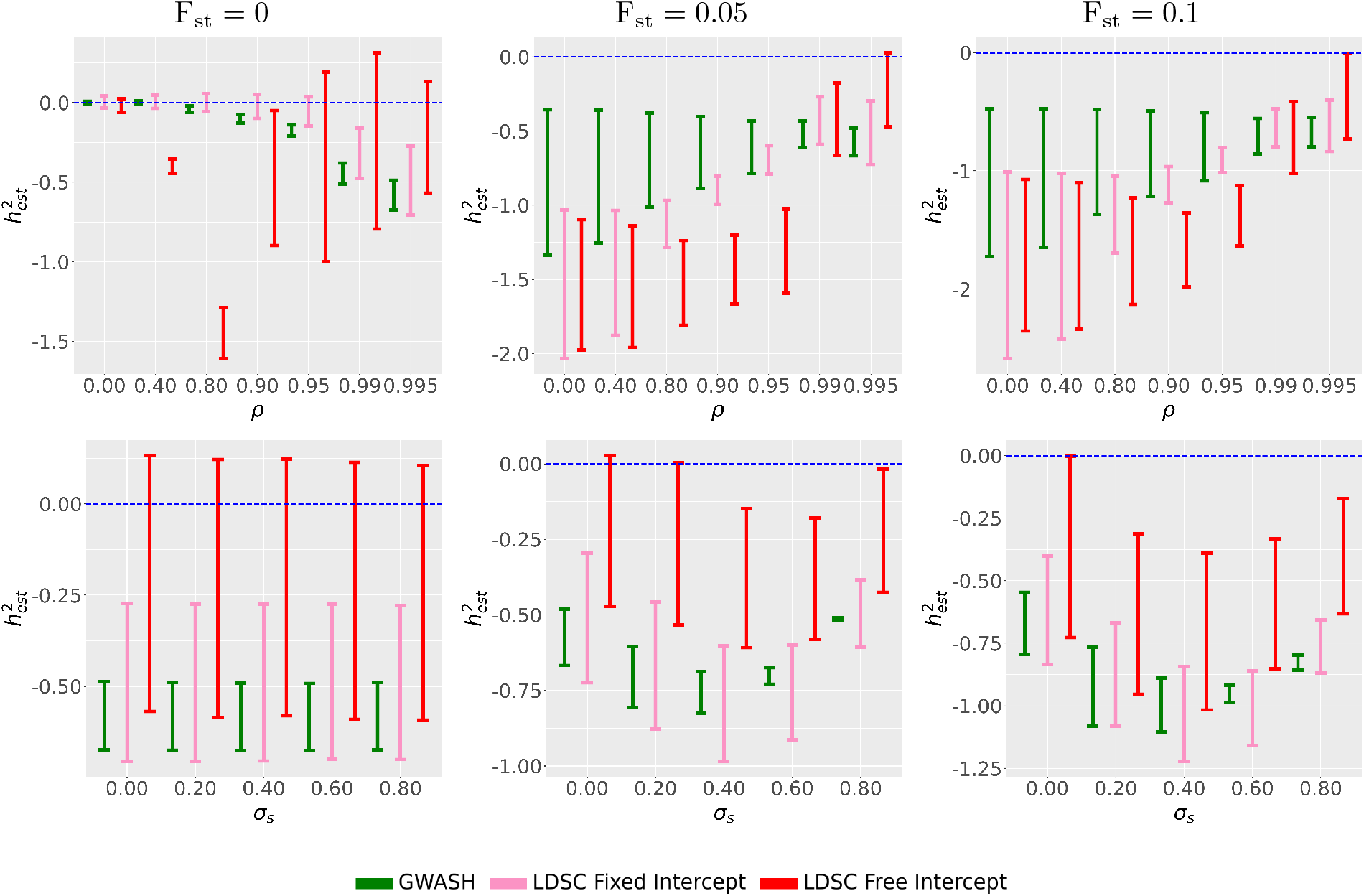
Evaluation of sample standard errors in AR1 simulated data with genetic confounding across 1000 replicates. The setting is the same as in Figure 1 except that the sample data is observed and is used as the reference panel.

#### S5.2 Standard Error Realistic Supplementary Simulations with Individual-Level Data

These Standard Error evaluations correspond to the Realistic Supplementary Figures with full individual-level data.

**Figure S13:**
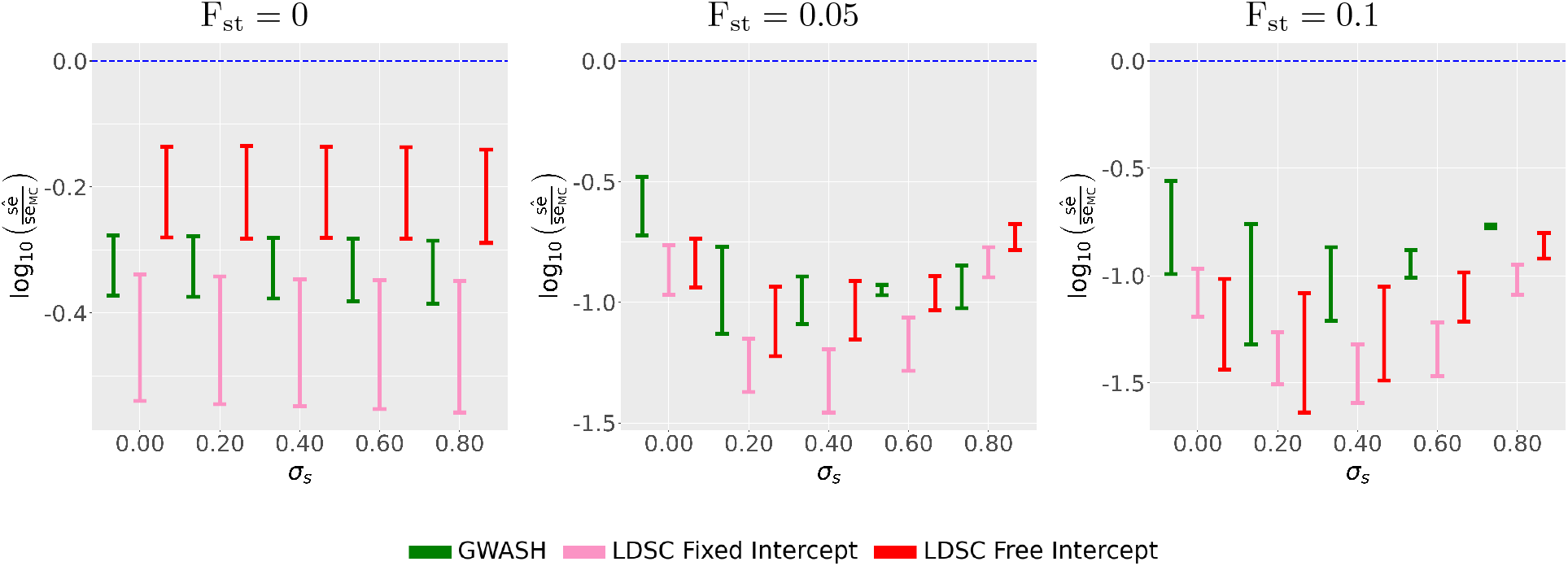
Evaluation of sample standard errors in LD simulated data with genetic confounding across 1000 replicates. The setting is the same as in Figure 4 except that the sample data is observed and is used as the reference panel.

#### S5.3 Impact of standard error estimates on Z-Scores in Individual-Level AR1 Simulations

**Figure S14:**
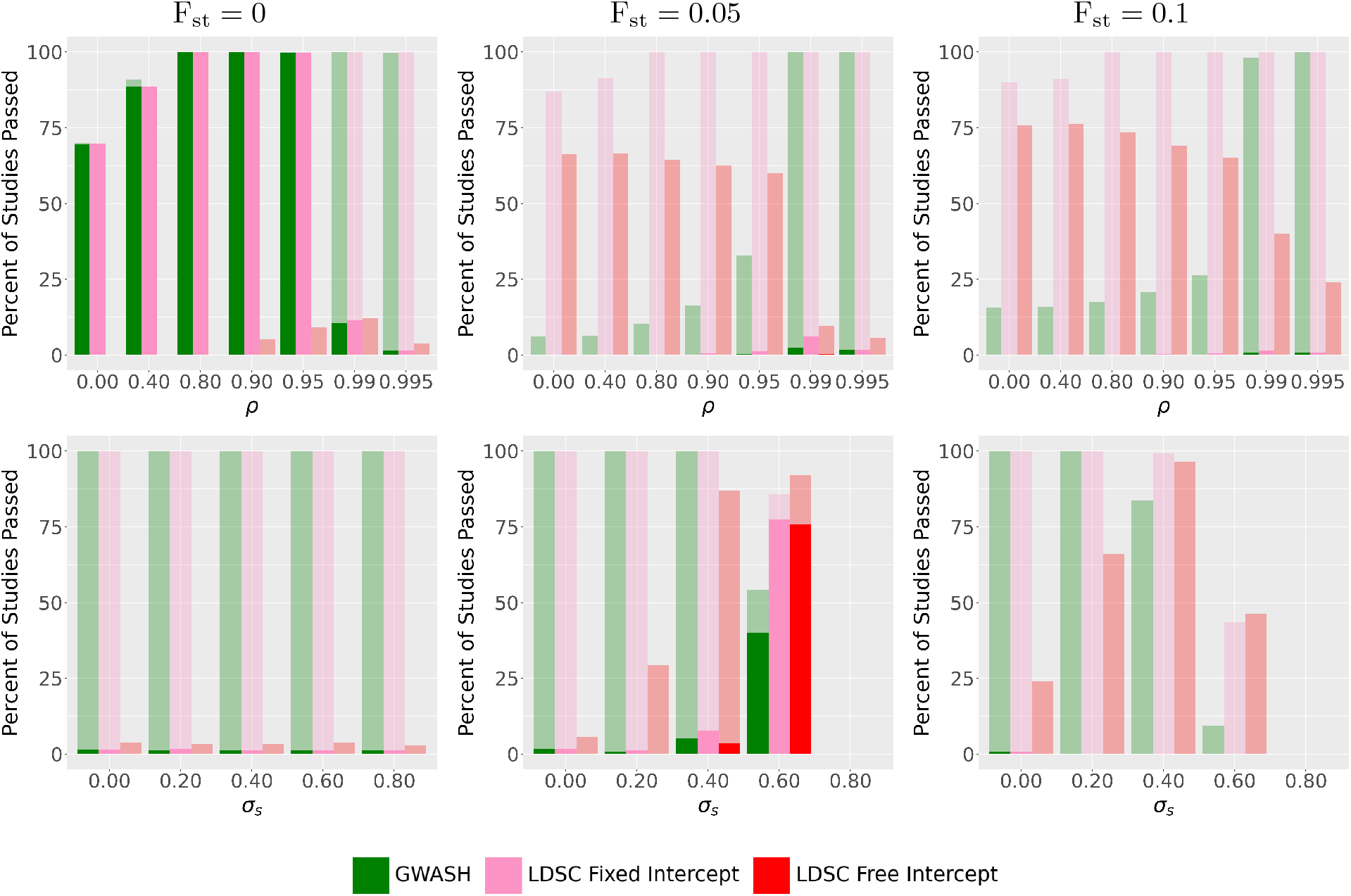
Evaluation of studies passed based on heritability z-scores in AR1 Simulations changing *ρ* or *σ*_*s*_ after F_st_ is increased. The left column shows no genetic confounding (F_st_ = 0.0), the middle column is with moderate genetic confounding (F_st_ = 0.05) and the right column is with high genetic confounding (F_st_ = 0.1). This figure is the same as Figure 5 except that the sample data is observed and is used as the reference panel.

#### S5.4 Impact of standard error estimates on Z-Scores in Individual-Level Realistic Simulations

**Figure S15:**
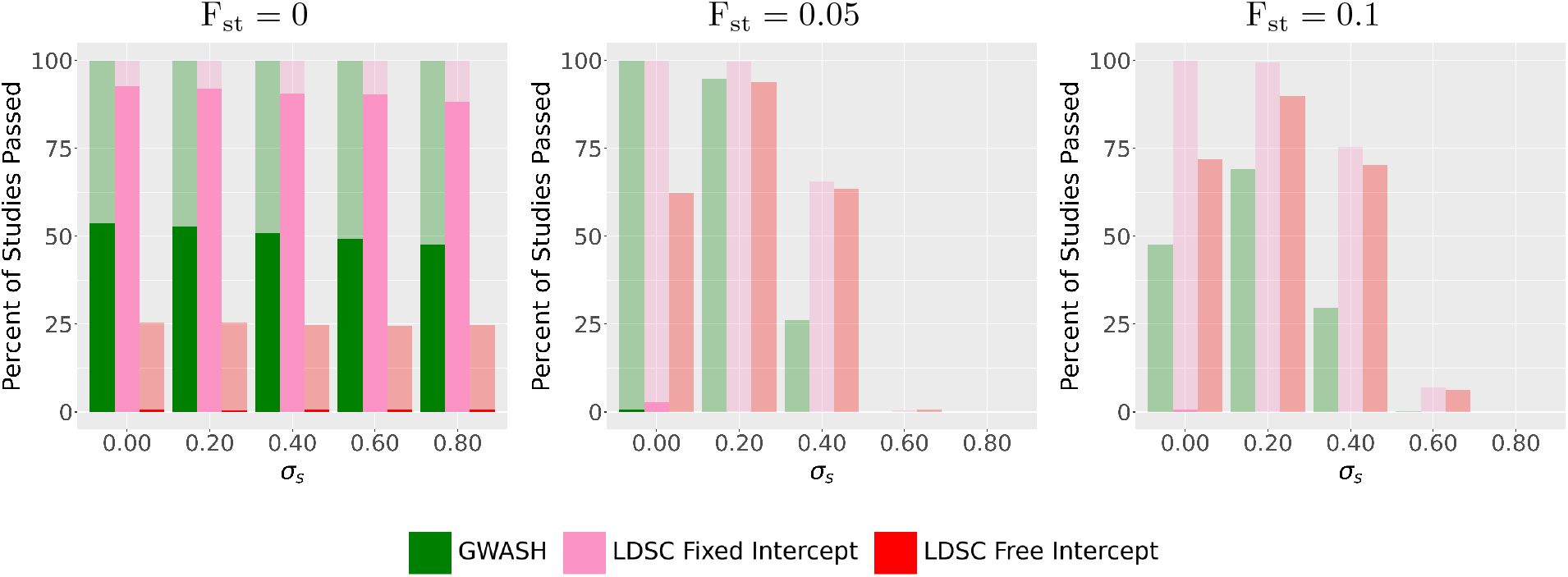
Evaluation of studies passed based on heritability z-scores in Realistic Simulations changing a parameter after F_st_ is increased from F_st_ = 0 (left panel) to F_st_ = 0.05(middle panel) and F_st_ = 0.1 (right panel). This is the same setting as in Figure 6 except that the sample data is observed and is used as the reference panel.

